# Nanoscopic resolution within a single imaging frame

**DOI:** 10.1101/2021.10.17.464398

**Authors:** Esley Torres García, Raúl Pinto Cámara, Alejandro Linares, Damián Martínez, Víctor Abonza, Eduardo Brito-Alarcón, Carlos Calcines-Cruz, Gustavo Valdés Galindo, David Torres, Martina Jabloñski, Héctor H. Torres-Martínez, José L. Martínez, Haydee O. Hernández, José P. Ocelotl-Oviedo, Yasel Garcés, Marco Barchi, Rocco D’Antuono, Ana Boskovic, Joseph G. Dubrovsky, Alberto Darszon, Mariano G. Buffone, Roberto Rodríguez Morales, Juan Manuel Rendon-Mancha, Christopher D. Wood, Armando Hernández-García, Diego Krapf, Álvaro H. Crevenna, Adán Guerrero

## Abstract

Mean-Shift Super Resolution (MSSR) is a principle based on the Mean Shift theory that extends spatial resolution in fluorescence images, beyond the diffraction limit. MSSR works on low- and high-density fluorophore images, is not limited by the architecture of the detector (EM-CCD, sCMOS, or photomultiplier-based laser scanning systems) and is applicable to single images as well as temporal series. The theoretical limit of spatial resolution, based on optimized real-world imaging conditions and analysis of temporal image series, has been measured to be 40 nm. Furthermore, MSSR has denoising capabilities that outperform other analytical super resolution image approaches. Altogether, MSSR is a powerful, flexible, and generic tool for multidimensional and live cell imaging applications.

## Introduction

Super-resolution Microscopy (SRM), which encompasses a collection of methods that circumvent Abbe’s optical resolution limit, has dramatically increased our capability to visualize the architecture of cells and tissues at the molecular level. There are several approaches to SRM which vary in terms of the final attainable spatial and temporal resolution, photon efficiency, as well as in their capacity to image live or fixed samples at depth [1, 2]. One class of techniques exceed the diffraction limit by engineering the illumination or the point spread function (PSF), such as SIM and STED [3–5]. These techniques can be used for live imaging although they require specialized hardware and dedicated personnel for maintenance and operation. Single-molecule localization methods (e.g., STORM, PAINT, PALM) [6–9] that localize individual emitters with nanometer precision require temporal analysis of several hundred-to-thousands of images and are prone to error due to fast molecular dynamics within live specimens.

Some SRM computational methods have few or no demands on hardware or sample preparation and provide resolution improvements beyond the diffraction limit, i.e., fluorescence fluctuation-based super resolution microscopy (FF-SRM) approaches [10–13]. The quantity and performance of computational methods have both increased over the past decade given the many advantages they present, such as their low barriers to entry and generic applicability to data acquired with any microscopy modality (wide-field, confocal, or light-sheet). However, these methods also present some limitations, such as the possible introduction of artifacts [14], the requirement for high signal-to-noise ratio (SNR) data and the acquisition of tens to hundreds of frames [10–13], which limit their applicability to reconstruct fast dynamical processes.

The problem of spatial resolution in optical microscopy can be addressed from the statistical point of view. In the case of fluorescence microscopy, the process of photon emission from punctual sources (fluorescence emitters) can be considered as a discrete distribution of information, where the unitary element of the distribution is the photon [15]. In this scenario, the problem of spatial resolution gets reduced to the problem of finding modes of information, regardless of the shape of the distribution, hence, disconnecting the problem of optical resolution from the diffraction boundary [16].

Here, we introduce the Mean Shift Super-Resolution principle for digital images ‘MSSR’ (pronounced as *messer*), derived from the Mean Shift (MS) theory [17, 18].

MSSR extends the resolution of any single fluorescence image up to 1.6 times, including its use as a resolution enhancement complement after the application of other super-resolution methods.

By computing the local magnitude of the Mean Shift vector, MSSR generates a probability distribution of fluorescence estimates whose local magnitude peaks at the source of information. As a result of that, the spatial distribution becomes ‘refined’ (i.e., for a Gaussian distribution of fluorescence its width shrinks). Additionally, we demonstrate the extended-, enhanced- and super-resolving capabilities of MSSR as a standalone method for a variety of fluorescence microscopy applications, through a single-frame and temporal stack analysis, allowing resolution improvements towards a limit of 40 nm.

Open-source implementations of MSSR are provided for ImageJ (as a plugin), R, and MATLAB, some of which take advantage of the parallel computing capabilities of regular desktop computers (Supplementary Note 7). The method operates almost free of parameters; users only need to provide an estimate of the point spread function (PSF, in pixels) of the optical system, choose the MSSR order, and decide whether a temporal analysis will take place (Supplementary Material and MSSR Manual). The provided open-source implementations of MSSR represent a novel user-friendly alternative for the bioimaging community for unveiling life at its nanoscopic level.

## Results

### The MSSR principle

MSSR is tailored around the assumption that fluorescence images are formed by discrete signals collected (photons) from point sources (fluorophores) convolved with the PSF of the microscope (Supplementary Notes 1, 2 and 3). Processing a single image with MSSR starts with the calculation of the MS, which guarantees that large intensity values on the diffraction-limited (DL) image coincide with large positive values in the MSSR image (Supplementary Note 4). Further algebraic transformations then restore the raw intensity distribution and remove possible artifacts caused by the previous step (edge effects and noise dependent artifacts), giving rise to an image that contains centers of density with a narrower full width at half maximum (FWHM) (Figure 1a). This procedure is denoted by MSSR of order zero (MSSR^0^), and it is the first stage which shrinks emitter distribution.

**Figure 1.**
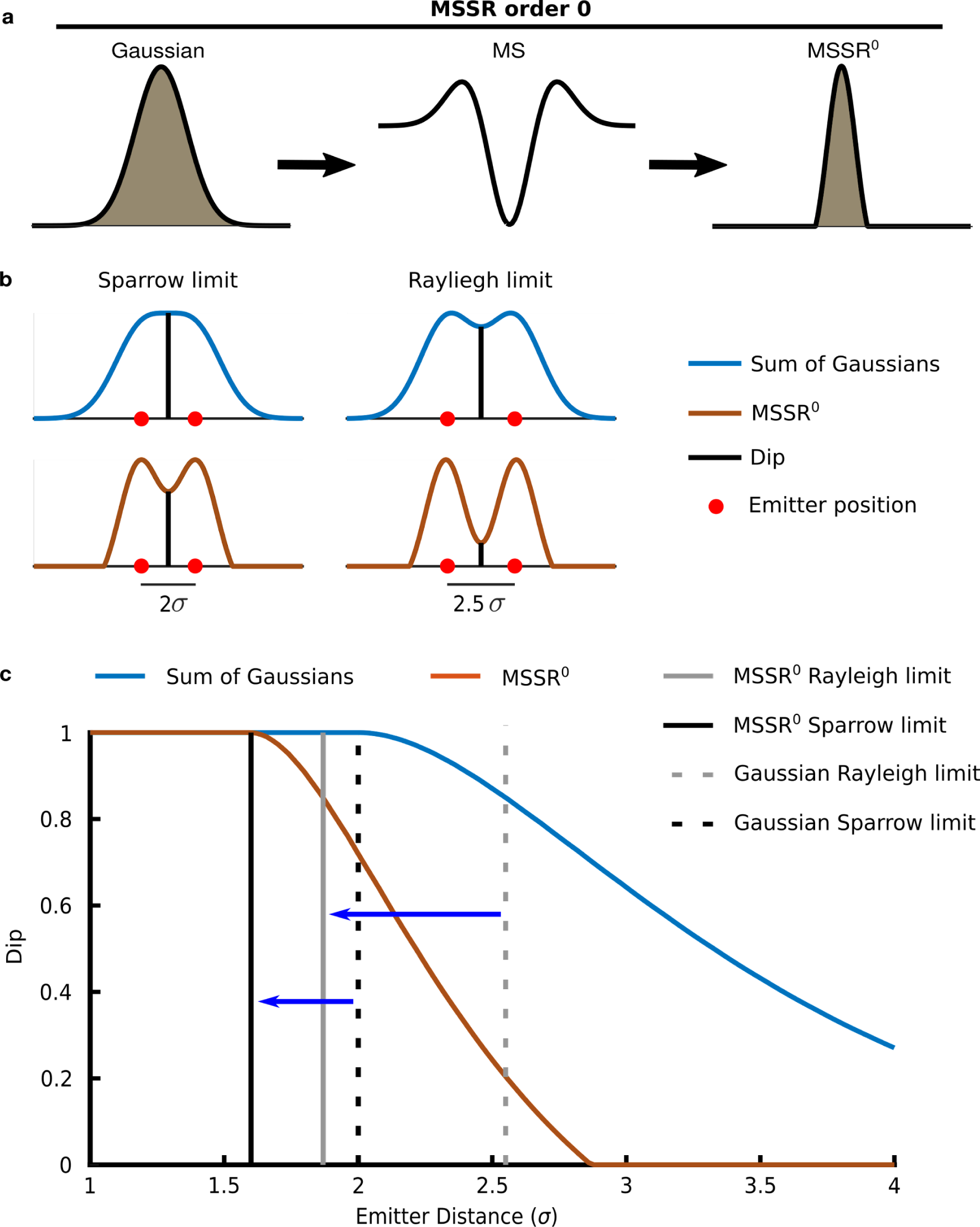
MSSR of zero order increases resolution by reducing the width of the spatial distribution of photons emitted from modeled fluorescent emitters. a) The MS is applied to the initial Gaussian distribution of photons emitted by a point-source (left) resulting in a MS graph (center). Application of further algebraic transformations (see Supplementary Note 5 and Figure S11 (ii-iv)) provides the MSSR^0^ distribution (right). b) Sparrow and Rayleigh limits (blue, diffraction-limited) and the corresponding MSSR^0^ transformation (brown) for two point-sources. Red dots represent each emitter’s location. The dip is indicated by a vertical black line. The inter-emitter distance is expressed as σ-times their individual standard deviation before MSSR processing. c) Dip computed for two point-source emitters of Gaussian distribution located away at distinct σ (blue line) where the corresponding MSSR^0^ result is also depicted (red line). For Gaussian: Rayleigh limit – gray discontinuous line, Sparrow limit - black discontinuous line. For MSSR^0^: Rayleigh limit - gray solid line, Sparrow limit - gray solid line. The solid vertical lines represent the distance between emitters such that when processed with MSSR^0^, the criterions of Rayleigh and Sparrow are obtained (For detail see Online Methods section *Simulation of fluorescent emitters*).

The MS is locally computed by a kernel window that slides throughout the entire image, subtracts the sample mean (weighted local mean) as well as the central value of the kernel using a spatial-range neighborhood (Supplementary Notes 2 and 3, Supplementary Figure S5, Supplementary Table S1) [17, 18]. The MS is a vector that always points towards the direction of the intensity gradient and its length provides a local measure of the fluorescence density and brightness [19–21]; its magnitude depends on the value difference between the central pixel of the neighborhood and the surrounding pixels. A mathematical proof, provided in Supplementary Note 4, demonstrates that the minimum MS value, computed from a Gaussian distribution, matches with the point of maximum intensity of the initial distribution (Supplementary Note 4, Supplementary Figure S6).

The increase in resolution offered by MSSR^0^ was evaluated by the Rayleigh and Sparrow limits [22–24], which are two criteria that establish resolution bounds for two near-point sources (Figure 1b). Processing with MSSR^0^ of two-point sources located at their resolution limit (2.5 σ and 2 σ for Rayleigh and Sparrow limit respectively, Figure 1c vertical discontinuous lines) decreases the dip (height at the middle point) [25] within their intensity distributions (Figure 1b and 1c). Processing a single image with

MSSR^0^ shifts the resolution limit by 26 % and 20 % according to the Rayleigh and Sparrow limits, respectively, and reduces the FWHM of individual emitters (Figure 1c vertical continuous lines). A comparison of the shrinkability of MSSR ^0^ applied to Gaussian and Bessel distributions are shown in Supplementary Figure S9. The reduction of FWHM of Bessel distribution at different wavelengths of the visible spectrum are shown in Supplementary Figure S10.

Since the result of MSSR^0^ is an image, we used the resulting image to seed an iterative process (Figure 2a). We refer to this as higher-order MSSR (MSSR^n^, with n>0), which delivers a further gain of resolution per *n*-iteration step (Figure 2a and Supplementary Figure S11). As the order of MSSR^n^ increases, both the FWHM of emitters (Supplementary Figure S12) and the dip of their intensity distribution decrease (Figure 2b). Numerical approximations indicate that two point-sources separated at 1.6 σ are resolvable with MSSR^3^, but not when their separation is 1.5 σ (Figure 2b). The separation of 1.6 σ sets the theoretical resolution limit of MSSR^n^.

**Figure 2.**
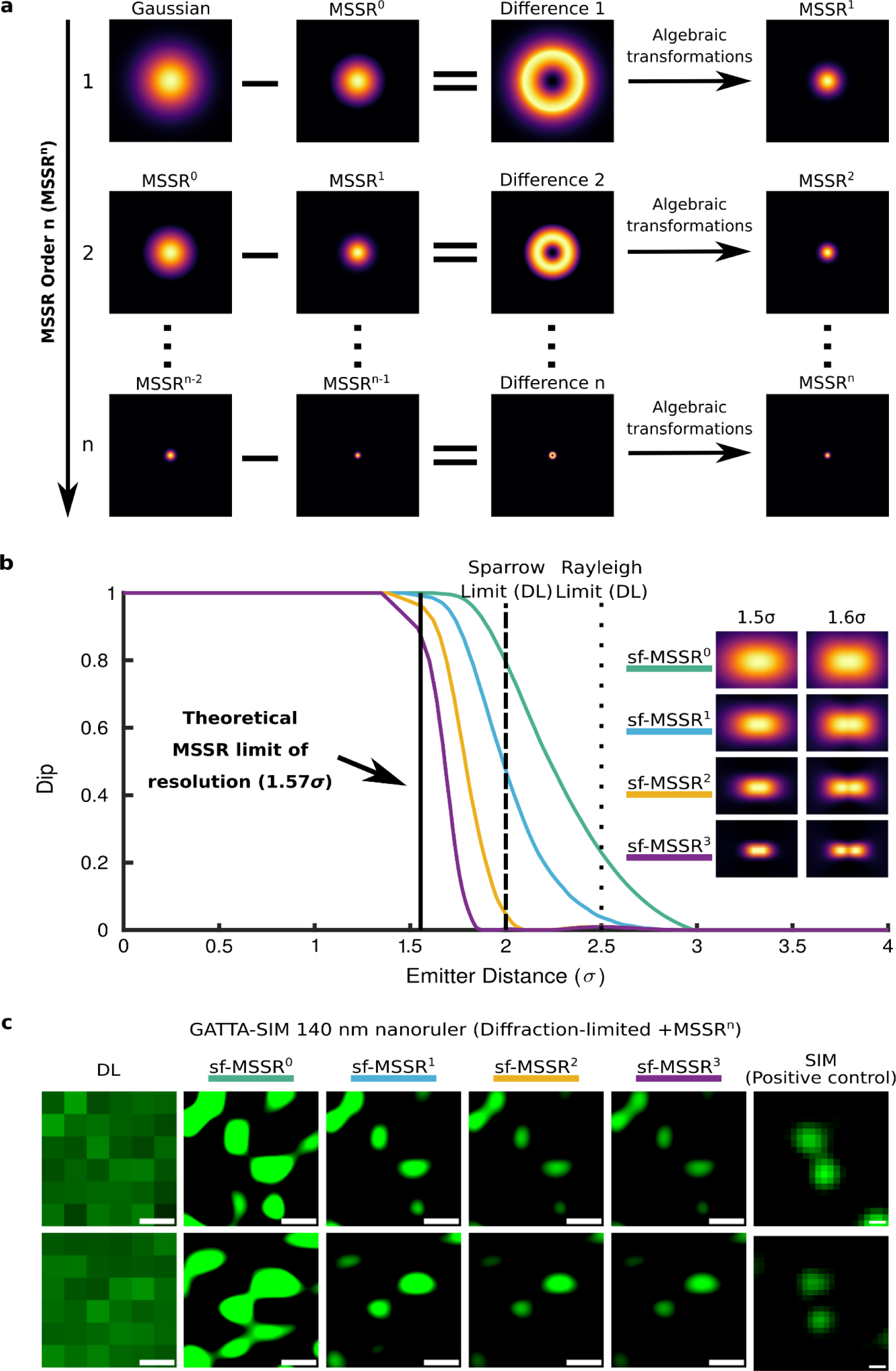
Single-frame MSSR analysis of higher order attains a resolution limit of 1.6 σ for nearby emitters. **a)** The algorithm for computing higher-order MSSR (MSSR^n^) is presented. The first iteration of MSSR (MSSR^1^) is given by subtracting the MSSR^0^ from the original image, resulting in a donut-like region centered at the emitter’s location. MSSR^1^ is computed after applying further algebraic transformations (see Supplementary Note 5 and Figure S11 (ii-iv) for a full description of the MSSR^n^ process). The second iteration encompasses the subtraction of MSSR^1^ from MSSR^0^ and the same algebraic transformations as used for generation of MSSR^1^. The process is repeated by updating consecutive MSSR images which generates higher MSSR orders. **b)** Theoretical limit of resolution achievable by MSSR^n^. Dip computed for two Gaussian emitters in accordance with the variation of the inter-emitter distance (expressed as σ-times their standard deviation before MSSR processing). Colored lines represent the dip of MSSR order, from 0 to 3, computed at a given σ distance between emitters. Images on the right are the bidimensional representation of the MSSR^n^ processing for two single emitters separated at distances of 1.5σ and 1.6σ. Note that, for 1.5σ, emitters are unresolved up to the third order of MSSR (For detail see Online Methods section *Simulation of fluorescent emitters*). Dot and dashed lines are referred to Rayleigh and Sparrow limits for the DL case and the continuous line marks the MSSR limit of resolution. **c)** Experimental demonstration of the resolution increases attainable with higher order MSSR using the GATTA-SIM 140B nanoruler system. The intensity distribution of the emitter shrinks, both in σ and intensity, as the order of the MSSR increases (Figure S12). Nearby emitters (Alexa Fluor® 488) located 140 nm apart are resolved using MSSR^1^, MSSR^2^ and MSSR^3^ (right side). SIM images collected from the same sample (distinct fields) are shown as a positive control. sf-MSSR parameters: AMP = 10, FWHM of PSF = 3.48, order = 0-3. ImageDecorrelation parameters for DL case: Rmin=0, Rmax = 0.7, Nr = 50, Ng = 10. ImageDecorrelation parameters for sf-MSSR^n^ (n=0-3): Rmin=0, Rmax = 0.3, Nr = 50, Ng = 10. Scale bar: 100 nm.

In summary, MSSR^n^ processing extends the spatial resolution of single DL images. The procedure of applying MSSR^n^ to a single DL image will be defined as sf-MSSR^n^.

### MSSR is a deconvolution approach at the nano scales

In optical microscopy, objects significantly closer to the diffraction boundary can be resolved with clever illumination and detection schemes (i.e., SIM, Airy Scan, 4 Pi, I5M, STED, SMLM, etc.) [3, 26–28], or by careful image analysis, reviewed in [29]. Rayleigh criterion is a bit too conservative, in the sense that achieving a decrease of the Dip formed by the joint distribution shaped by two adjacent emitters might be interpreted as surpassing the diffraction boundary (Supplementary Figure S13 a, b).

Repeating the same procedure using a joint distribution shaped by two adjacent emitters located at the Sparrow limit there is no further gain of resolution, as the dip remains constant, taking the value of 1 (Supplementary Figure S13 e, f).

MSSR aims to revert the effect of diffraction on optical microscopy, so it can be considered as a deconvolution process. As the diffraction can be modeled with a Gaussian spread, the pixel value is a superimposition of spreads from individual emitters (Supplementary Notes 1, 2 and 3). The goal is to reduce the spread. The latter can be accomplished by “sharpening by blur” [30, 31]. In MSSR, the computation of the MS is the blurring process used to sharpen the image. What makes sf-MSSR^n^ unique is the fact that it extends spatial information down below the Sparrow limit. Processing the joint distributions of Supplementary Figure S13 with sf-MSSR^n^ of any order n leads to a decrease of the dip value (Supplementary Figure S13 c, d, g, f). Furthermore, sf-MSSR^3^ processing collapses the dip to zero for both Rayleigh and Sparrow conditions (Supplementary Figure S13 d, h).

To illustrate how MSSR works by sharpening features down the diffraction barrier, we provide comparative data against: Wiener deconvolution [32], Richardson-Lucy deconvolution [33, 34] and the Radiality Maps (RMs) [11].

Supplementary Figure S14 shows that Wiener deconvolution partially restores the effect of diffraction, but without a dramatic increase in spatial resolution. Interestingly, Richardson-Lucy deconvolution provides a noticeable increase in resolution at the boundaries of the Rayleigh limit but fails to extend spatial resolution down below the Sparrow limit (Supplementary Figure S14).

Gustafsson et al. showed that the RMs of SRRF provide a resolution increase down to 0.7 times the Gaussian FWHM [11], where when the peak separation between isolated is greater than 0.7 times the FWHM of the PSF, they can be directly resolved without further enhancement provided by higher-order statistical analysis. MSSR^0^ and the Radiality Maps are similar in the sense that both perform sharpening and smoothing. Supplementary Figure S14a shows that both MSSR and the RMs overcome the Rayleigh diffraction limit [22]. However, the RMs produce undesired artifacts which are absent when using sf-MSSR^0^ (Supplementary Figure S14b), which is in agreement with the reported spatial artifacts introduced [35]. Figures 1, 2 and Supplementary Figure S14 show that sf-MSSR^0^ reliably provides spatial resolution gains, free of image analysis artifacts, down to the Rayleigh limit, hence, allowing the study of nanoscopic regimes at the boundaries of the Sparrow limit.

With such data in hand, it is concluded that sf-MSSR^n^ is a deconvolution process that extends spatial information of DL images at the nano scales.

### MSSR extends spatial resolution in fluorescence microscopy images

To empirically test the ability of MSSR to extend spatial resolution within a single DL image, a commercial nanoruler sample (GATTA-SIM140B, GATTAquant) was imaged by Structured Illumination Microscopy (SIM) and Total Internal Reflection Fluorescence (TIRF) microscopy, which was then processed by sf-MSSR^n^. The iterative processing of the widefield data with sf-MSSR^3^ reveals the two fluorescence emitters located at a separation of 140 nm, which is consistent with the result obtained by SIM (Figure 2c).

A theoretical approximation of the Rayleigh limit for a GATTA-SIM 140B nanoruler (Figure 2c of the main manuscript) with λ_em_= 525 nm and NA = 1.4, is d = (0.61*525)/1.4 = 229 nm (Table 1). Figures 1c and 2b show that MSSR processing theoretically extends spatial resolution. Computation of resolution on sf-MSSR^0^ using the Rayleigh criterion (at 0.74 of the Dip) gives a spatial resolution of 160 nm, which corresponds to a resolution change to 0.69 times (0.69x) the resolution limit. Using higher orders of sf-MSSR^n^ with n = 1, 2, 3, gives a resolution change of 0.66-0.64x the resolution limit (Table 1). Table 1 also shows the spatial resolution measured on the GATTA-SIM 140B nanorulers (experimental data), computed through decorrelation [36], using the Image Decorrelation plug-in for Fiji/ImageJ. Decorrelation computes the maximal observable frequency in an image (K_0_) as a proxy of spatial resolution (Resolution = 1/ K_0_). Note that sf-MSSR^n^ dramatically reduces resolution as a function of the n-order.

**Table 1.** sf-MSSR extends spatial resolution on simulated and real experimental conditions. For simulated conditions the Rayleigh limit and Sparrow limit were computed by the simulation of two fluorescent emitters. The values of FWHM of PSF (values in nm units) have been computed by measuring directly on the PSF distribution. The values of PSF sigma (values in nm units) were computed by fitting a Gaussian distribution and reporting the corresponding sigma parameter. sf-MSSR parameters: AMP = 1, FWHM of PSF = 198 nm, order = 0-3. ImageDecorrelation parameters for DL case: Rmin=0, Rmax = 0.7, Nr = 50, Ng = 10. ImageDecorrelation parameters for sf-MSSR^n^ (n = 0-3): Rmin=0, Rmax = 0.3, Nr = 50, Ng = 10.

| GATTA-SIM 140B | Resolution | diffraction limited | sf-MSSR<sup>0</sup> | sf-MSSR<sup>1</sup> | sf-MSSR<sup>2</sup> | sf-MSSR<sup>3</sup> |
| --- | --- | --- | --- | --- | --- | --- |
| Simulation | Rayleigh limit | 229 nm (1x)<br>2.90 $\sigma$ | 160 nm (0.69x)<br>2.02 $\sigma$ | 152 nm (0.66x)<br>1.92 $\sigma$ | 148 nm (0.65x)<br>1.87 $\sigma$ | 146 nm (0.64x)<br>1.841 $\sigma$ |
| | PSF FWHM | 192 nm<br>2.35 $\sigma$ | 114 nm<br>1.10 $\sigma$ | 84 nm<br>0.74 $\sigma$ | 56 nm<br>0.49 $\sigma$ | 38 nm<br>0.34 $\sigma$ |
| | PSF sigma | 79 nm<br>1 $\sigma$ | 44 nm<br>0.63 $\sigma$ | 34 nm<br>0.43 $\sigma$ | 24 nm<br>0.28 $\sigma$ | 16 nm<br>0.20 $\sigma$ |
| Experiment | resolution | 260 nm (1x) | 58 nm (0.22x) | 33 nm (0.13x) | 21 nm (0.08x) | 13 nm (0.05x) |

To further test the attainable resolution by sf-MSSR^n^ we used the ArgoLight test slide, acquiring images of the pattern formed by gradually spaced lines of fluorescent molecules (Argo-SIM, pattern E). The distance between lines increases from 0 to 390 nm with a step change of 30 nm: 0 nm, 30 nm, 60 nm, etc. Figure 3 shows the application of sf-MSSR^n^ to the confocal microscopy images of the Argo-SIM micropattern. As expected, the confocal acquisition allows to resolve parallel rows of fluorophores located at 240 nm. Remarkably, sf-MSSR^0^ processing extended spatial resolution down to 0.5 times the confocal resolution, allowing to discriminate parallel rows of fluorophores located at 120 nm. It is worth mentioning that this improvement comes at zero hardware cost, compared to other methods requiring specific optics/detectors such as the Airyscan [37] and Re-scan confocal microscopy [38].

**Figure 3.**
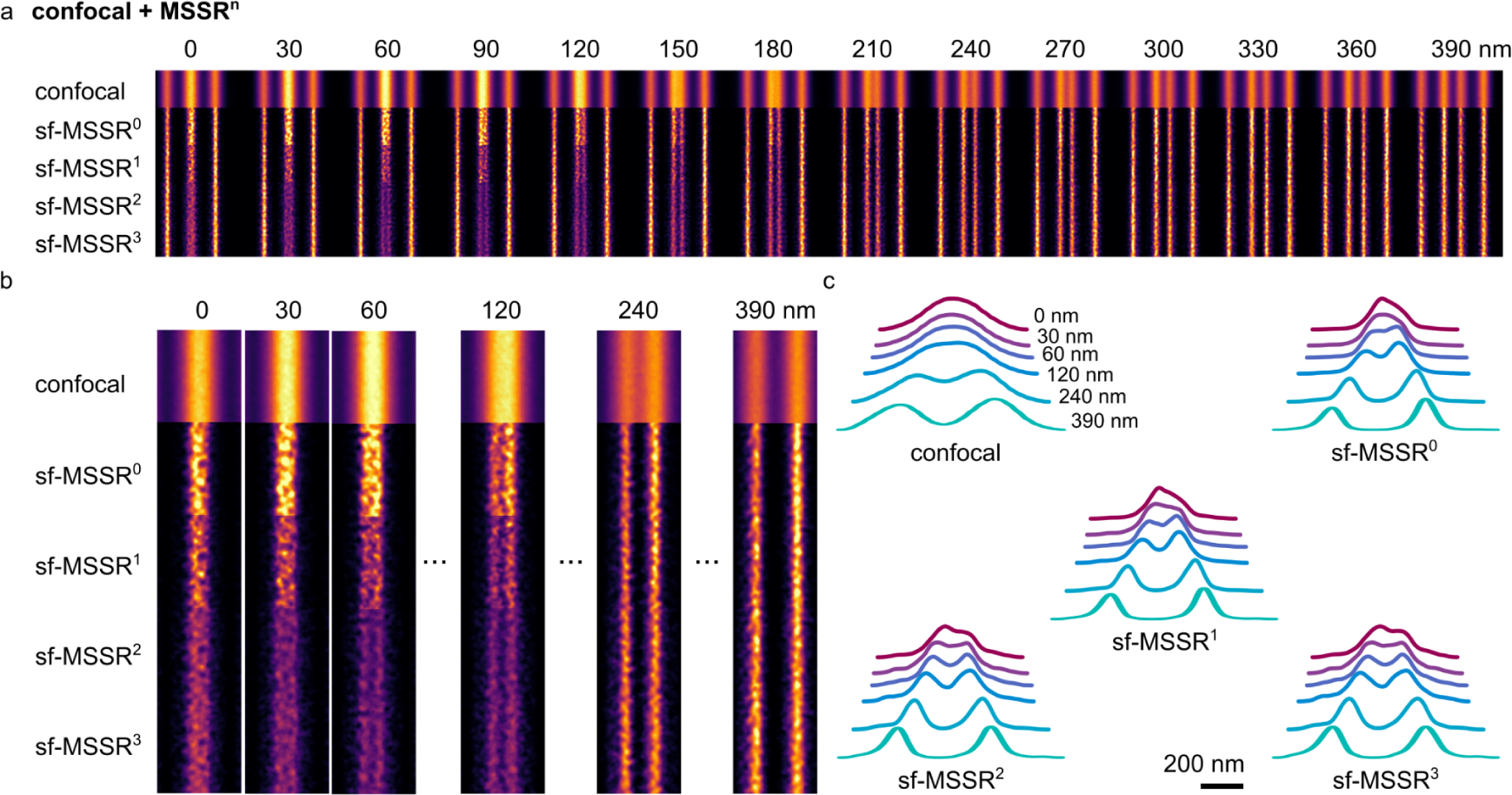
sf-MSSR^n^ extends spatial resolution in confocal microscopy. **a)** Comparison of confocal and sf-MSSR^n^ reconstruction (n=0-3) of ArgoLight test slide applied to a spaced fluorescent line pattern. Central lines are gradually being separated by 30 nm (0 nm, 30nm, 60 nm, … 390 nm). **b)** Results of confocal and sf-MSSR^n^ reconstruction (n=0-3) of line patterns separated at 0 nm, 30 nm, 60 nm, 120 nm, 240 nm and 390 nm. **c)** Average profiles of images obtained in **(b)**. Images were acquired using a Plan-Apochromat 63x/1.4 Oil immersion objective, exciting the Argo-SIM micropattern E with a 405 nm laser and detecting the fluorescence in the range 420 – 480 nm (Zeiss LSM880). The pixel size was 44 nm. Images in (a) a (b) correspond to the ensemble average of 31 consecutive sections (width = 1 µm) along the ArgoSIM micropattern E.

Higher orders of sf-MSSRn create a saddle point between parallel rows of emitters located in the range of 60 - 90 nm at the boundaries of the Rayleigh limit (Figure 3b, c).

### MSSR enhances the resolution of images with extended resolution

Based on the MSSR capabilities to generate a micrography with extended spatial resolution after processing a single fluorescence image, we explored if a pre-existing image with extended resolution can be further enhanced by sf-MSSR^n^.

The Argo SIM micropattern was imaged using an Airyscan detector with other experimental settings as in Figure 3. Images were further deconvolved with the corresponding Airyscan algorithm [39]. Figure 4 shows the application of sf-MSSRn to the Airyscan microscopy images of the Argo-SIM micropattern. Within the Airyscan processed image it is possible to resolve parallel rows of fluorophores located at 180 nm, but not 120 nm or less. sf-MSSR0 processing enhanced the spatial resolution of the same Airyscan data, allowing to discriminate parallel rows of fluorophores located at 120 nm, or less (Figure 4 a, b). Higher orders of sf-MSSR^n^ create a saddle point between parallel rows of emitters located in the range of 60 - 120 nm (Figure 4 b, c).

**Figure 4.**
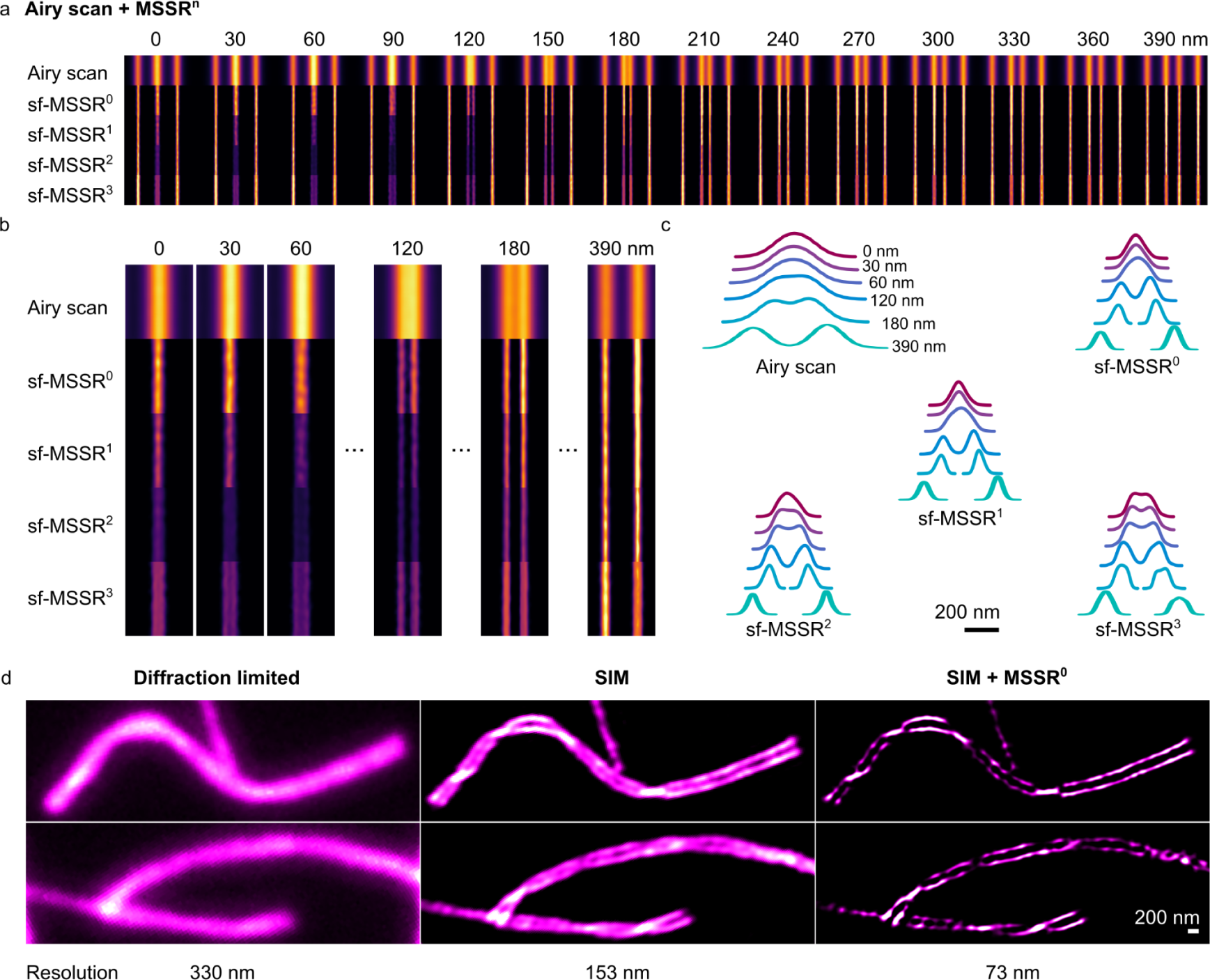
sf-MSSR^n^ enhances the resolution and contrast of Airy scan and SIM reconstructions. **a)** Comparison of confocal and sf-MSSR^n^ reconstruction (n=0-3) of ArgoLight test slide applied to a spaced fluorescent line pattern. Central lines are gradually being separated by 30 nm (0 nm, 30nm, 60 nm, … 390 nm). **b)** Results of confocal and sf-MSSR^n^ reconstruction (n=0-3) of line patterns separated at 0 nm, 30 nm, 60 nm, 120 nm, 240 nm and 390 nm. **c)** Average profiles of images obtained in (b). d) sf-MSSR^0^ applied to SIM reconstruction of chromosome axis. **d)** Synapsed homologs of meiotic mouse chromosomes visualized by TIRFM (left), SIM (middle) and SIM + MSSR^0^ (right). Images in (a) a (b) correspond to the ensemble average of 31 consecutive sections (width = 1 µm) along the ArgoSIM micropattern E. Resolution in **(d)** was measured with the image decorrelation plugin of FIJI/image [36].

Remarkably, compared with the confocal original data in Figure 3, where the last resolvable line pair is the one corresponding to a distance of 240 nm, the value of 120 nm obtained in Figure 4 by applying sf-MSSR^0^ to Airyscan processed data corresponds to a 2-fold improvement in resolution. The first reported applications of Airyscan technology allowed an improvement in resolution of 1.7X [40], while only following protocols claim that a 2-fold improvement might be achieved [39], compared to standard confocal detection.

We then applied sf-MSSR^0^ on a SIM image of sister meiotic chromatids of mouse chromosomes [41]. Similar to the results obtained above with Airyscan, Figure 4d shows that processing with SIM images with sf-MSSR^0^ enhances both the contrast and resolution of the final image.

### MSSR enhances the resolution of super-resolved images

We explored if a pre-existing super-resolved image can be further enhanced by sf-MSSR.

First, we used a temporal stack of DL images of tubulin-labeled microtubules collected at high fluorophore density [42] (previously used to test and compare a variety of SRM algorithms) [43], which were subject to FF-SRM analysis [10, 44]. ESI, SRRF or MUSICAL were used to compute a single enhanced-resolved image (Figure 5a) [11–13]. Supplementary Note 9 contains an in-depth comparison of sf-MSSR^0^ reconstructions combined with either ESI, SRRF and MUSICAL, which achieve super-resolution through a temporal analysis [11–14]. Post-processing of ESI, SRRF or MUSICAL images with sf-MSSR^0^ enhances contrast and spatial resolution (Figure 5a).

**Figure 5.**
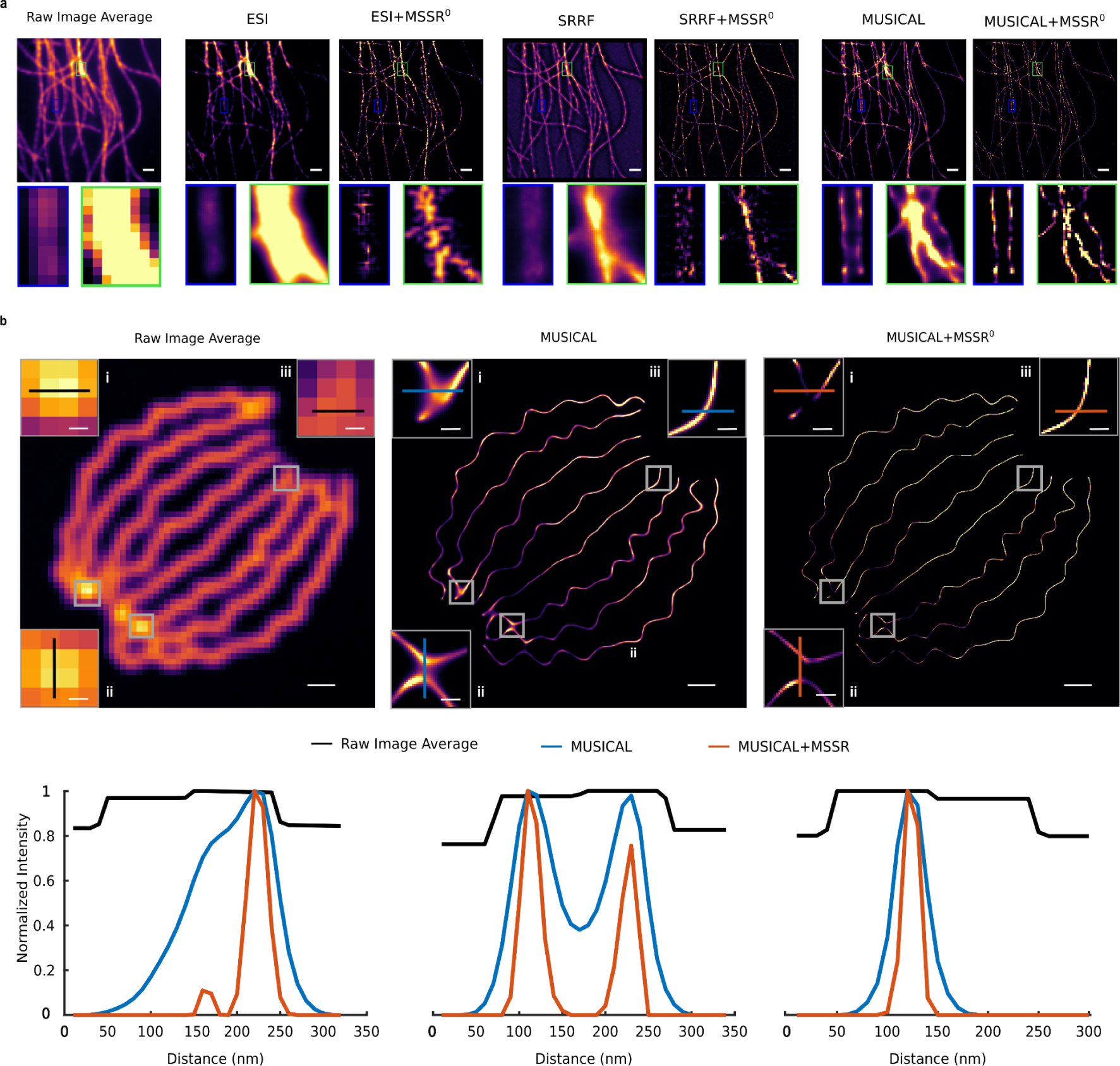
MSSR enhances resolution and contrast of SIM or enhanced-resolved images. **a)** Comparison of SRM results of ESI, SRRF and MUSICAL alone and after post-processing with MSSR^0^ (ESI + sf-MSSR^0^, SRRF + sf-MSSR^0^, MUSICAL + sf-MSSR^0^), over a temporal stack of 500 DL images of tubulin-labeled microtubules. The average projection of the DL stack is shown on the leftmost side. b) Comparison of the increase in spatial resolution of MUSICAL with and without post-processing with MSSR^0^ (MUSICAL + sf-MSSR^0^), over a temporal stack of 361 DL images of modeled fluorophores bounded to a synthetic array of nanotubes (average projection shown on left). The graphs show the intensity profiles along the lines depicted in each of the insets in the images of the upper row; black, blue and red lines correspond to the average DL, MUSICAL and MUSICAL + sf-MSSR^0^ images, respectively. Scale bars: **a)** 200 nm; **b)** 1 μm, insets = 200 nm; **c)** 500 nm, insets = 100 nm. MSSR parameters: AMP = 1, PSF FWHM = 2, order = 0; ESI parameters: two iterations, first iteration: image output = 250, bin = 50, order = 2, second iteration: image output = 50, bins = 5, order = 2; SRRF parameters: default parameters; MUSICAL parameters: lambda_em = 650 nm, NA = 1.4, pixel size = 100 nm, threshold = - 0.6, α = 4, subpixel per pixel = 4.

Second, a sequence of images of randomly blinking emitters placed along a synthetic tubular structure [45] was processed with sf-MSSR^0^ after analysis with MUSICAL. In both reconstructions, three regions (small squares in Figure 5b) were chosen to assess the gain in resolution, visualized in terms of the distance between the normalized intensity distributions peaks. MSSR further resolves the edges of the synthetic structures on the MUSICAL-processed image without changing the position of the distribution peaks (Figure 4c) as predicted by our theory (Supplementary Notes 1 to 4).

Lastly, we set up an experimental assay to examine the achievable resolution by MSSR alone, or in combination with either confocal or stimulated emission depletion (STED) microscopy. Figure 6 a, b, show that STED, but not confocal imaging allows to discern chromosomal territories within the chromatin of 2-cell stage embryo, as observed by the lack of colocalization of acetylated (ac, i.e., ‘active’) or methylated (me, i.e., ‘inactive’) chromatin states, H3K27me3, and H3K27ac respectively [46].

**Figure 6.**
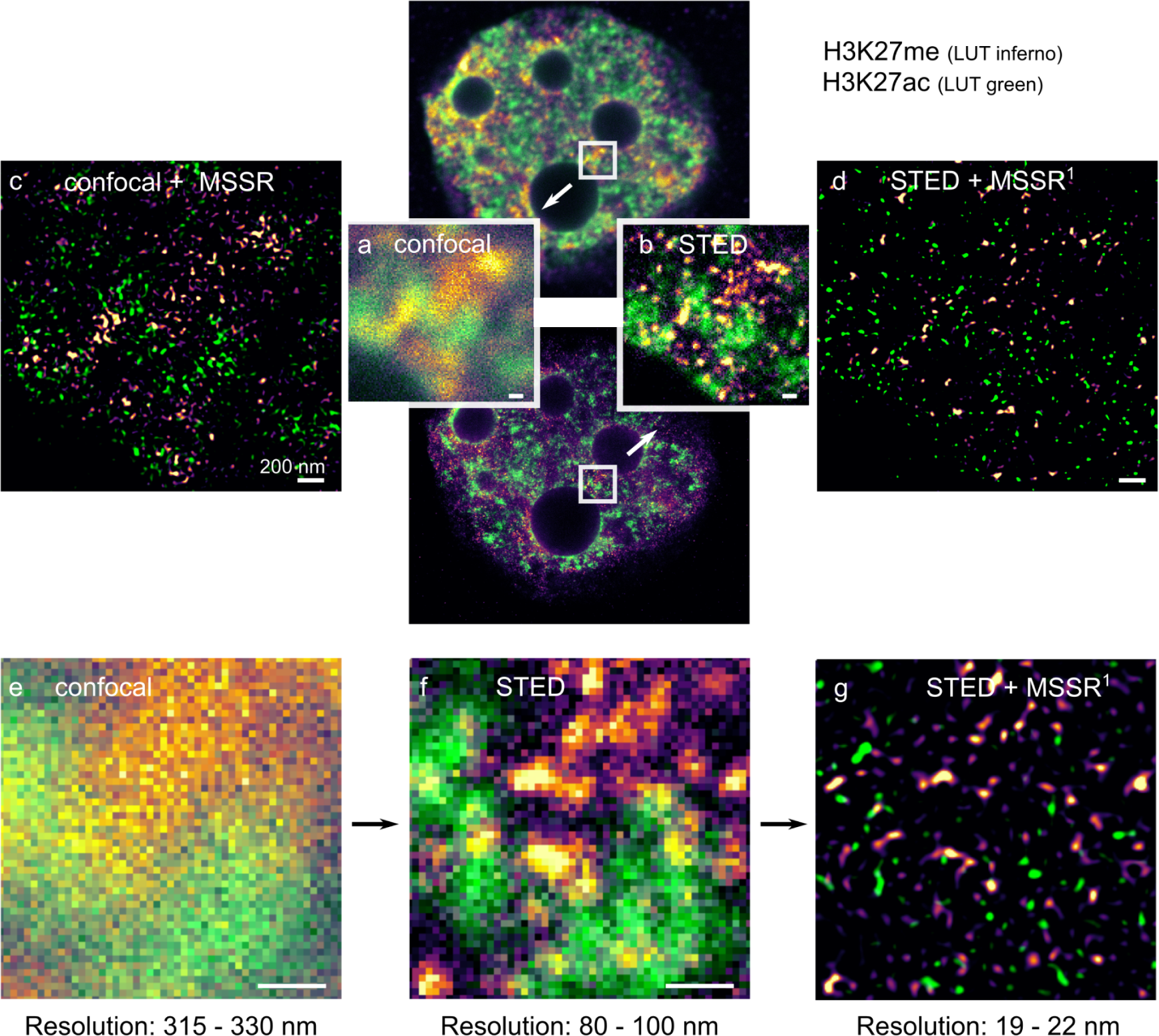
sf-MSSR^n^ enhances spatial resolution in STED microscopy. **a)** Confocal and **b)** STED micrographs of fluorescent histone proteins (H3K27me3, H3K27ac) imaged in a 2-cell stage mice embryo (top-center). **c)** and **d)** show the corresponding enhanced-resolved scenes provided by post-processing of the confocal and STED images with sf-MSSR^1^, respectively. The bottom row shows the sequential increase in resolution when comparing the same sample imaged with a **(e)** confocal microscope, an **(f)** STED microscope and **(g)** after processing the STED image with sf-MSSR^1^. For each image, the resolution range (in nanometers) was assessed using the ImDecorr algorithm in ImageJ [36]. Immunofluorescence Imaging was carried out using a STEDYCON mounted on an upright Zeiss microscope in confocal or STED modes. Samples were imaged with a 20 nm pixel size. Primary antibodies: anti-H3K27me3 (Abcam ab6002) and anti-H3K27ac (Active Motif, 39034). Secondary antibodies: anti-mouse labeled with STAR Red and anti-rabbit labeled with STAR Orange. The parameters used for sf-MSSR processing were AMP = 5, order = 1, FWHM of PSF = 4, interpolation = bicubic, mesh-minimization = true.

Remarkably, Figure 6c shows that sf-MSSR^1^ processing of the confocal image allows to reach a similar experimental conclusion, that there is no colocalization between H3K27me3 and H3K27ac signals.

Figure 6 also shows the result of post-processing of the STED image with sf-MSSR1 (Figure 6g). We note an increase in contrast and resolution from confocal to STED and then to STED + sf-MSSR^1^ (compare panels e, f, and g in Figure 6). To assess the resolution change provided by either STED or STED + sf-MSSR^1^ we used the image decorrelation approach [36]. The bottom panels of Figure 6 e, f, and g, show the resolution computed at a confocal image (315 – 330 nm), STED (80 – 100 nm) and STED + sf-MSSR^1^ (19 – 22 nm). With these results, we conclude that sf-MSSR^1^ processing of STED data provides a further increase in spatial resolution.

### MSSR achieves super resolution by analyzing fluorescence intermittency over time

Analyzing the temporal dynamics of fluorescence is central to achieving the highest attainable gain of resolution by FF-SRM approaches (reviewed in [36]). FF-SRM methods analyze higher order temporal statistics looking to discern correlated temporal information (due to fluorescent fluctuations) from uncorrelated noise (i.e., noise detector). In theory, MSSR can be applied to a sequence of images (Supplementary Note 5). Based on the increase in resolution offered by FF-SRM approaches (SRRF, MUSICAL), we investigated whether a further resolution gain could be achieved by applying a temporal analysis to a sequence of single-frame MSSR images (denoted by t-MSSR^n^) (Figure 7a). Pixel-wise temporal functions (*PTF*), such as average (*Mean*), variance (*Var*), the temporal product mean (*TPM*), coefficient of variation (*CV*) or auto-cumulant function of orders 2 to 4 (*SOFI_2_*, *SOFI_3_*, *SOFI_4_*) [10], can be used to create an image with enhanced spatial resolution (Supplementary Note 5.2, Supplementary Table S2).

**Figure 7.**
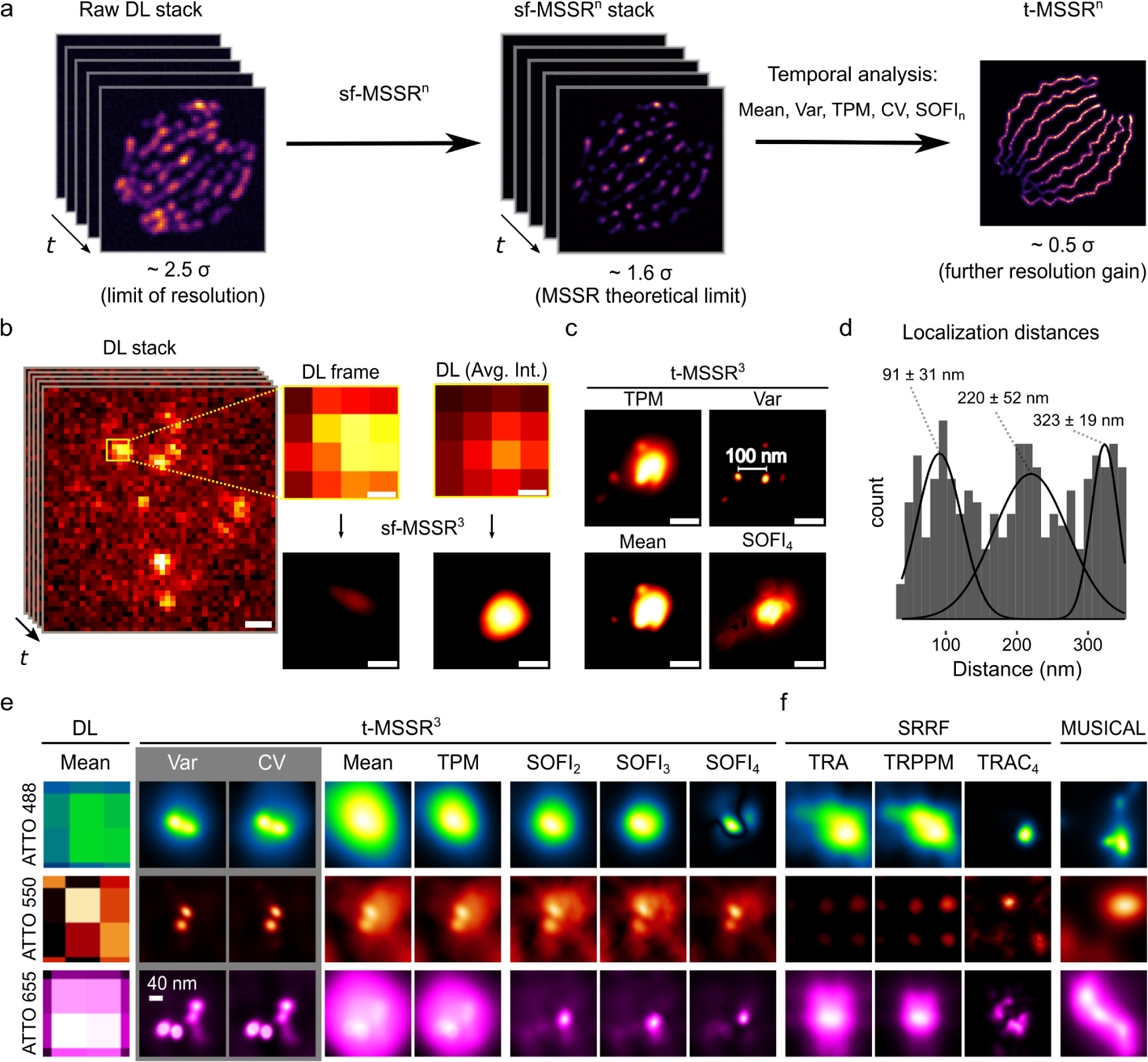
The temporal analysis of MSSR provides a further increase in resolution to 40 nm. **a)** Single-frame analysis of MSSR of a given order *n* is applied to each frame of a sequence, becoming the sf-MSSR^n^ stack. Next, a pixel-wise temporal function (PTF) converts the MSSR stack into a single super-resolved t-MSSR^n^ image. Depending on the temporal fluorescence dynamics of the dataset and on the PTF used, a resolution increase is obtained. **b)** Left: a stack of DL images of a CRISPR/dCas12a nanoruler system. Right: zoomed region of the first frame in the stack, along with the average projection (DL-AVG) of a stack of 100 images, before and after MSSR processing. **c)** PTF applied to a stack of MSSR^3^ images (t-MSSR^3^). Fluorescent emitters are separated by 100 nm, as established by the CRISPR/dCas12a nanoruler system. Four types of PTF were computed: *TPM*, *Var*, *Mean* and *SOFI_4_*. MSSR parameters: AMP = 20, FWHM of PSF = 3.74, Order = 3, Number of images for PTF: 300. Pixel size of the DL dataset = 100 nm. **d)** Euclidean distances between nearby emitters automatically computed from t-MSSR^3^-*Var* images, following a worm-like chain model (16 regions of interest used, 1.5 µm^2^ each). **e)** t-MSSR^3^ analysis (see Table S3) for a commercially available GATTA-PAINT nanoruler system. The *Var* column shows inter-emitter distances resolved at 40 nm. ATTO 488 (green), ATTO 550 (orange) and Atto 655 (magenta) fluorescent probes were used. **f)** Same nanorulers shown in (e) but analyzed using SRRF and MUSICAL. MSSR parameters: AMP = 20, FWHM of PSF for Gattpaint ATTO 488 = 2.79, Gattapaint ATTO 550 = 3.31, Gattapaint ATTO 655 = 3.74, Order = 3, Number of images for PTF: 300. Nano J - SRRF parameters: bicubic interpolation magnification = 2, ring radius = 1, radiality magnification = 10, axes in ring = 8, PTF = TRA, TRPPM, TRCA_4_, Other parameters were setted as default. MUSICAL parameters: emission wavelength = same as ATTO emission wavelength (520, 575, 680) used for each row in (e) and (f), NA = 1.49, pixel size = 100 nm, threshold = -0.3 (ATTO 488), -0.02 (ATTO 550), -0.2 (ATT0655), α = 4, subpixel per pixel = 20. frames = 300, musicJ v0.94 of imageJ.

To experimentally validate the increase in resolution by both sf-MSSR^n^ and t-MSSR^n^ we used two different nanoruler systems, an in-lab CRISPR/dCas12a nanoruler, used to score nanoscopic distances between individual fluorescent sites down to 100 nm, and a commercial nanoruler with fluorophores positioned at 40 nm of separation (GATTA-PAINT, 40G, and 40RY. Gattaquant).

The CRISPR/dCas12a nanoruler system consists of a dsDNA with four binding sites for dCas12a uniformly distributed every 297 bp (equivalent to ∼ 100 nm of separation) (Supplementary Figure S42a). To validate this system, we imaged the association of the CRISPR/dCas12a complex to the binding sites on the dsDNA by atomic force microscopy (AFM) and measured the distance between each dCas12a complex (Supplementary Figure S42b).

The CRISPR-dCas12a nanorulers were then imaged by TIRF microscopy for further MSSR analysis. We used a DNA-PAINT approach for fluorescence indirect tagging [47], in which a fluorescent ssDNA probe hybridizes with an extension of the gRNA. The “blinking” of the fluorescence signal is attained by events of association and dissociation between the fluorescent probe and the gRNA on the CRISPR/dCas12a nanoruler at the binding site.

In the DL image, amorphous spot-like fluorescent patterns were observed (Figure 7b). sf-MSSR^3^ processing of either an isolated frame or an average projection of the corresponding stack of 100 images (DL-AVG) could not resolve individual CRISPR/dCas12a binding sites (Figure 6b), and only after processing by t-MSSR^3^ did individual binding sites became resolved (Figure 7c). The result of t-MSSR^3^ varied in relation to the temporal function used (Figure 7c). The best result for this nanoruler was obtained by the pixel-wise temporal variance (*Var*) of the sf-MSSR^3^ stack (Figure 6c). t-MSSR^3^-*Var* resolved nearby emitters engineered to recognize binding sites located at 100 nm (Supplementary Movie S1), provided by scoring association-dissociation events between the imaging probe and the gRNA.

Once the CRISPR/dCas12a nanorulers were super-resolved by t-MSSR^3^-*Var* (Figure 7c), we scanned all the individual emitters in close proximity to determine the average distances between different dCas12a associated to the dsDNA binding sites (theoretically interspaced by 100 nm, Supplementary Figure S42). Since DNA in solution is a semi-flexible polymer [48], the measured distances resulted in three different distributions: 91 ± 31 nm, 220 ± 52 nm, 323 ± 19 nm (Figure 7d), which accounts for dCas12a separated by one, two and three binding sites along the dsDNA, respectively. These results confirm that t-MSSR^3^ can successfully resolve nanoscopic distances.

To explore the resolution limit attainable by t-MSSR^n^ even further, we looked at a nanoruler system with smaller separation between fluorophore sites (from Gattaquant) (Supplementary Figure S43a). Figure 7e shows that using the TRA or TRM pixel-wise temporal functions in combination with MSSR^n^ (t-MSSR^n^) does not provide extra resolution enhancement. TRA provides t-MSSR^3^ reconstructions less influenced by noise (recommended in imaging scenarios of marginal signal-to-noise ratios) and TRM delivers reconstructions whose intensity distribution is less dominated by constantly emitting sources, i.e., fiducial markers. In stark contrast, higher spatial resolution regimes, ∼ 0.15 times the FWHM of the PSF (PSF σ: [0.4 - 0.5]), are achieved by t-MSSRn when encompassing pixel-wise temporal functions of higher-order statistics such as Variance (Var), Coefficient of Variation (CV), or SOFI (TRAC2-4). Analysis with t-MSSR^3^ of 300 DL images using either Var or CV revealed individual fluorescent spots at 40 nm apart (Figure 7e and Supplementary Figure S43b). The data presented in Figure 7e demonstrate that t-MSSR^3^ resolves nanoscopic distances at 40 nm, validating a lower experimental spatial resolution bound of 0.5 σ (≈ 40 nm), which depends on the emission wavelength of the fluorophore (Figure 7e, Supplementary Figure S10c). In comparison, SRRF and MUSICAL were not able to resolve fluorescent emitters located 40 nm apart, consistent with their limit within the range of 50–70 nm (Figure 7f) [11–13].

Analyzing the temporal dynamics of fluorescence intermittency is central for any FF-SRM approach [10–13]. The highest attainable spatial resolution is influenced by blinking statistics, i.e., photokinetics for fixed fluorophores [11-13,49], and binding energies for diffusible fluorescent probes [50]. These factors, including the density of fluorophores emitting per frame, impinge on the final resolution of the reconstruction. Supplementary Figure S44 shows the pixel-wise temporal dynamics of two GATTA paint ranorulers (ATTO 655), where the temporal dynamics of a given nanoruler containing three binding sites can be studied by analyzing the temporal fluorescence fluctuations at nearby pixels. Note that diffraction imposes constraints to distinguish the temporal dynamics at individual PAINT binding sites. Noteworthy, sf-MSSR^3^ processing allows to untangle, in space and in time, the fluorescence dynamics at individual PAINT binding sites, where the fluorescence signal peaks due to the transient binding of a fluorescent labeled DNA strand to its corresponding binding site within the nanoruler.

### Single frame nanoscopy, free of noise-dependent artifacts

The theory of image processing by MSSR (Supplementary Note 5), suggests that it should be robust over a wide range of SNR, granted by four factors. First, when processing a single frame, MS works as a local spatial frequency filter (a smoothing filter); regions corresponding to the image background are homogenized by the kernel window, reducing variation in background noise. Second, one of the steps of the MSSR procedure is to remove the MS negative constraints. The goal of this step is to remove an artifact caused by the MS calculation itself when applied to Gaussian and Bessel distributions. When calculating the complement of the resulting distribution, a valley (or depression) is generated between the peak intensity and the tails, which lies in the negative values and is referred to as negative constraints. After this, the artifact is removed. Third, when using a PTF, nanoscopic information is enriched due to temporal oversampling of the hidden fluorescent structure. Fourth, the spatial kernel of the MSSR algorithm operates within the subpixel realm; the number of neighboring pixels is digitally increased through interpolation (i.e., bicubic interpolation [51] for single frame analysis and Fourier interpolation [52] for temporal analysis, Supplementary Note 6) providing digital oversampling of the emitters’ locations (Supplementary Note 6).

We then experimentally assessed the capacity of MSSR to denoise fluorescence images and determine whether it introduces noise-related artifacts. We used a PSFcheck slide [53], which contains an array of regular fluorescent nanoscopic patterns shaped by laser lithography (Figure 8). Analysis with sf-MSSR^n^ or t-MSSR^n^ showed, in comparison to alternative approaches, striking denoising capabilities without introducing noticeable artifacts (Figure 8a, Supplementary Note 9). These artifacts, resembling amorphous nanoscopic structures around the fluorescent ring or within it, were commonly found at reconstructions generated by other analytical techniques (Supplementary Figure S29).

Starting at a SNR > 2, sf-MSSR^1^ provides reliable SRM reconstructions of comparable quality to other SRM approaches, which demand the temporal analysis of the fluorescence dynamics (Figure 8a and Supplementary Note 9). We quantified the quality of the reconstructions by calculating the Resolution Scaled Pearson (RSP) coefficient and the Resolution Scaled Error (RSE), which provide a global measurement of the quality of the reconstruction by comparing the super-resolution image and the reference image (in this case, the DL image) [14]. Higher RSP and lower SRE values are associated with reliable reconstructions (Supplementary Note 8). When the SNR is above 5, all tested algorithms perform similarly well in quality (Figure 8b), but their global errors differ from each other (Figure 8c). As expected, the RSE increased as a function of the SNR of the input images for any tested algorithm (Figure 8c).

The performance of MSSR in achieving a satisfactory reconstruction was assessed by varying the number of input images using a temporal analysis scheme (Supplementary Note 8). With SNR > 2 input data, RSP reaches near maxima values and RSE near minima values when processing a single frame (Supplementary Figure S28, Supplementary Movie S2). However, when computing MSSR using low SNR input data (SNR ∼ 2) a temporal analysis is required as RSP and RSE values reach a plateau only when a temporal stack of as few as 25 images is used (Supplementary Movie S3). These findings illustrate that the minimal number of frames needed by MSSR to provide a reliable reconstruction depends on the information itself, i.e., on the SNR and on the fluorescence blinking statistics of the specimen (Supplementary Movies S1-S3); and can be determined by computing RSP and RSE as function of the number of analyzed frames with t-MSSR^n^ (Supplementary Figure S28).

### Nanoscopic resolution with conventional fluorescence imaging

MSSR addresses the problem of spatial resolution in optical fluorescence microscopy from the statistical point of view, where the process of photon emission from punctual sources is considered as a discrete distribution of information, provided by detected photons. Enhanced or extended resolution is achieved by finding local modes of information without taking any prior about the local shape of the distribution. The latter makes the MSSR principle compatible with any imaging approach where the distribution of information is of discrete nature (i.e., photons), and emanates from discrete sources (i.e., fluorophores).

To showcase the versatility of MSSR to extended-, enhanced- and super-resolve data acquired from different fluorescence applications, we evaluated its performance over a collection of experimental scenarios (Figure 9) (Supplementary Note 10).

**Figure 9.**
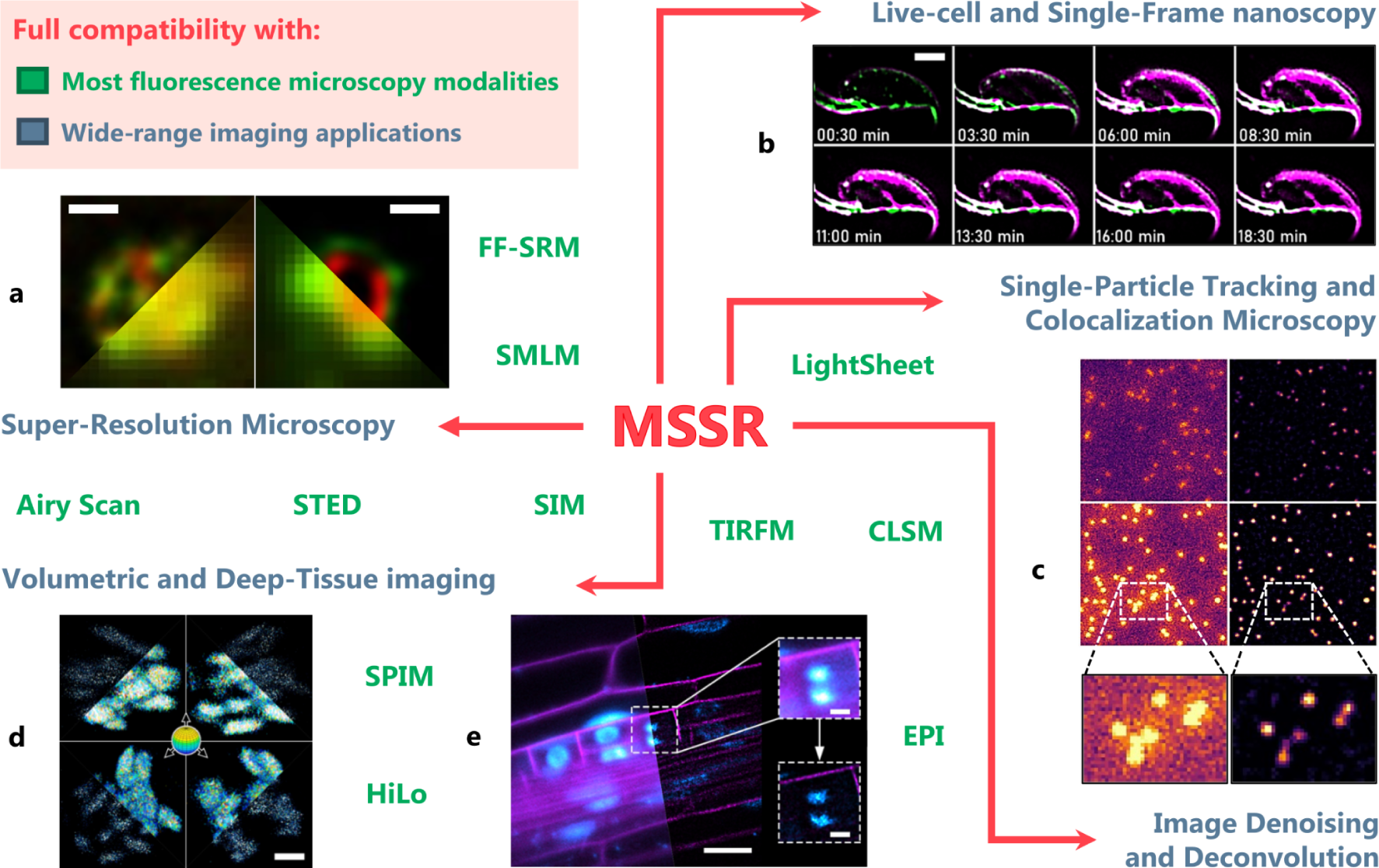
MSSR applications in fluorescence microscopy. MSSR operates over images acquired with most fluorescence microscopy modalities available (e.g., widefield, confocal, light-sheet, etc.), denoted by the text in green. It can be applied to achieve enhanced image resolution, SNR and contrast in live-cell imaging, single-particle tracking, deep tissue and volumetric imaging, among others (denoted by the text in pale blue). Some examples of these applications are detailed in the main text and Supplementary Note 10. Abbreviations: FF-SRM, Fluorescence Fluctuation-based Super-Resolution Microscopy; SMLM, Single-Molecule Localization Microscopy; STED, Stimulated Emission Depletion microscopy; SIM, Structured Illumination Microscopy; SPIM, Selective Plane Illumination Microscopy; EPI, Epifluorescence; HiLo, highly inclined and Laminated optical microscopy; TIRFM, Total Internal Reflection Fluorescence Microscopy; CLSM, Confocal Laser-Scanning Microscopy. Scale bars: a) 500 nm; b) 2 μm; c) no scale provided; d) 2 μm; e) 20 μm, insets = 5 µm.

Analysis with MSSR provided nanoscopic resolution of rotavirus replication machineries (Figure 9a, Supplementary Figure S32), which were recently described by Garcés *et al* as a layered array of viral protein distributions [54]. Originally, it took the authors several days to weeks to generate a single extended-resolution image by means of analyzing several stacks of hundreds of DL images using 3B-ODE FF-SRM. With MSSR, we were able to achieve comparable results, through analyzing single DL frames within seconds with a regular desktop computer with either sf-MSSR^1^ or t-MSSR^1^ (Supplementary Note 7).

Mouse sperm cells are used to study the acrosomal exocytosis (AE), a unique secretory process which results from fusion events between the plasma membrane and a specialized vesicle called acrosome located in the sperm head [55, 56]. Nanoscopic remodeling of both plasma membrane and actin cytoskeleton was imaged during the AE by means of sf-MSSR^1^, showing single frame temporal resolution (of milliseconds) (Figure 8b, Supplementary Figures S33-34). At the onset of the AE, the FM4-64 fluorescence (a probe that fluoresces when bound to membranes) was confined to the plasma membrane and was visible above a F-actin cytoskeleton fringe. During the AE, several fenestration events were observed to occur at both the plasma and acrosome membranes, as consequence of that, a notorious increase of FM4-64 was observed close-bellow the F-actin fringe (Supplementary Movie S5 a-f). The AE is a dynamic remodeling process that takes minutes to occur, sf-MSSR^1^ allows the observation of events occurring at the millisecond scales, which are hindered when using other SRM multi-frame analytical approaches, such as SRRF or 3B [11, 57], due to their mandatory need of a temporal analysis of the fluorescence dynamics to unveil nanoscopic detail (compare Supplementary Figures S33 and S34).

Background noise is known to be an important issue in single-particle tracking (SPT) applications as it decreases the ability to faithfully localize particles and follow them through time [58, 59]. Moreover, the spatial overlap of PSFs derived from individual particles makes it challenging for SPT algorithms to recognize them as separate entities. The denoising capabilities of sf-MSSR^1^ enhanced both the contrast and spatial resolution of freely diffusing in-silico particles (Figure 8c, Supplementary Figure S35), previously used as benchmarks to test a variety of SPT algorithms [60].

Pre-processing of the images with sf-MSSR^1^ improved the tracking performance of three commonly employed SPT tracking algorithms: (i) the LAP framework for Brownian motion as in [61, 62], (ii) a linear motion tracker based on Kalman filter [63, 64], and (iii) a tracker based on Nearest neighbors [65–68] within a wide range of particle densities and SNR (Supplementary Figure S36). Additional testing with sf-MSSR^n^ showed an increase in contrast for moving comet-like particles related to microtubule growth dynamics in live LLC-PK1 cells (Supplementary Figure S37 and Supplementary Movies S10 and S11), as well as higher nanoscopic colocalization accuracy in double imaging experiments in single-molecule DNA curtain assays (Supplementary Figure S38) [69].

Plasmalemma- and nuclear-labeled transgenic *Arabidopsis thaliana* plants are routinely used to study cell fate and proliferation during root development in time-lapse confocal microscopy experiments in two and three dimensions [70, 71]. When applied to lateral root primordium cells, located deep inside the parent root, sf-MSSR^1^ demonstrated the capacity to achieve multidimensional nanoscopic resolution as it revealed isolated nanodomains resembling nucleosome clutches, previously reported in mammalian cells [72, 73], within the nuclei of a lateral root primordium cells (Figure 8d-e, Supplementary Figure S39 and Supplementary Movie S12). Similar observations were performed upon epidermal root tissues visualized via selective plane illumination microscopy (SPIM) after examination of volumetric data with sf-MSSR^1^ (Supplementary Figure S40).

Supplementary Figure S41 shows a comparison of 3D reconstruction from a stack of epifluorescence images of bovine pulmonary artery endothelial (BPAE) cells. When a 2D image is processed by sf-MSSR^1^, an improvement in spatial resolution and contrast is observed (Supplementary Figure S41a,b). On the contrary, when a stack is used for a 3D representation (images taken at z-planes), the resolution obtained by sf-MSSR along the z-axis is not as refined as in the xy-plane (Supplementary Figure S41c,d). Nonetheless, a noticeable increase in contrast is attained in the overall dataset (Supplementary Figure S41, see Supplementary Movies S13 and S14). To further extend the resolution of DL images in 3D, it is necessary to extend the current implementations of the MSSR algorithm to account 3D information for MSSR processing.

In combination, these studies provide evidence for the capabilities of MSSR to resolve biological detail at nanoscopic scales using either simple or advanced fluorescence microscopy technologies.

## Discussion

### Novel theoretical contributions

From the historical point of view, since the seminal development of the MS theory [17, 18] and until the present day, few statistical and imaging applications based on the theory of MS compute the MS vector itself [74]. This can be explained, in part, because previous applications of MS are based on finding modes in the features space but did not calculate the MS vector, which is the key component of the working principle of MSSR. In contrast, MSSR represents an application of MS theory which also operates in the second derivative space. By computing the MS vector and estimating (photon) densities among pixels, MSSR computes a probability function for the fluorophore estimates whose individual fluorescence distributions are narrowed in comparison with the PSF of the optical system. The exploration of the information stored on the second derivative space of the image can be also achieved by substituting the MS by similar functions that operate in such space, e.g., Laplacian, Hessian, Difference of Gaussians [75] which, in comparison with the MS, offer computational advantages as they can be expressed in the Fourier space and implemented using the FFT algorithm [75]. The information harbored in the second derivative space of the DL image is used by MSSR to compute an image with higher spatial frequencies than the corresponding DL image, hence, overcoming both the Rayleigh and Sparrow limits, and setting up an undescribed limit of spatial resolution which deserves further exploration and characterization.

The MS theory is not restricted by the number of dimensions of the information required to compute the kernel windows over which MSSR operates (Supplementary Notes 2 and 3). Given that, MSSR parameters are suitable to extend its application to assess data with higher dimensions. For example, in 2D images, the spatial parameter of MSSR, which encompasses the lateral resolution width of the PSF, is defined to be the same in the x and y dimensions of the image. In such a case, the shape of the kernel is circle- or square-like, depending on the application used. For three-dimensional (3D) microscopy imaging, the lateral (x-y plane) and axial (x-z and y-z planes) dimensions are affected in different ways by diffraction. The MSSR principle can be further extended for explicit volumetric imaging by means of using an asymmetric kernel which can be defined following the 3D lateral-axial aspect ratio of the PSF (Supplementary Note 11). In addition, the definition of the spatial kernel can be refined to also consider possible deformations of axial symmetry of the PSF due to optical aberrations introduced by the imaging system or by the sample itself. A similar reasoning aimed to extend the portfolio of applications of MSSR can be envisaged considering spatial-range parameters, the latter narrowing down the working intensity space where local calculations of MSSR take place.

### Novel contributions to microscopy

We present a new SRM approach capable of achieving multidimensional nanoscopy through single-frame analysis under low SNR conditions and with minimal noise-dependent artifacts. Limited only by the imaging speed of the optical system setup, MSSR increases resolution by analyzing either a single frame, or by applying MSSR to each individual image in a stack followed by the application of a pixel-wise temporal function. MSSR is a powerful stand-alone method for either single or multi-frame SRM approaches, or as a post-processing method which can be applied to other analytical multi-frame (restricted to camera-based systems) or hardware dependent SRM methods for further enhancement of resolution and contrast. We demonstrated MSSR compatibility with other SRM methods and showed that its usage enhanced resolution and overall image quality in all the cases tested.

FF-SRM analytical multi-frame approaches such as SRRF, ESI, MUSICAL and 3B demand a temporal analysis which limits their utility for multi-dimensional imaging of live samples [65]. The need to collect hundreds to thousands of images of the same pseudo-static scene, challenges the applicability of these methods in multidimensional imaging. The temporal multi-frame requirement imposes a tradeoff between the achievable temporal and spatial resolutions. MSSR removes these constraints while maintaining computational efficiency (Supplementary Note 7).

We present applications of the MSSR principle that revealed fast molecular dynamics through single-frame analysis of live-cell imaging data, with reduced processing times in comparison with similar SRM approaches (Supplementary Notes 7 and 10). Moreover, MSSR greatly improves the tracking efficacy of SPT methods by means of reducing background noise and increasing both the contrast and SNR of noisy SPT movies, enhancing the ability to resolve the position of single emitters. MSSR further pushes the limits of live-cell nanoscopy by its excellent single-frame performance. This flexibility extends its utility to most fluorescence microscopy and alternative SRM methods.

Achieving both high (or sufficient) temporal and spatial resolution within a broad range of fluorescence microscopy applications is a common goal among the bioimaging community. With recent advances in microscopy equipment and imaging protocols, the gap between the highest attainable resolution in the temporal and spatial dimensions within the same experiment, has narrowed. This has been a challenge especially because both parameters often involve mutually exclusive optical instrumentation and experimental strategies. The introduction of MSSR represents one more step in the right direction as it drastically reduces the amount of data needed to reconstruct a single super-resolved micrography.

No longer having to sacrifice either temporal or spatial resolution over the other, has led some scientists to propose new ways to analyze imaging data. Some approximations have been tailored to study millisecond molecular dynamics and structural feature changes within the same experiment [76], e.g., by taking advantage of the simultaneous use of image correlation spectroscopy (ICS) and FF-SRM methods such as SRRF [11]. In these contexts, MSSR could improve the analysis in three ways: it delivers reliable SRM images in low SNR scenarios, which are common in the experimental regimes of ICS due to the relatively fast frame rates of its applications, b) MSSR introduces no noise-dependent artifacts which further refines the quality of the spatial analysis and c) since no temporal binning is necessary for MSSR, there is no restriction in the level of temporal detail retrievable from the ICS analysis.

Sub-millisecond time-lapse microscopy imaging can now be achieved by sCMOS technologies, with applications for particle velocimetry [77], rheometry [78], and optical patch clamp [79]. We envisage further applications for MSSR in these areas through unveiling nanoscopic detail hidden in single DL images. Moreover, MSSR can facilitate correlative nanoscopic imaging through crosstalk with other imaging techniques such as electron microscopy, i.e., CLEM: correlative light electron microscopy [80]; or atomic force microscopy, i.e., CLAFEM: Correlative light atomic force electron microscopy [81]. In addition, MSSR can be applied to nanoscopic volumetric imaging by using it together with expansion microscopy [82], oblique angle microscopy [83], SPIM and lattice light sheet microscopy [84], extending their capabilities to previous unattainable resolution regimes.

A recent study by Chen R. et al., suggests that deep-learning based artificial intelligence (AI) can reconstruct a super-resolution image from a single DL image [85]. Such AI-based SRM approaches are promising, however, they are limited to the existence of a maximum likelihood image obtained with another SRM, such as STORM, that is required for neural network training and error minimization. Otherwise, the method is prompted to bias the final reconstruction toward the topological information used to train the AI - network [85]. Our approach works completely independent of other SRM methods and provides evidence of the existence of a new resolution limit which lies on the second derivative space of the DL image, information inaccessible when using neural networks.

MSSR applications might impact far beyond the field of microscopy, as its principles can be applied to any lens-based system such as astronomy [86] and high-resolution satellite imagery [87].

## Supporting information

MSSR Supplementary Information

MSSR User Manual

movie s1 (CRISPR_PAINT nanorulers resolved by MSSR)

Movie S2 (t-MSSR sequential SNR 5.4)

movie s3 (Laser-written lithography pattern reconstruction at high SNR 10.4)

movie s4 (Super-resolution MSSR head of single frame)

movie s5 (Super-resolution MSSR fenestration of single frames)

Movie S6 (Super-resolution t-MSSR head of 100 frames)

Movie S7 (Super-resolution t-MSSR fenestration of 100 frames)

movie s8 (Super-resolution MSSR head of 100 frames)

movie s9 (Super-resolution MSSR fenestration of 100 frames)

Movie S10 (sf-MSSR video of live LLC-PK1 cells expressing mEmerald-EB3)

Movie S11 (sf-MSSR video of an apoptotic LLC-PK1 cell expressing mEmerald EB3)

Movie S11 (sf-MSSR video of an apoptotic LLC-PK1 cell expressing mEmerald EB3)

Movie S13 (DL reconstruction of epifluorescence BPAE cells for 2D and 3D images)

Movie S14 (sf-MSSR reconstruction of epifluorescence BPAE cells for 2D and 3D images)

## Acknowledgements

To the Microscopy Facility at EMBL Rome and IBYME-CONICET for providing samples and materials for experiments. To the ECE and ICIMAF institutes for theoretical and experimental input to this work. The authors thank the LNMA staff for providing support with microscopy imaging. The OpenSPIM project was technically executed and extended by Oliver Valdez Escalona. We thank the Analytical and Quantitative Light Microscopy course staff from the Marine Biological Laboratory for providing samples and data collection assistance. All authors thank Paul Hernández, Luis Mochan, Federico Lecumberry, Angelica Flores Navarrete and Yuriney Abonza Amaro for their valuable feedback during the elaboration of the manuscript and review process. The comments raised by both reviewers of the manuscript increased the overall scientific quality of the manuscript. Thank you all for making it possible.

## Author contributions

Conceptualization: ETG, AG.

Methodology: ETG, AG, RPC, VA, EBA, HTM, DT, AHC, RRM, JGD, RDA.

Investigation: ETG, AG, RPC, AHC, DM, VA, EBA, CCC, GVG, AHG, DT, JMR, YG, MGB, RRM, MJ, HTM, HOH, JOO, RDA, AB.

Visualization: ETG, AG, RPC, HTM, VA, EBA, AL, DT.

Supervision: AG, ETG, AHC, DK.

Funding Acquisition: AG, AD, MGB, JGD, CDW. Project Administration: AG.

Drafting Main Document: ETG, AG, AL, AHC, CDW.

Drafting Supplementary Material: ETG, AG, AL, RPC, RRM, JMR, DK, VA, EBA, HOH.

All authors contributed to the writing and reviewing of the manuscript.

## Competing interests

The authors declare no competing interests.

## Additional information

The online version contains supplementary methods, supplementary notes, supplementary movies, the MSSR manual for FIJI/ImageJ, MSSR scripts R, python MATLAB, and additional references.

## Data availability

All raw imaging data which support the findings of this study are available from the corresponding author upon request. Source data are provided with this paper.

Correspondence and requests for materials should be addressed to A.G.

## Code availability

Source code for R, python and MATLAB platforms is available as supplementary materials, the MSSR plugin for FIJI/ImageJ is available at https://github.com/MSSRSupport/MSSR.

## Funding

This research was supported by Dirección General de Asuntos del Personal Académico (DGAPA) – Programa de Apoyo a Proyectos de Investigación e Innovación Tecnológica-UNAM (PAPIIT). and by Mexican Consejo Nacional de Ciencia y Tecnología (CONACyT) grant A1-S-9236 to JGD, grant number IN211821 to AG, IN211216 to C.D.W., IN204221 to JGD. Microscopy equipment was provided and maintained through CONACYT grants 123007, 232708, 260541, 280487, 293624 and 294781. CW and AG acknowledge the Chan Zuckerberg Initiative, grants:

GBI-0000000093 (to CDW and AG) 2021-240504 (to MGB and AG). MGB thank to

IBYME-CONICET: Grants: PICT 2017-3047 (Agencia Nacional de Promoción de la Investigación, el Desarrollo Tecnológico y la Innovación), Fundación Williams; Fundación Rene Barón. DK acknowledges the support of the National Science Foundation grant 2102832. AL acknowledges the Howard Hughes Medical Institute (HHMI Award GT12050-2019). We also acknowledge the Programa de Becas de Posgrado of CONACYT for granting scholarships to ETG, RPC, AL, DM, VA, EBA, CCC, GVG, DT, HTM, JLM, HH, and to JPO. YG acknowledges DGAPA/UNAM for postdoctoral fellowship.

## Notes

https://github.com/MSSRSupport/MSSR

### Competing Interest Statement

The authors have declared no competing interest.

### Summary of Updates

The changes to the manuscript are the following: Main manuscript New figures: - Figure 3: sf-MSSRn extends spatial resolution in confocal microscopy, - Figure 4: sf-MSSRn enhances the resolution and contrast of Airy scan and SIM reconstructions. - Figure 6: sf-MSSRn enhances spatial resolution in STED microscopy. - Figure 9: Showcase of MSSR wide-range fluorescence microscopy applications. Other figures have been updated. Supplementary material A description of Fourier interpolation and its comparison with bicubic interpolation (see section 6.2). otencial adaptation of MSSR to 3D images has been added to Supplementary (see section 11). New supplementary figures: - Supplementary Figure S1: Behavior of ks according to the distance in a neighborhood. - Supplementary Figure S8: Selection of spatial parameter hs related to FWHM. - Supplementary Figure S13: Thresholding and normalizing processes are not enough to overcome the Sparrow limit. - Supplementary Figure S14: Resolution increase provided by deconvolution methods, radiality maps and sf-MSSRn. - Supplementary Figure S23: Fourier interpolation in 2D. - Supplementary Figure S24: sf-MSSR processing times for bicubic and Fourier interpolations. - Supplementary Figure S25: Comparison between Fourier and bicubic interpolation for temporal analysis on experimental dataset. - Supplementary Figure S37: Nanoscopic, single-frame, live-cell imaging of microtubule dynamics in LLC-PK1 cells. - Supplementary Figure S41: Comparison DL images of sf-MSSR reconstruction of epifluorescence BPAE cells for 2D and 3D images. - Supplementary Figure S44: Temporal fluctuation characteristics of two GattaPaint Nanorulers. Most of the figures have been updated with the mpl-infierno LUT using FIJI/ImageJ. Manual Figures 3, 5, 6, 8 and 9 have been updated. Movies New movies: - Supplementary Movie 10: sf-MSSR video of live LLC-PK1 cells expressing mEmerald-EB3. - Supplementary Movie 11: sf-MSSR video of an apoptotic LLC-PK1 cell expressing mEmerald-EB3. - Supplementary Movie 13: DL reconstruction of epifluorescence BPAE cells for 2D and 3D images. - Supplementary Movie 14: sf-MSSR1 reconstruction of epifluorescence BPAE cells for 2D and 3D images. - Other movies have been updated. MSSR Plugin Fiji/ImageJ and other codes With the new changes MSSR Plugin has been updated to version 2.0.0. "PSFp" parameter has been redefined as "FWHM of PSF". An option to select the type of interpolation has been added. An option related to perform intensity normalization on sequence of images has been added. Codes for Matlab, R, and Python have been updated according to the new changes on the MSSR theory.

## References

1. Galbraith, C. G. & Galbraith, J. A. Super-resolution microscopy at a glance, .J. cell science, 124, 1607–1611, (2011).

2. Thorley, J. A., Pike, J. & Rappoport, J. Z. Chapter 14 - Super-resolution Microscopy: A Comparison of Commercially Available Options. In Cornea, A. & Conn, P. M. (eds.) Fluorescence Microscopy, 199–212, (Academic Press, Boston, 2014).

3. Hell, S. W. & Wichmann, J. Breaking the diffraction resolution limit by stimulated emission: stimulated-emission-depletion fluorescence microscopy. Opt. Lett. 19, 780–782, (1994).

4. Prakash, K., et al. Super-resolution structured illumination microscopy: past, present and future. Phil. Trans. R. Soc. A. 379: 202001432020014, (2021).

5. Manton J. Answering some questions about structured illumination microscopy. Phil. Trans. R. Soc. A.380: 20210109, (2022).

6. Betzig, E. et al. Imaging intracellular fluorescent proteins at nanometer resolution. Science 313, 1642–1645, (2006).

7. Bock, H. et al. Two-color far-field fluorescence nanoscopy based on photoswitchable emitters. Appl. Phys. B 88, 161–165, (2007).

8. Wegel, E. et al. Imaging cellular structures in super-resolution with SIM, STED and Localization Microscopy: A practical comparison. Sci. Reports 6, 27290, (2016).

9. Oi, C. et al. LIVE-PAINT allows super-resolution microscopy inside living cells using reversible peptide-protein interactions. Commun. Biol. 3, 458, (2020).

10. Dertinger, T., Colyer, R., Iyer, G., Weiss, S. & Enderlein, J. Fast, background-free, 3d super-resolution optical fluctuation imaging (sofi). Proc. Natl. Acad. Sci. 106, 22287–22292, (2009).

11. Gustafsson, N. et al. Fast live-cell conventional fluorophore nanoscopy with ImageJ through super-resolution radial fluctuations. Nat. Commun. 7, 12471, (2016).

12. Agarwal, K. & Machán, R. Multiple signal classification algorithm for super-resolution fluorescence microscopy. Nat. Commun. 7, 13752, (2016).

13. Yahiatene, I., Hennig, S., Müller, M. & Huser, T. Entropy-Based Super-Resolution Imaging (ESI): From Disorder to FineDetail.ACS Photonics 2, 1049–1056, (2015).

14. Culley, S.et al. Quantitative mapping and minimization of super-resolution optical imaging artifacts. Nat. Methods 15, 263–266, (2018).

15. Takeshima, T., et al. A multi-emitter fitting algorithm for potential live cell super-resolution imaging over a wide range of molecular densities. Journal of Microscopy, 271: 266–281 (2018).

16. Carrington, W. et al. Superresolution three-dimensional images of fluorescence in cells with minimal light exposure. Science 268, 1483–1487 (1995).

17. Fukunaga, K. & Hostetler, L. The estimation of the gradient of a density function, with applications in pattern recognition. IEEE Transactions on Inf. Theory 21, 32–40, (1975).

18. Cheng Y. Mean shift, mode seeking, and clustering.IEEE Transactions on Pattern Analysis Mach. Intell. 17, 790–799, (1995).

19. Comaniciu, D. and Meer, P., Mean shift: a robust approach toward feature space analysis, IEEE Transactions on Pattern Analysis Mach. Intell. 24, 603–619 (2002).

20. Emami, E., Fathy, M. and Kozegar, E. Online failure detection and correction for camshift tracking algorithm, 2013 8th Iranian Conference on Machine Vision and Image Processing (MVIP), 180–183 (2013).

21. 21. Fazekas, F. J., Shaw, T. R., Kim, S., Bogucki, R. A. & Veatch, S. L. A mean shift algorithm for drift correction in localization microscopy. Biophys. Reports 1 (2021).

22. Rayleigh, L. On the theory of optical images, with special reference to the microscope. J. Royal Microsc. Soc. 23, 474–482, (1903).

23. Sparrow, C. On Spectroscopic Resolving Power. Astrophys. J. 44, 76, (1916).

24. Diaspro, A. & Bianchini, P. Optical nanoscopy. La Rivista del Nuovo Cimento 43, 385– 455, (2020).

25. Sharma, K. Optics: Principles and Applications (Academic Press, Elsevier Science, 2006).

26. Gustafsson, M., Agard, D., and Sedat, J. Sevenfold improvement of axial resolution in 3D wide-field microscopy using two objective-lenses, in Three-Dimensional Microscopy: Image Acquisition and Processing II, 2412, T. Wilson and C. J. Cogswell. International Society for Optics and Photonics, 147–156 (1995).

27. Guerra, J. Super-resolution through illumination by diffraction-born evanescent waves,” Appl. Phys. Lett. 66, 3555–3557 (1995).

28. Tychinsky V. & Odintsov A. New concept of optical super resolution, in Laser Dimensional Metrology: Recent Advances for Industrial Application, M. J. Downs, International Society for Optics and Photonics, 2088, 206 – 210. (1993).

29. Michalet X. & Weiss S. Using photon statistics to boost microscopy resolution,” Proc. Natl. Acad. Sci. 103, 4797–4798 (2006).

30. Terwilliger, T., Adams, P., Afonine, P., & Sobolev, O. A fully automatic method yielding initial models from high-resolution cryo-electron microscopy maps. Nat. Methods 15 (2018).

31. Hayashi T. & Tsubouchi T., Estimation and sharpening of blur in degraded images captured by a camera on a moving object. Sensors 22 (2022).

32. Gonzalez R. and Woods, R. Digital image processing (Addison-Wesley, 1992).

33. Richardson, W. Bayesian-based iterative method of image restoration J. Opt. Soc. Am. 62, 55–59 (1972).

34. Lucy, L. An iterative technique for the rectification of observed distributions. J. Astron. 79, 745–754 (1974).

35. Marsh, R. et al. Artifact-free high-density localization microscopy analysis. Nat. Methods 15, 689–692 (2018).

36. Descloux, A., Grußmayer, K. & Radenovic, A. Parameter-free image resolution estimation based on decorrelation analysis. Nat. methods 16, 918—924 (2019).

37. Huff, J. The Airyscan detector from ZEISS: confocal imaging with improved signal-to-noise ratio and super-resolution. Nat Methods 12, i–ii (2015).

38. De Luca, G. et al. Re-scan confocal microscopy: scanning twice for better resolution. Biomed. Opt. Express 4, 2644–2656 (2013).

39. Wu X., Hammer J.A. ZEISS Airyscan: Optimizing Usage for Fast, Gentle, Super-Resolution Imaging. In: Brzostowski J., Sohn H. (eds) Confocal Microscopy. Methods in Molecular Biology, 2304, (2021).

40. Tavrov, A & Tychinsky, V. Wavefront dislocations and phase image formation inside diffraction spot. Proc. SPIE 2004, Interferometry VI: Applications (1994).

41. S. Yoon, E.-H. Choi, J.-W. Kim, & K. P. Kim. Structured illumination microscopy imaging reveals localization of replication protein A between chromosome lateral elements during mammalian meiosis. Exp. & Mol. Medicine 50, 1–12 (2018).

42. 42. Dataset: Tubulin 2d high density. http://bigwww.epfl.ch/smlm/challenge2013/.

43. Sage, D. et al. Super-resolution fight club: assessment of 2D and 3D single-molecule localization microscopy software. Nat. Methods 16, 387–395, (2019).

44. Opstad, I. et al. Fluorescence fluctuation-based super-resolution microscopy using multimodal waveguided illumination,” Opt. Express 29, 23368–23380 (2021).

45. 45. Dataset: Bundled tubes of high density. http://bigwww.epfl.ch/smlm/challenge2013/

46. Boškovi A., Bender, A., Gall, L., Ziegler-Birling, C., Beaujean, N., & Torres-Padilla, M.-E. Analysis of active chromatin modifications in early mammalian embryos reveals uncoupling of h2a.z acetylation and h3k36 trimethylation from embryonic genome activation. Epigenetics 7, 747–757 (2012).

47. Schnitzbauer, J., Strauss, M. T., Schlichthaerle, T., Schueder, F. & Jungmann, R. Super-resolution microscopy with DNA-PAINT. Nat. Protoc. 12, 1198–1228, (2017).

48. Wang, H. & Milstein, J. N. Simulation Assisted Analysis of the Intrinsic Stiffness for Short DNA Molecules Imaged with Scanning Atomic Force Microscopy. PLOS ONE 10, 1–11, (2015).

49. Sebastian Acuña, Ida S. Opstad, Fred Godtliebsen, Balpreet Singh Ahluwalia, and Krishna Agarwal. Soft thresholding schemes for multiple signal classification algorithm. Opt. Express 28, 34434–34449 (2020).

50. Elson, Elliot L. Biophysical Journal, Volume 101, Issue 12, 2855-2870, (2011).

51. Reichenbach, S. & Geng, F. Two-dimensional cubic convolution. IEEE Transactions on Image Process. 12, 857–865 (2003).

52. Eddyy, W., Fitzgerald, M. & Noll, D. “Improved image registration by using fourier interpolation,” Magn. Reson. Medicine 36, 923–931 (1996).

53. Corbett, A. D.et al. Microscope calibration using laser written fluorescence. Opt. Express 26, 21887–21899, (2018).

54. Garcés Suárez, Y., et al. Nanoscale organization of rotavirus replication machineries. Elife 8, e42906, (2019).

55. Romarowski, A. et al. Super-resolution imaging of live sperm reveals dynamic changes of the actin cytoskeleton during acrosomal exocytosis. J. Cell Sci. 131, (2018).

56. Balestrini, P. A. et al. Seeing is believing: Current methods to observe sperm acrosomal exocytosis in real time. Mol. Reproduction Dev. 87, 1188–1198, (2020).

57. Cox, S. et al. Bayesian localization microscopy reveals nanoscale podosome dynamics. Nat. Methods 9, 195–200, (2012).

58. Jin, S., Haggie, P. M. & Verkman, A. S. Single particle tracking of membrane protein diffusion in a potential: Simulation, detection, and application to confined diffusion of cftr cl-channels. Biophys. J. 93, 1079–1088, (2007).

59. Yamashita, N. et al. Three-dimensional tracking of plus-tips by lattice light-sheet microscopy permits the quantification of microtubule growth trajectories within the mitotic apparatus. J. Biomed. Opt. 20, 1 – 18, (2015).

60. Chenouard, N. et al. Objective comparison of particle tracking methods. Nat. Methods 11, 281–289, (2014).

61. Jaqaman, K., Loerke, D., Mettlen, M., Kuwata, H., Grinstein, S., Schmid, S. L. and Danuser, G. Robust single-particle tracking in live-cell time-lapse sequences, Nat. Methods 5, 695 (2008).

62. Fields, A. P. and Cohen, A. E. Optimal tracking of a brownian particle, Opt. Express 20, 22585–22601 (2012).

63. Kalman, R. E. A New Approach to Linear Filtering and Prediction Problems, J. Basic Eng.82, 35–45 (1960).

64. Yüce M. Y., Erdoğan, A., Jonáš, A. & Kiraz, A. Single molecule tracking with kalman filtering. Frontiers in Optics 2011/Laser Science XXVII, FTuH5, (Optical Society of America, 2011).

65. Tinevez, J.-Y., Perry, N., Schindelin, J., Hoopes, G. M., Reynolds, G. D., Laplantine, E., Bednarek, S. Y., Shorte, S. L. and Eliceiri, K. W. Trackmate: An open and extensible platform for single-particle tracking, Methods 115, 80–90 (2017).

66. Patel, M., Leggett, S. E., Landauer, A. K., Wong, I. Y., and Franck, C. Rapid, topology-based particle tracking for high-resolution measurements of large complex 3D motion fields, Sci. Reports 8, 5581 (2018).

67. Gross, J., Köster, M. & Krüger, A. Fast and efficient nearest neighbor search for particle simulations. Computer Graphics and Visual Computing CGVC, 55–63, (The Eurographics Association, 2019).

68. Cheezum, M. K., Walker, W. F., and Guilford, W. H. Quantitative Comparison of Algorithms for Tracking Single Fluorescent Particles, Biophys. J. 81, 2378–2388 (2001).

69. Calcines-Cruz, C., Finkelstein, I. J. & Hernandez-Garcia, A. Crispr-guided programmable self-assembly of artificial virus-like nucleocapsids. Nano Lett. 21, 2752–2757, (2021).

70. Federici, F., Dupuy, L., Laplaze, L., Heisler, M. & Haseloff, J. Integrated genetic and computation methods for *in planta* cytometry. Nat. Methods 9, 483–485, (2012).

71. Torres-Martínez, H. H., Hernández-Herrera, P., Corkidi, G. & Dubrovsky, J. G. From one cell to many: Morphogenetic field of lateral root founder cells in *arabidopsis thaliana* is built by gradual recruitment. Proc. Natl. Acad. Sci. 117, 20943–20949, (2020).

72. Ricci, M. A., Manzo, C., García-Parajo, M. F., Lakadamyali, M., & Cosma, M. P. (2015). Chromatin fibers are formed by heterogeneous groups of nucleosomes in vivo. Cell, 160, 1145–1158.

73. Rutowicz, K., Lirski, M., Mermaz, B., Teano, G., Schubert, J., Mestiri, I., Kroten, M. A., Fabrice,T. N., Fritz, S., Grob, S., Ringli, C., Cherkezyan, L., Barneche, F., Jerzmanowski, A., and Baroux, C. Linker histones are fine-scale chromatin architects modulating developmental decisions in arabidopsis, Genome Biol. 20, 157 (2019).

74. Wu, G., Zhao, X., Luo, S. & Shi, H. Histological image segmentation using fast mean shift clustering method. BioMedical Eng. OnLine 14, 24, (2015).

75. Szeliski, R. Computer vision: Algorithms and applications (Springer Science & Business Media, 2011).

76. Sankaran, J. et al. Simultaneous spatiotemporal super-resolution and multi-parametric fluorescence microscopy. Nat. Commun. 12, 1748, (2021).

77. Lin, Y.-H., Chang, W.-L., and Hsieh, C.-L. Shot-noise limited localization of single 20 nm gold particles with nanometer spatial precision within microseconds, Opt. Express 22, 9159–9170 (2014).

78. Glover, Z. J., Ersch, C., Andersen, U., Holmes, M. J., Povey, M. J., Brewer, J. R. and Simonsen, A. C. Super-resolution microscopy and empirically validated autocorrelation image analysis discriminates microstructures of dairy derived gels,” Food Hydrocoll. 90, 62–71(2019).

79. Betzig, E. Nobel lecture: Single molecules, cells, and super-resolution optics, Rev. Mod. Phys. 87, 1153–1168 (2015).

80. Liss, V., Barlag, B., Nietschke, M. and Hensel, M. Self-labelling enzymes as universal tags for fluorescence microscopy, super-resolution microscopy and electron microscopy, Sci. Reports 5, 17740 (2015).

81. de Boer, P., Hoogenboom, J. P. & Giepmans, B. N. G. Correlated light and electron microscopy: ultrastructure lights up! Nat. Methods 12, 503–513, (2015).

82. Cahoon, C. K., et al. Superresolution expansion microscopy reveals the three-dimensional organization of the Drosophila synaptonemal complex. Proc. Natl. Acad. Sci. 114, 6857–6866, (2017).

83. Kim, J., Wojcik, M., Wang, Y., Moon, S., Zin, E.A., Marnani, N., Newman, Z.L., Flannery, J.G., Xu, K. and Zhang, X. Oblique-plane single-molecule localization microscopy for tissues and small intact animals, Nat. Methods 16, 853–857 (2019).

84. Hu, Y. S., Zimmerley, M., Li, Y., Watters, R. & Cang, H. Single-molecule super-resolution light-sheet microscopy,” ChemPhysChem 15, 577–586 (2014).

85. Chen, R., et al. Deep-learning super-resolution microscopy reveals nanometer-scale intracellular dynamics at the millisecond temporal resolution, Preprint at https://doi.org/10.1101/2021.10.08.463746 (2021).

86. Puschmann, K. G. & Kneer, F. On super-resolution in astronomical imaging. Astronomy & Astrophysics, 436, 373–378, (2005).

87. Guo, R., Shi, X., Zhu, Y. & Yu, T. Super-resolution reconstruction of astronomical images using time-scale adaptive normalized convolution. Chin. J. Aeronaut, 31, 1752–1763, (2018).

