## Supplementary material for "Nanoscopic resolution within a single imaging frame": MSSR Supplementary Information

### Nanoscopic resolution within a single imaging frame
### Supplementary information

#### CONTENTS

**0 Symbols and notation** **2**

**1 Supplementary Note 1. Microscope Imaging Model** **4**

**2 Supplementary Note 2. MeanShift Theory** **5**

**3 Supplementary Note 3. Meanshift Imaging Model** **7**

**4 Supplementary Note 4. MeanShift local minimum over a Gaussian distribution. Theorem** **10**

**5 Supplementary Note 5. MeanShift Super-Resolution algorithm** **14**

**7 Supplementary Note 7. MSSR processing time comparison for different analysis platforms** **33**

**8 Supplementary Note 8. Single-frame nanoscopy free of noise-dependent artifacts** **35**

**9 Supplementary Note 9. Comparing MSSR with other analytical SRM methods** **37**

**10 Supplementary Note 10. Empirical demonstration of MSSR applicability to a wide range of fluorescence bioimaging modalities** **41**

**11 Applicability of MSSR to 3D microscopy** **53**

**12 Extended supplementary figures** **54**

**13 Extended supplementary movies** **56**

### 0. SYMBOLS AND NOTATION

| notation | Description |
|---|---|
| | Microscope imaging |
| $PSF$ | general term of point spread function |
| $PSF_B$ | Airy pattern or point spread function using a Bessel distribution |
| $PSF_G$ | point spread function using a Gaussian distribution |
| $NA$ | numerical aperture |
| $\lambda_{em}$ | emission wavelength of a given fluorophore |
| $n$ | refractive index of the medium |
| $w$ | pixel size |
| $b(x)$ | binary representation of the collected signal |
| $e_m$ | emission power of emitter $m$ |
| $\Delta t$ | acquisition time between frames |
| $c_m$ | total emission of emitter $m$ in the period of time $\Delta t$ |

| | MeanShift theory |
|---|---|
| $K, k$ | kernel, profile |
| $\mathbf{K}, \mathbf{k}$ | spatial-range kernel, spatial-range profile respectively |
| $I, \mathbf{I}$ | image, image stack respectively |
| $M$ | number of frames of $\mathbf{I}$ |
| $h, \mathbf{h}$ | bandwidth, spatial-range bandwidth respectively |
| $B^*_{\mathbf{h}}(p_0)$ | neighborhood of radius h centered and punctured at $p_0$ |
| $h_s, h_r$ | spatial bandwidth, range bandwidth respectively |
| $p_0$ | central pixel of the $B^*_h(p_0)$ |
| $p_n$ | neighboring pixel |
| $x_0, y_0$ | coordinates and intensity of $p_0$ respectively |
| $x_n, y_n$ | coordinates and intensity of $p_n$ respectively |
| $q_n$ | weighted coefficients associated to $y_n$. |
| $f$ | unknown probability density function |
| $\widehat{f}$ | kernel density estimator function |
| $N, n$ | number of elements in $B^*_h(y)$, element number in $B^*_h(y)$ respectively |
| $SM$ | sample mean |
| $MS$ | meanshift vector |
| $t_s, t_r$ | arguments related to the spatial and range components respectively |

| $MSSR$ algorithm | |
|---|---|
| $MSSR$ | MeanShift Super-Resolution algorithm |
| $MSSR^n$ | MeanShift Super-Resolution of order $n$ |
| $sf - MSSR^n$ | single frame MeanShift Super-Resolution of order $n$ |
| $t - MSSR^n$ | temporal MeanShift Super-Resolution of order $n$ |
| $FWHM$ | full width at half maximum |
| $AMP$ | image magnification |
| $FWHM$ of $PSF$ | is directly related to the kernel window size $h_s$, $h_s = FWHM/2$ |
| $order$ | $MSSR$ iteration |
| $PTF$ | Pixel-wise Temporal Function |
| $Mean$ | Pixel-wise Temporal Average |
| $Var$ | Pixel-wise Temporal Variance |
| $TPM$ | Pixel-wise Temporal Product Mean |
| $CV$ | Pixel-wise Temporal Coefficient of Variation |
| $SOFI_2$ | Auto-cumulant Temporal of order 2 |
| $SOFI_3$ | Auto-cumulant Temporal of order 3 |
| $SOFI_4$ | Auto-cumulant Temporal of order 4 |
| $RSP$ | Resolution Scaled Pearson |
| $RSE$ | Resolution Scaled Error |

#### 1. SUPPLEMENTARY NOTE 1. MICROSCOPE IMAGING MODEL

In imaging microscopy, light emitters whose dimensions are within the order of the emission wavelength are considered as point sources which are affected by the diffraction of light. This effect is modeled by convolution of the fluorescence distribution of the emitter with the point spread function ($PSF$) of the optical system [1]:

$$PSF_B = \left(\frac{2J_1(\nu)}{\nu}\right)^2 \quad \text{(S1)}$$

where $J_1$ is the Bessel function of the first kind of order 1, $\nu = \frac{2\pi \mathrm{NA} r}{\lambda n}$ is a dimensionless distance from the optical axis in the focal plane, $NA$ is the numerical aperture of the objective, $r$ is the radial distance, $\lambda$ is the emission wavelength of the emitter and $n$ is the refractive index of the medium used in the objective of the microscope [2].

The result of convolving the $PSF$ with a single emitter is an Airy pattern in which approximately 86% of the emitted light is harbored within the central disk. Concentric rings are arranged around the central disk whose intensity decreases as a function of the distance from the center. The location of the minimum of the central disk with respect to its central point is given at a distance, this is define as the $PSF$ parameter denoted by $r_{min}$ and is calculated as:

$$r_{min} = \frac{0.61\lambda}{\mathrm{NA}} \quad \text{(S2)}$$

Optical considerations suggest a robust computation of the $NA$ by the expression

$$NA = \frac{n_{medium}}{n_{oil}} NA_{obj} \quad \text{(S3)}$$

The light emanating from the image plane is captured and digitized as gray levels within pixels represented as a regular 2-dimensional lattice $I$. Pixels are assumed to be squares of width $w$. The radius of an emitter collected at the image plane is characterized by

$$r_{min}(w) = \frac{0.61\lambda}{\mathrm{NA}w} \quad \text{(S4)}$$

Pixels located within this region hold the largest amount of information of the emitter's location. On the other hand, pixels beyond this region commonly correspond to the background noise of the image.

The $PSF_B$ can also be approximated by a Gaussian function of the form:

$$PSF_G = e^{-\frac{1}{2}\frac{x^2+y^2}{\sigma^2}}, \quad \text{(S5)}$$

where $x$ and $y$ are spatial coordinates and $\sigma$ is the width along the x and y-axis [3].

Let $\Delta t$ be the acquisition time between frames and $M$ the number of emitters. The emission from a single blinking emitter $e$ is given by:

$$c_m = e_m \int_0^{\Delta t} b(x)dx, \quad \text{(S6)}$$

where $e_m$ is the emission power of the emitter $m$ and $b(x)$ is a binary representation of the collected signal. Let $p$ be a pixel with its center located at $x$ in the image plane, the contribution at $p$ of all $M$ emitters is given by:

$$I(p) = \sum_{m=1}^{M} PSF_B(x) c_m w^2 \quad \text{(S7)}$$

or

$$I(p) = \sum_{m=1}^{M} PSF_G(x) c_m w^2. \quad \text{(S8)}$$

Note that $PSF_B(x)$ indicates a first-order Bessel approximation of the $PSF$ and $PSF_G(x)$ indicates a Gaussian approximation of the $PSF$. In the following sections, we will assume that images are affected by either of these two types of $PSF$ representations.

#### 2. SUPPLEMENTARY NOTE 2. MEANSHIFT THEORY

Let $Y = \{y_n\}_{n=1}^{N}$ be a finite set of data points in the $d$-dimensional Euclidean space $\mathbb{R}^d$. The unknown density function $f$ at a given point $y_0 \in \mathbb{R}^d$ is described by a kernel density estimator $\widehat{f}$ based on the symmetric radial kernel $K$ and scalar parameter $h > 0$ (bandwidth) by [4–6]:

$$\widehat{f}_{K,h}(y_0) = \frac{c_K}{Nh^d} \sum_{y_n \in B_h^*(y_0)} K\left(\frac{y_0 - y_n}{h}\right), \tag{S9}$$

where $B_h^*(y_0)$ is the punctured neighborhood at $y_0$ of radius $h$, $N$ is the number of points, the term $h^d$ is an scaled factor proportional to $h$ and the number of dimensions, and $c_K$ is a normalization constant, so $\widehat{f} \in [0,1]$. According to Cheng [5], a function $K : \mathbb{R}^d \to R$ is a kernel if there exists a profile $k : [0,\infty) \to \mathbb{R}$, such that $K(x) = k(\|x\|^2)$, where $\|\cdot\|$ is the Euclidean norm. $k$ satisfies the following conditions: non-negative, non-increasing, differenciable, integrable and piecewise-defined. By using profile notation, formula S9 is written as:

$$\widehat{f}_{k,h}(y_0) = \frac{c_k}{Nh^d} \sum_{y_n \in B_h^*(y_0)} k\left(\left\|\frac{y_0 - y_n}{h}\right\|^2\right). \tag{S10}$$

Examples of kernels and properties are described in [5, 7]. For the rest of the document we will use the profile notation.

As the value of $\widehat{f}$ decreases, the heterogeneity of $B_h^*(y_0)$ increases, which means that the difference between the elements of the analyzed region increases. If $Y \cap B_h^*(y_0) = \emptyset$ means that there are not points in $B_h^*(y_0)$, in this case $f(y_0) = 0$.

The estimated gradient density of $f$ at $y_0$ is given by:

$$\begin{aligned}
\widehat{\nabla f}_{k,h}(y_0) &= \nabla \widehat{f}_{k,h}(y_0) \\
&= \frac{2c_k}{Nh^d} \sum_{y_n \in B_h^*(y_0)} \frac{y_0 - y_n}{h^2}\, k'\left(\left\|\frac{y_0 - y_n}{h}\right\|^2\right) \\
&= \frac{2c_k}{Nh^{d+2}} \sum_{y_n \in B_h^*(y_0)} (y_0 - y_n) \left[-g\left(\left\|\frac{y_0 - y_n}{h}\right\|^2\right)\right] \\
&= \frac{2c_k}{Nh^{d+2}} \sum_n (y_n - y_0)\, g_n,
\end{aligned} \tag{S11}$$

where $k'(\cdot)$ is the derivative of $k$, $g(\cdot) = -k'(\cdot)$ and $g_n = g\left(\left\|\frac{y_0 - y_n}{h}\right\|^2\right)$.

Applying algebraic transformations, the gradient density estimator can be written as:

$$\widehat{\nabla f}_{k,h}(y_0) = \frac{2c_k}{Nh^{d+2}} \cdot \sum_n g_n \cdot \left[\frac{\sum_n g_n y_n}{\sum_n g_n} - y_0\right]. \tag{S12}$$

Note that the first factor in equation S12 is a constant. The second factor is proportional to the kernel density estimator by using profile $g$ and bandwidth $h$:

$$\widehat{f}_{g,h}(y_0) = \frac{c_g}{Nh^d} \sum_n g_n. \tag{S13}$$

Finally the last term in brackets is the mean shift vector with profile $g$ and bandwidth $h$:

$$MS_{g,h} = \frac{\sum_n g_n y_n}{\sum_n g_n} - y_0, \tag{S14}$$

it refers to the difference between the weighted mean or sample mean ($SM$) of all $y_n$ that belong to $B_h^*(y_0)$ and $y_0$.

Isolating $MS$ in formula S12 (last term in brackets), we obtain that $MS$ always points towards the same direction of the gradient density using profile $k$ as shown in equation S15.

$$MS_{g,h}(y_0) = h^2 \frac{c_g}{c_k} \frac{\widehat{\nabla f}_{k,h}(y_0)}{\widehat{f}_{g,h}(y_0)}. \tag{S15}$$

The profiles $k$ and $g$ are related by the *shadow kernel* definition, also given in [5]. Let $K, G$ be kernels with profiles $k, g$ respectively. Then $K$ is a shadow kernel of $G$ if the meanshift using $G$ is in the gradient direction of the density estimation using $K$. As a direct consequence of that, $k$ is a shadow profile of $g$ and they must satisfy that

$$k'(t) = -c \cdot g(t), \tag{S16}$$

where $c$ is a scalar positive constant. Supplementary Table S1 shows examples of two popular profiles and their respective shadows. Note that the Epanechnikov profile is a linear function and its shadow is a constant, however, the Gaussian profile and its shadows are the same being both an exponential function. See [5] for other types of profiles and shadows.

| Name | Kernel $k(\|u\|^2)$ | Profile $k(t)$ | Derivative $k^{'}(t)$ | $g(t)$ | Shadow |
|---|---|---|---|---|---|
| Epanechnikov | $1 - \|u\|^2$ | $1 - t$ | $-1$ | 1 | Uniform |
| Gaussian | $e^{-\frac{1}{2}\|u\|^2}$ | $e^{-at}$ | $-ae^{-at}$ | $e^{-at}$ | Gaussian |

**Table S1.** Epanechnikov and Gaussian profiles, derivatives and their corresponding shadows.

$MS_{g,h}$ at $y_0$ measures the homogeneity of $B_h^*(y_0)$. Points of minimum local density are assigned to large values of $MS$. For this reason the *Meanshift Theory* was designed around the idea of moving away from minimum local density points and converges towards higher density regions. The process repeats until convergence or until a fixed point is reached.

In this work, the *Meanshift Theory* is not implemented in a classical fashion. The concept of the shadow kernel is crucial for the understanding the MS and its derivation, however, this concept has not been formalized rigorously for digital images. Here, we developed an extension of the definition of the shadow kernel given by Cheng to the imaging field over the spatial-range domain (Supplementary Note 3). Our model was then evaluated for a Gaussian distribution (Supplementary Note 4) and finally adjusted to fit super-resolution microscopy theory, which gives rise to $MSSR$ (Supplementary Note 5).

#### 3. SUPPLEMENTARY NOTE 3. MEANSHIFT IMAGING MODEL

The characterization of $MS$ in images is defined in terms of two pixel properties: position and intensity. A digital image $I$ is frequently represented as a $m$-dimensional lattice (coordinates or position) of $d$-dimensional vectors (color or intensity). The spatial-range neighborhood is a suitable tool to compute the kernel values simultaneously throughout the location and intensity pixels. In this way, it is convenient to represent pixel components as $p = [x, y(x)]$, where $x$ represents the coordinate component and $y(x)$ represents the color component at $x$. Therefore, a digital image is represented by $I = \{p_n\}_{n=1}^{M}$, where $M$ is the number of pixels.

Let $p_0 = [x_0, y(x_0)]$ be a pixel, $\mathbf{h} = (h_s, h_r)$ a spatial-range bandwidth (a scalar bandwidth for each component: a spatial $h_s$ bandwidth and an intensity range $h_r$ bandwidth). The spatial-range neighborhood $B^*_{\mathbf{h}}(p_0)$ is determined by $p_0$ and $\mathbf{h}$, the symbol $^*$ indicates that the neighborhood excludes only the host pixel. The spatial-range profile $\mathbf{k}$ at $p_0$ is given by the product of the spatial and range profiles by using the common profile $k$ (Supplementary Note 2). The following equations define the spatial-range neighborhood, profile, and probability density function at $p_0$, respectively:

$$B^*_{\mathbf{h}}(p_0) = \{p = [x, y] : 0 < \|x_0 - x\| \leq h_s,\ 0 \leq \|y(x_0) - y\| \leq h_r\}\,, \tag{S17}$$

$$\mathbf{k_h}(p_0, p_n) = k_s \cdot k_r = k\left(\left|\left|\frac{x_0 - x_n}{h_s}\right|\right|^2\right) \cdot k\left(\left|\left|\frac{y(x_0) - y_n}{h_r}\right|\right|^2\right),\ p_n \in B^*_{\mathbf{h}}(p_0), \tag{S18}$$

$$\widehat{f}(p_0) = \frac{1}{N h_s^d h_r^m} \sum_{p_n \in B^*_{\mathbf{h}}(p_0)} \mathbf{k_h}\left(p_0, p_n\right). \tag{S19}$$

From equation S18 we can deduce the profile that will be used to calculate the $MS$ vector over a spatial-range space. We need to define a function that lies in the gradient direction of both spaces, circumscribed by $h_s$ and $h_r$, simultaneously. Note that the profile $\mathbf{k}$ is implicitly dependent on the spatial and range components, but profiles $k_s$ and $k_r$ are independent by argument, as in equation S20:

$$\mathbf{k} = k_s(t_s) \cdot k_r(t_r), \tag{S20}$$

where

$$t_s = \left|\left|\frac{x_0 - x_n}{h_s}\right|\right|^2 \tag{S21}$$

and

$$t_r = \left|\left|\frac{y_0 - y_n}{h_r}\right|\right|^2 \tag{S22}$$

are the arguments related to the spatial and range components respectively.

Note that, for a given neighborhood $B^*_{\mathbf{h}}(p_0)$, $p_0$ and $h$ are fixed. This means that in expressions S21 and S22, $x_0$, $h_s$, $y_0$ and $h_r$ are also fixed, but their values vary according the position $x_n$ and $y_n$ respectively. In particular, the term $t_s$ depends on the euclidean distance between the spatial coordinates of $x_0$ and $x_n$. Consequently, $k_s$ varies along the whole neighborhood according the distance between the location of this pixel as shown in Figure S1. As the position moves away from the central pixel, the distance increases (Figure S1a) and therefore the value of $k_s$ decreases (Figure S1b). Similarly it happens with $k_r$.

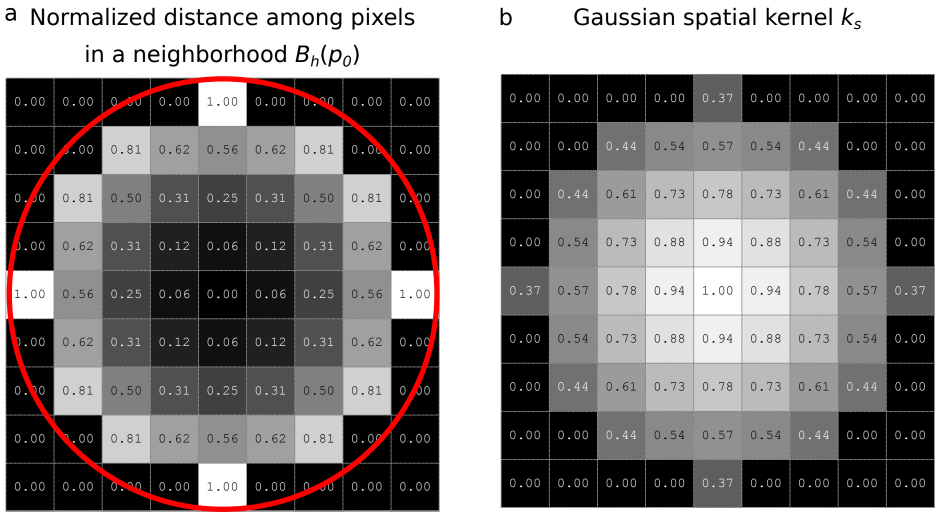

**Figure S1. Behavior of $k_s$ according the distance in a neighborhood. a)** Normalized distance of pixels in a neighborhood $B_h(p_0)$ related to the central pixel. The neighborhood has 5 pixels of radius and its limits are marked by the continuous red line. As the position moves away from the central pixel, the distance increases. Pixels beyond the red line are not considered on the neighborhood and the distance is set to zero value. **b)** $k_s$ calculated from a). As the position moves away from the central pixel, values for $k_s$ the distance decreases. Pixels beyond the neighborhood are set to zero value.

Partial derivatives of $\mathbf{k}$ respect to $t_s$ and $t_r$ are given by:

$$\frac{\partial \mathbf{k}}{\partial t_s} = k_r \cdot k_s' = -k_r \cdot g_s \quad \text{and} \quad \frac{\partial \mathbf{k}}{\partial t_r} = k_s \cdot k_r' = -k_s \cdot g_r, \tag{S23}$$

where $g_s = -k_s'$ and $g_r = -k_r'$. In both equations the first factor is a negative constant with respect to the derivative variable. Defining

$$\mathbf{g} = g_s \cdot g_r = k_s' \cdot k_r', \tag{S24}$$

the condition given in equation S16 is satisfied by $\mathbf{g}$ for both components $g_s$ and $g_r$.

**Definition 1.** *Let $g_s$ and $g_r$ be two profiles with their respective shadow kernels $k_s$ and $k_r$. Then $\mathbf{k} = k_s \cdot k_r$ is a shadow kernel of $\mathbf{g} = g_s \cdot g_r$.*

The generalization of the previous definition to a product of $n$ profiles is straightforward, as long as the profiles are functions of independent arguments.

Looking back to notation used in equation S18, the equation S24 can be rewritten as:

$$\mathbf{g_h}(p_0, p_n) = g_s \cdot g_r = g\left(\left\|\frac{x_0 - x_n}{h_s}\right\|^2\right) \cdot g\left(\left\|\frac{y(x_0) - y_n}{h_r}\right\|^2\right), \; p_n \in B^*_{\mathbf{h}}(p_0). \tag{S25}$$

Now, the *MS* vector in a spatial-range space can be formulated as:

$$MS_h(p_0) = \frac{\sum_n \mathbf{g_h}(p_0, p_n) y_n}{\sum_n \mathbf{g_h}(p_0, p_n)} - y(x_0). \tag{S26}$$

Note that $MS_{\mathbf{h}}(p_0)$ is only used to compute the intensity at $p_0$ on the range space, which harbors the fluorescence intensity at the coordinates of $p_0$.

The first term in equation S26 will be denoted by $SM_{\mathbf{h}}$ and it is the sample mean (weighted local mean) in the spatial-range space:

$$SM_h(p_0) = \frac{\sum_n \mathbf{g_h}(p_0, p_n) y_n}{\sum_n \mathbf{g_h}(p_0, p_n)} = \sum_n \mathbf{q}_n y_n. \tag{S27}$$

where

$$\mathbf{q}_n = \frac{\mathbf{g_h}(p_0, p_n)}{\sum_n \mathbf{g_h}(p_0, p_n)}. \tag{S28}$$

are coefficients associated to each $y_n$. Note that, independently of how large $q_n$ is, all $\mathbf{q}_n$ lie in the interval $(0, 1]$ and $\sum_n \mathbf{q}_n = 1$. For this reason, $\mathbf{SM_h}(p_0)$ is a convex linear combination of all $y_n \in B^*_{\mathbf{h}}$ and also $\mathbf{SM_h}(p_0) \in B_{\mathbf{h}}$.

If $\mathbf{h}$ has been predefined based on the physical properties of an optical system (Supplementary Note 5), the subindex $\mathbf{h}$ is removed. The notation for applying $MS$ on every pixel of the image $I$ is $MS(I)$. Similar notation will be used for sample mean as $SM(I)$. Now, the following expressions are equivalent:

$$MS(I) = SM(I) - I = \sum_n \mathbf{q}_n y_n - I \tag{S29}$$

At this point, we have presented $MS$ from a digital image point of view. The following section contains a theorem about the localization of a Gaussian distribution maximum using **MS**. Sections 5 and 6 describe the $MSSR$ algorithm of order 0 and higher orders.

#### 4. SUPPLEMENTARY NOTE 4. MEANSHIFT LOCAL MINIMUM OVER A GAUSSIAN DISTRIBUTION. THEOREM

In this section, the following theorem has been proved for a Gaussian function over a $d$-dimensional space. In the context of microscopy, this space is restricted to two dimensions. The point of maximum value represents the center of emitters and the Gaussian values simulate the intensities. This principle applies beyond the microscopy field and is relevant in any imaging-related area that involves the use of lenses, which impinge upon the diffractive nature of light (*e.g.* astronomy).

**Theorem 1.** *Let $G$ be a Gaussian distribution and $p_0$ the point of maximum value, then $MS(p_0)$ is a local minimum.*

*Proof.* The general idea of this proof is divided into two steps. The first step consists of showing that $p_0$ is a stationary point by demonstrating that the gradient of $MS$ is always a zero vector. Second, the hessian matrix of $MS$ at this point is demonstrated to be positive-defined.

Without loss of generality, lets assume that $G$ has mean 0, variance $\sigma$ in a $d$-dimensional space and the point of maximum value is located at the origin $x_0$ (center of the image). A digital representation of $G$ is given by $p = [x, y(x)]$, where $x = (x_1, \cdots, x_d)$ and $y = e^{-\frac{1}{2}\sum_{l=1}^{d}\frac{x_l^2}{\sigma^2}}$.

Given a neighborhood $B^*(p_0)$, the elements on this region are symmetrically distributed around $p_0$ and are represented by

$$p_{x_n} = [x_0 + \Delta x_n, y(x_0 + \Delta x_n)] = [x_0 + \Delta x_n, y_n], \tag{S30}$$

and

$$\Delta y_n = y(x_0) - y_n, \tag{S31}$$

here $\Delta x_n$ is the shift of the $n$-th elements of $x$.

Expressions for the Gaussian function and its value in $p_{x_n}$ are $y = e^{-\frac{1}{2}\sum_{l=1}^{d}\frac{x_l^2}{\sigma^2}}$ and $y_n = e^{-\frac{1}{2}\sum_{l=1}^{d}\frac{(x_l+\Delta x_{n_l})^2}{\sigma^2}}$, respectively.

The gradient of $MS$ is given by:

$$\nabla MS = \left(\frac{\partial MS}{\partial x_1}, \ldots, \frac{\partial MS}{\partial x_l}, \ldots, \frac{\partial MS}{\partial x_d}\right), \tag{S32}$$

$$\frac{\partial MS}{\partial x_l} = \sum_n \mathbf{q}_n \frac{\partial y_n}{\partial x_l} - \frac{\partial y}{\partial x_l} = -\sum_n \mathbf{q}_n e^{-\frac{1}{2}\sum_{l=1}^{d}\frac{(x_l+\Delta x_{n_l})^2}{\sigma^2}} \frac{x_l + \Delta x_{n_l}}{\sigma^2} + e^{-\frac{1}{2}\sum_{l=1}^{d}\frac{x_l^2}{\sigma^2}} \frac{x_l}{\sigma^2}, \tag{S33}$$

where $l$ is the index along each spatial component and $d$ is the total number of spatial components.

Evaluation of $\nabla \mathbf{MS}$ at $p_0$ is equivalent to assign $x = 0$ on each component and due to the parity of $y$ and the $\Delta x_n$ they are distributed symmetrically in the $B^*(p_0)$. Then

$$\begin{aligned} \frac{\partial MS}{\partial x_l}(p_0) &= -\sum_n \mathbf{q}_n e^{-\sum_{l=1}^{d}\frac{(\Delta x_{n_l})^2}{\sigma^2}} \frac{\Delta x_{n_l}}{\sigma^2} \\ &= -\left[\sum_{\Delta x_{n_l}} \mathbf{q}_n e^{-\frac{1}{2}\sum_{l=1}^{d}\frac{(\Delta x_{n_l})^2}{\sigma^2}} \frac{\Delta x_{n_l}}{\sigma^2} + \sum_{\Delta x'_{n_l}} \mathbf{q}_n e^{-\frac{1}{2}\sum_{l=1}^{d}\frac{(\Delta x'_{n_l})^2}{\sigma^2}} \frac{\Delta x'_n}{\sigma^2}\right]. \end{aligned} \tag{S34}$$

In this way, for every $x_n$ there exists a symmetric element $x'_n$ such as

$$\Delta x_n + \Delta x'_n = \mathbf{0} \text{ and } \Delta y_n = \Delta y_n\prime. \tag{S35}$$

Also, note that $\mathbf{q}_n$ (defined on equation S28 in previous section) only depends on $\Delta y_n$ and, given their symmetry, they satisfy

$$\mathbf{q}_n(\Delta y_n) = \mathbf{q}_n(\Delta y_n\prime). \tag{S36}$$

Supplementary Figure S2 shows the relationship of two points $x_n$ and $x'_n$ symmetrically distributed within a neighborhood of center $x_0$.

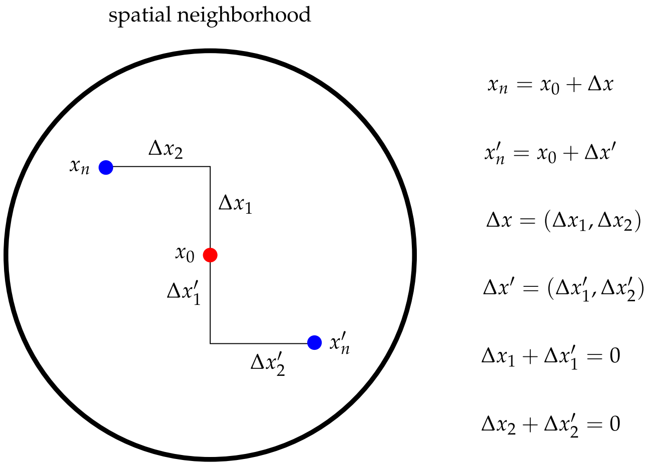

**Figure S2. Spatial distribution of symmetrical points around the center of a spatial neighborhood that cancel each other out by addition.** Point in red $x_0$ is the center of the neighborhood, points in blue $x_n$ and $x'_n$ shape a symmetric spatial distribution of two elements of the neighborhood. $\Delta x_n$ and $\Delta x'_n$ are the shifts of $x_n$ and $x'_n$ from $x_0$ respectively. Note that the sum of the shifts is zero: $\Delta x_1 + \Delta x'_1 = 0$, $\Delta x_2 + \Delta x'_2 = 0$.

$$\begin{aligned} \frac{\partial MS}{\partial x_l}(p_0) &= -\sum_{\Delta x_{n_l}} \mathbf{q}_n e^{-\frac{1}{2}\sum_{l=1}^{d}\frac{(\Delta x_{n_l})^2}{\sigma^2}} \frac{\Delta x_{n_l}}{\sigma^2} + \sum_{\Delta x'_{n_l}} \mathbf{q}_n e^{-\frac{1}{2}\sum_{l=1}^{d}\frac{(\Delta x'_{n_l})^2}{\sigma^2}} \frac{\Delta x_{n_l})'}{\sigma^2} \\ &= 0 \end{aligned} \tag{S37}$$

This holds to be true for every component $MS_h$, so $\nabla MS = \mathbf{0}$. Therefore, $p_0$ is a point of local extreme of $MS$.

Now we will prove that $p_0$ is a minimum of $MS_h$.

The Hessian matrix of $MS$ is given by the following expression:

$$H(MS) = \left[\frac{\partial^2 MS}{\partial x_i \partial x_j}\right]_{i,j=\overline{1,d}} \tag{S38}$$

The elements out of the diagonal of the Hessian matrix are given by:

$$\left(\frac{\partial^2 MS}{\partial x_i \partial x_j}\right)_{i\neq j} = \sum_n \mathbf{q}_n e^{-\frac{1}{2}\sum_{l=1}^{d}\frac{(x_{n_l}+\Delta x_{n_l})^2}{\sigma^2}} \left(-\frac{x_{n_i}+\Delta x_{n_i}}{\sigma^2}\right)\left(-\frac{x_{n_j}+\Delta x_{n_j}}{\sigma^2}\right) \tag{S39}$$

Evaluating the previous equation in $p_0$, we obtain:

$$\left(\frac{\partial^2 MS}{\partial x_i \partial x_j}\right)_{i\neq j}(p_0) = \sum_n \mathbf{q}_n e^{-\frac{1}{2}\sum_{l=1}^{d}\frac{(\Delta x_{n_l})^2}{\sigma^2}} \frac{\Delta x_{n_i}\Delta x_{n_j}}{\sigma^4} \tag{S40}$$

Once again, given the symmetry of $B^*(p_0)$ for every $\Delta x_{n_i}$ and $\Delta x_{n_j}$ there exist $\Delta x'_{n_i}$ and $\Delta x'_{n_j}$ such as (Supplementary Figure S3):

- $\Delta x_{n_i}\Delta x_{n_j} + \Delta x'_{n_i}\Delta x'_{n_j} = 0$,
- $\sum_{l=1}^{d}(\Delta x_{n_l})^2 = \sum_{l=1}^{d}(\Delta x'_{n_l})^2$,
- $\mathbf{q}_n(\Delta y_n) = \mathbf{q}_n(\Delta y_n\prime)$.

So,

$$\left(\frac{\partial^2 MS}{\partial x_i \partial x_j}\right)_{i\neq j}(p_0) = \sum_{\Delta x_{n_i}, \Delta x_{n_j}} \mathbf{q}_n e^{-\frac{1}{2}\sum_{l=1}^{d}\frac{(\Delta x_{n_l})^2}{\sigma^2}} \frac{\Delta x_{n_i}\Delta x_{n_j}}{\sigma^4} + \sum_{\Delta x'_{n_i}, \Delta x'_{n_j}} \mathbf{q}_n e^{-\frac{1}{2}\sum_{l=1}^{d}\frac{(\Delta x'_{n_l})^2}{\sigma^2}} \frac{\Delta x'_{n_i}\Delta x'_{n_j}}{\sigma^4} = 0 \tag{S41}$$

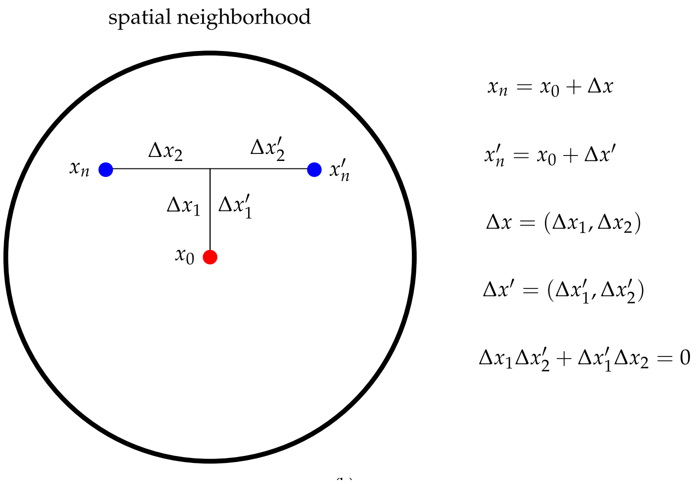

**Figure S3. Spatial distribution of symmetrical points around the center of a spatial neighborhood that cancel each other out by product and addition.** The red point $x_0$ is the center of the neighborhood, points in blue $x_n$ and $x'_n$ represents a symmetrical spatial distribution of two elements of the neighborhood. $\Delta x_n$ and $\Delta x'_n$ are the shift of $x_n$ and $x'_n$ from the $x_0$ respectively. Note that the sum of the shift product by coordinates is zero: $\Delta x_1 \Delta x'_2 + \Delta x'_1 \Delta x_2 = 0$.

The elements in the diagonal of the Hessian matrix are given by:

$$\frac{\partial^2 MS}{\partial x_i^2} = \sum_n \mathbf{q}_n e^{-\frac{1}{2}\sum_{l=1}^{d}\frac{(x_{n_i}+\Delta x_{n_l})^2}{\sigma^2}} \frac{(x_{n_i}+\Delta x_{n_i})^2 - \sigma^2}{\sigma^4} - e^{-\frac{1}{2}\sum_{l=1}^{d}\frac{(x_{n_l})^2}{\sigma^2}} \frac{x_{n_i}^2 - \sigma^2}{\sigma^4}. \tag{S42}$$

Evaluating $p_0$ in equation S42, we obtain:

$$\begin{aligned}\frac{\partial^2 MS_{\mathbf{h}}}{\partial x_i^2}(p_0) &= \sum_n \mathbf{q}_n e^{-\frac{1}{2}\sum_{l=1}^{d}\frac{(\Delta x_{n_l})^2}{\sigma^2}} \frac{(\Delta x_{n_i})^2-\sigma^2}{\sigma^4} + \frac{1}{\sigma^2} \\ &= \sum_n \frac{\mathbf{q}_n}{\sigma^2}\left[e^{-\frac{1}{2}\sum_{l=1}^{d}\frac{(\Delta x_{n_l})^2}{\sigma^2}} \frac{(\Delta x_{n_i})^2-\sigma^2}{\sigma^2} + 1\right].\end{aligned} \tag{S43}$$

Since the values of $\mathbf{q}_n$ are always positive, the fraction $\frac{\mathbf{q}_n}{\sigma^2}$ also is positive. On the other hand, note that $\sum_{l=1}^{d}\frac{(\Delta x_{n_l})^2}{\sigma^2} > 0$ and $0 < |\Delta x_{n_i}| \leq \sigma$ which imply that:

$$0 < e^{-\frac{1}{2}\sum_{l=1}^{d}\frac{(\Delta x_{n_l})^2}{\sigma^2}} < 1 \quad \text{and} \quad -1 < \frac{(\Delta x_{n_i})^2 - \sigma^2}{\sigma^2} < 0 \quad \forall n_i. \tag{S44}$$

Therefore the expression

$$e^{-\frac{1}{2}\sum_{l=1}^{d}\frac{(\Delta x_{n_l})^2}{\sigma^2}} \frac{(\Delta x_{n_i})^2 - \sigma^2}{\sigma^2} \tag{S45}$$

belongs to $(0, 1)$ and the term in brackets in equation S43 is always positive:

$$e^{-\frac{1}{2}\sum_{l=1}^{d}\frac{(\Delta x_{n_l})^2}{\sigma^2}} \frac{(\Delta x_{n_i})^2 - \sigma^2}{\sigma^2} + 1 > 0. \tag{S46}$$

The hessian matrix then takes the form:

$$H(MS(p_0)) = \begin{bmatrix} a_1 & \dots & 0 \\ \vdots & \ddots & \vdots \\ 0 & \dots & a_d \end{bmatrix}, \tag{S47}$$

where $a_i = \frac{\partial^2 MS}{\partial x_i^2}(p_0)$. Note that only the elements in the diagonal are positive, on the other hand, the remaining elements are 0. Leading minors of $H(MS(p_0))$ are $m_l = \prod_{s=1}^{l} a_s > 0$ and $H(MS(p_0))$ is positive-definite. The proof is complete. □

#### 5. SUPPLEMENTARY NOTE 5. MEANSHIFT SUPER-RESOLUTION ALGORITHM

The *MS* can be conceptualized as an uphill iterative procedure to characterize data density. *MS* was proposed by Fukunaga and Hostetler [4] with statistical purposes for clustering data as the *MS* vector converges locally to cluster centroids by choosing an appropriate bandwidth. A generalization of *MS* was given by Cheng describing concepts as "blurring process" and "shadow kernel" [5]. The most popular application was provided by Comaniciu and co-workers using *MS* for image segmentation, filtering and object tracking [6, 8, 9]. Moreover, the *MS* has been employed for super-resolution microscopy algorithms as a secondary tool for single molecule localization and cluster segmentation [10]. An interesting application of *MS* was used for denoising and deblurring of images affected by Poisson noise [11] and drift correction [12].

*MSSR* (**M**ean**S**hift **S**uper-**R**esolution) is based on the MS theory and aims to reduce the loss of spatial resolution caused by the diffraction of light. It focuses on density gradients rather than the position of single emitters. In this sense, *MSSR* differs from the classical iterative conception of *MS* theory for finding modes (emitters centers) by successive computation of the *sample mean*. Our approach is based on the calculation of the *MS* value at the first iteration.

The *MSSR* algorithm is supported by spatial analysis which results in an extended-resolved *MSSR* image and temporal analysis that integrates all the information over an extended-resvolved *MSSR* stack. The appropriated combination of both analyses can reveal nanoscopic details from diffraction-limited images. Supplementary Figure S4 shows a flowchart of the *MSSR* algorithm and a description of each step (further explained in the following sections of this supplementary note).

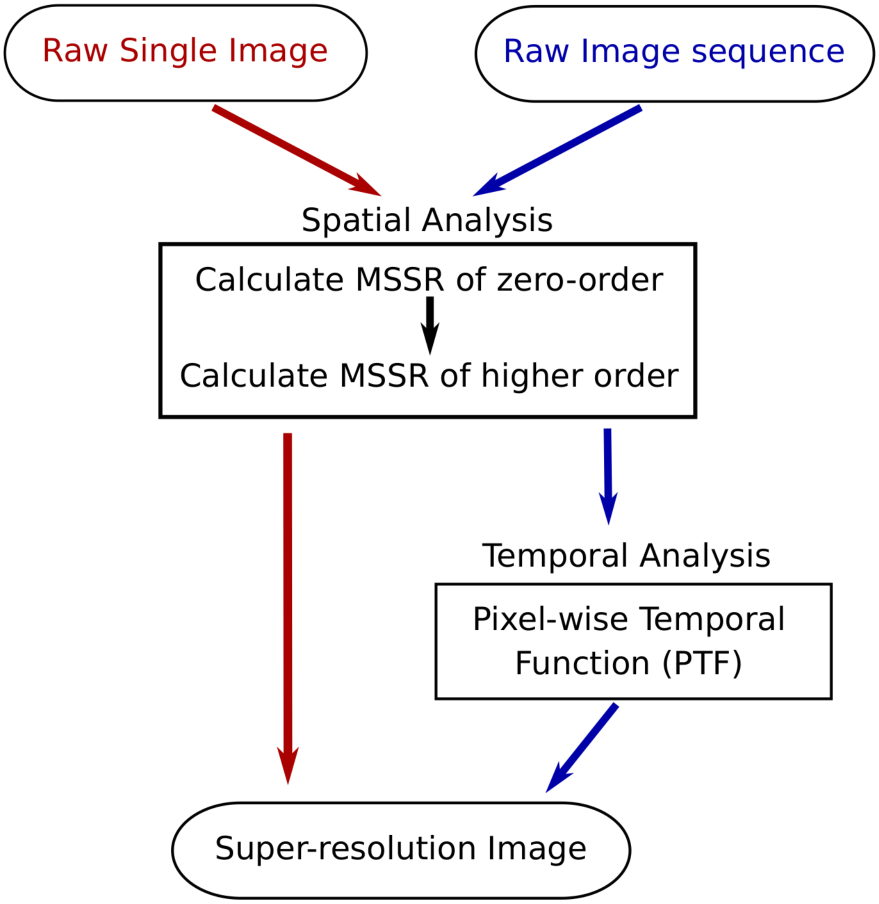

**Figure S4. Flowchart of *MSSR* algorithm.** *MSSR* processes a single image or an image sequence resulting an extended- or super-resolution reconstruction respectively. Spatial analysis is a common step for both type of data, in which, *MSSR* of zero-order or higher-order is calculated. Temporal analysis is an exclusive step for image sequence data type using a given pixel-wise temporal function (*PTF*) for super-resolution reconstruction.

##### 5.1. Spatial analysis of MSSR

The single-frame analysis of $MSSR$ is divided into two main stages: order 0 ($MSSR^0$) and higher orders ($MSSR^n$, $n > 0$). Each stage contributes to the enhancement of the image resolution. A detailed description of each stage is described next.

###### 5.1.1. MSSR of zero order

Given an image $I$, we compute $MS$ with spatial-range $\mathbf{h} = (h_s, h_r)$ bandwidth and profile $\mathbf{g}$ (Supplementary Notes 2 and 3). The kernel slides throughout the hold image computing **SM** and subtracting it from the image value at the center of the kernel (Supplementary Figure S5, Supplementary Note 3). We assume that emitters are affected by a Gaussian or Bessel $PSF$ distribution (Supplementary Note 1).

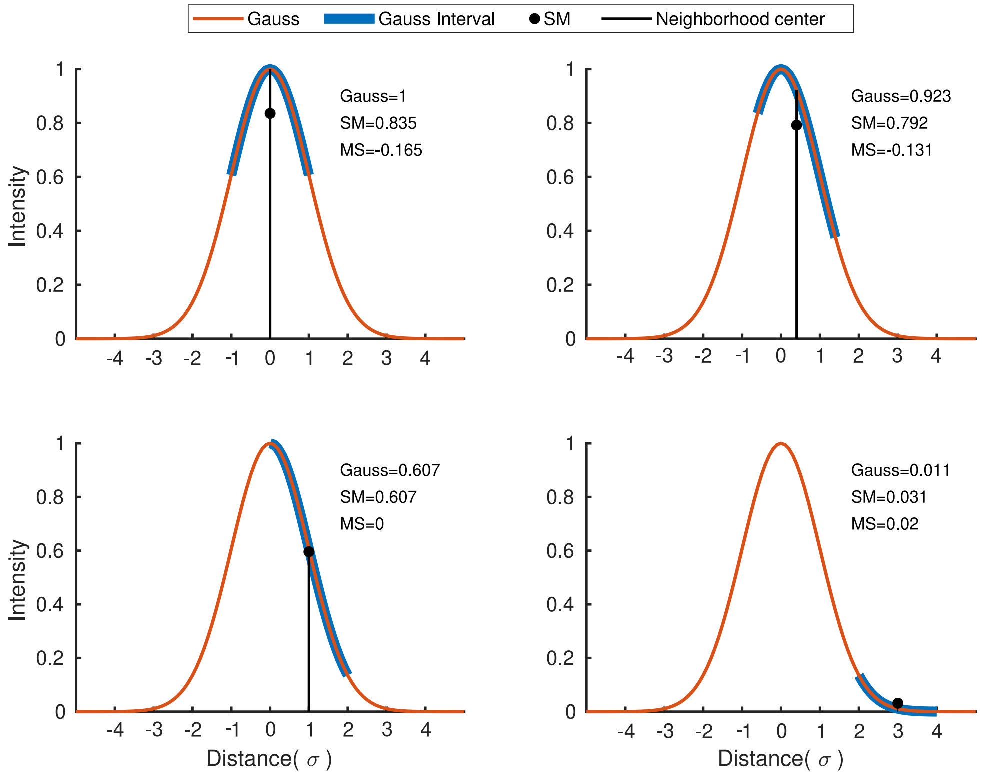

**Figure S5. Different sample mean values according to different neighborhoods of radius $\sigma$ in a Gaussian distribution.** The red, blue and black lines correspond to a Gaussian distribution, the Gaussian interval and the center of the interval, respectively. The black dot is the sample mean (SM) of the interval. In all graphs, the black vertical line intercepts with SM and the Gaussian curve. In the top-left graph, the interval is symmetric and centered at zero, the sample mean associated to this region is below the curve. A similar case is shown in the top-right graph, where the center of the interval has been shifted slightly to the right and the sample mean is closer than that of the previous graph. The bottom-left graph shows the interval centered at $\sigma$ and the sample mean match with the curve. Bottom-right graph shows the interval center at $3\sigma$ (a point located beyond than $\sigma$), note that the sample mean is above the curve.

The $MS$ vector points out towards the point of maximum local density (Supplementary Note 2); its magnitude is greater in local minimum density points than in local maximum density points. For isolated emitters, the point of local maximum density matches the $MS$ and draws a valley for $MS$ (Supplementary Note 4). On the other hand, the points of local minimum density of $MS$ slice the MS curve in three regions: the central region contains the fluorophore's position centered at the local maximum density point (Supplementary Figure S6). Theorem 1 demonstrates that the point of maximum value of a Gaussian function is a point of local minimum of $MS$ (Supplementary Note 4).

The more negative the $MS$ value, the greater the probability of the presence of fluorophores, while positive values of $MS$ indicate their absence. For this reason, the negative value of $MS$ is subtracted to reduce the effect of the $PSF$, hence obtaining an image with higher spatial resolution. The following expression is the negative of $MS$:

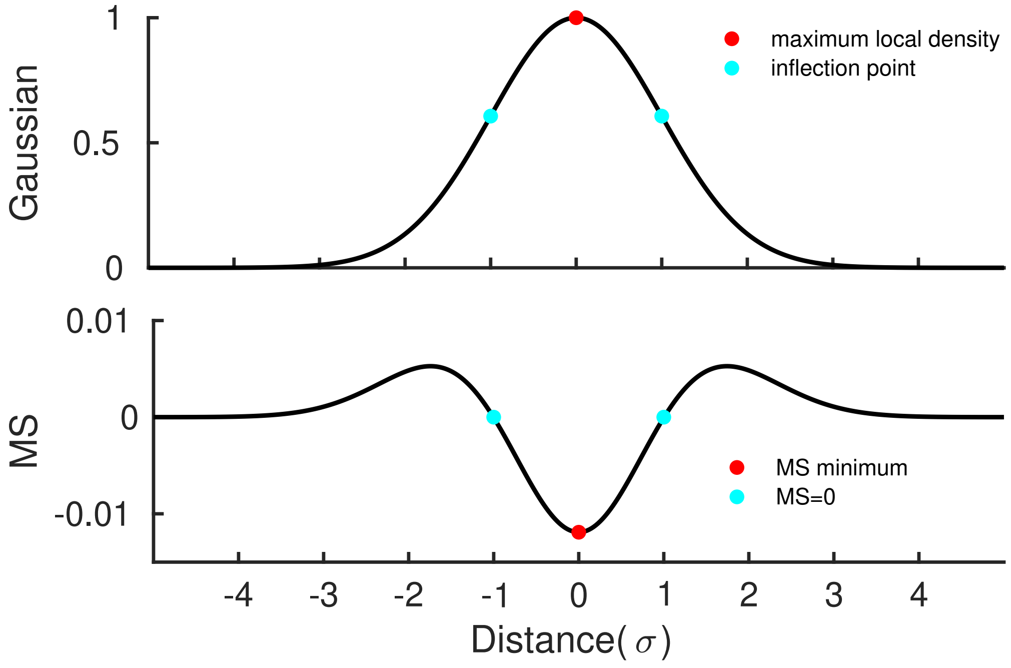

**Figure S6. Main points of Gaussian distribution and its corresponding** $MS$ **result.** On the top graph, the red point is corresponds to both the maximum intensity point and maximum local density point of the Gaussian distribution at the same time. On the bottom graph, the red point is a minimum local point for $MS$. On the other hand, blue points mark where $MS$ is equal zero at $\sigma$, which match with the Gaussian inflection points.

$$MSSR(I) = -MS(I) \tag{S48}$$

Negative values of $MSSR$ are set to zero and the resulting image is normalized to the maximum value of the original image $I$. This step is named by $MSSR$ of order zero and is denoted by $MSSR^0$.

Since $MSSR$ is the negative of $MS$, a direct consequence from theorem 1 is the following corollary:

**Corollary 1** (Inferred from theorem 1)**.** *Let G be a Gaussian distribution and* $p_0$ *the point of maximum value of G, then* $MSSR(p_0)$ *is a local maximum.*

This is an important result for our purpose, since it indicates that fluorophores can be located with high precision by $MSSR$.

A comparison of $MSSR^0$ for different values of $h_r$ and $h_s$ applied to a Gaussian distribution suggests the use of an adaptive range bandwidth (Supplementary Figure S7 a-b). Note that for fixed $h_r$ and varying $h_s$, there is almost no difference in the $MSSR^0$ outcome (Supplementary Figure S7a). On the other hand, by fixing $h_s$ and varying $h_r$, artifacts start to appear at the tails of the distributions (Supplementary Figure S7b). For this reason, the range bandwidth $h_r$ is adaptive to each range neighborhood taking the maximum difference among its center and neighbors (Supplementary Figure S7c). The use of a uniform kernel will cause the effect in S7a (see inset box), where distributions for different $h_s$ would be practically indistinguishable. While a Gaussian kernel is more suitable to achieve the distinction among $MSSR^0$ results as shown in S7c.

In Fourier space, the objective lens of a microscope act as a finite aperture, the the optical transfer function of a traditional microscope can conveniently be represented by a circle, the diameter of which can be linked directly to the maximum observable spatial frequency $K_0$, where

$$K_0 \approx \frac{1}{FWHM}, \tag{S49}$$

reviewed in [13]. The $PSF$ can be simulated as an Airy pattern or Bessel beam according to the features of the optical system: wavelength, numerical aperture, and the refractive index of the medium. It can also be measured experimentally from imagining diffraction limited fiducial markers. The selection of $h_s$ was based on the fact that, the $PSF$ contains valuable information of

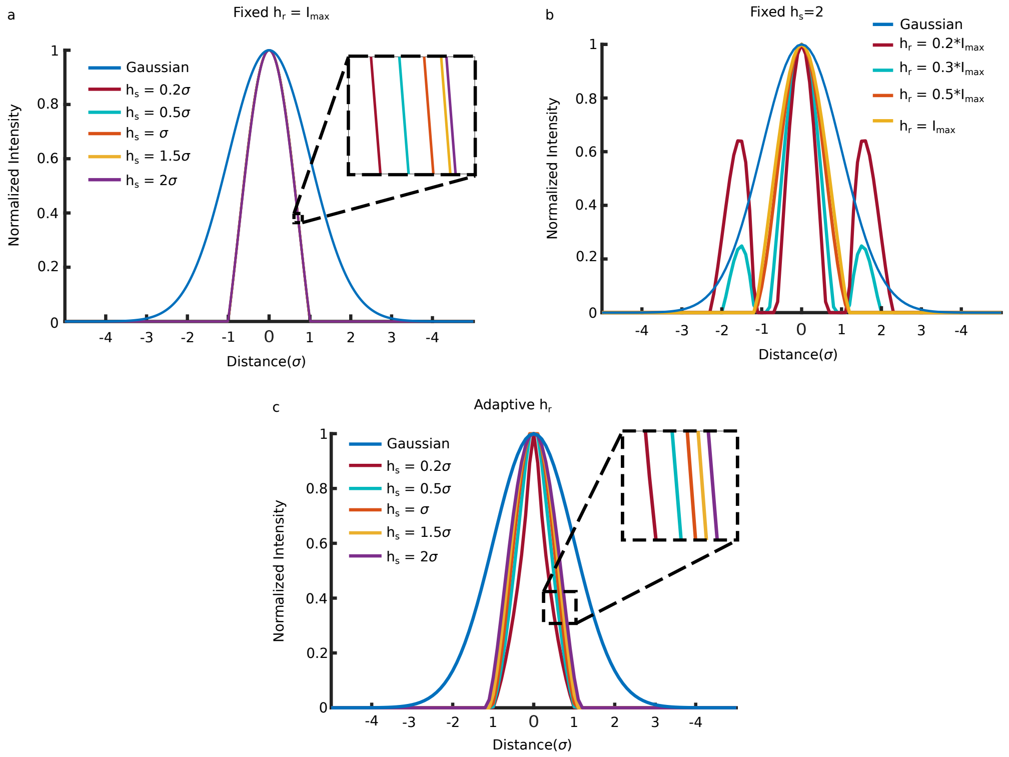

**Figure S7.** $MSSR^0$ **on variations of** $h_r$**,** $h_s$**, adaptive** $h_r$ **and the selection of** $h_s$**. a)** Comparison of a Gaussian distribution and $MSSR^0$ by using a fixed $h_r$ equal to the maximum intensity of the distribution and varying $h_s$. $MSSR$ values are almost indistinguishable. **b)** Comparison of a Gaussian distribution and $MSSR^0$ by using a fixed $h_s$ equal to the maximum intensity of the distribution and varying $h_s$. Lower values of $h_r$ generate artifacts on the tail of the $MSSR^0$ distribution. **c)** Comparison of a Gaussian distribution and $MSSR^0$ by using an adaptive $h_r$.

the used optical system (Supplementary Note 1). $h_s$ is defined as the half of the full width at half maximum ($FWHM$) of the $PSF$. Supplementary Figure S8 shows the relationship between the $h_s$ and $FWHM$ in a Airy pattern.

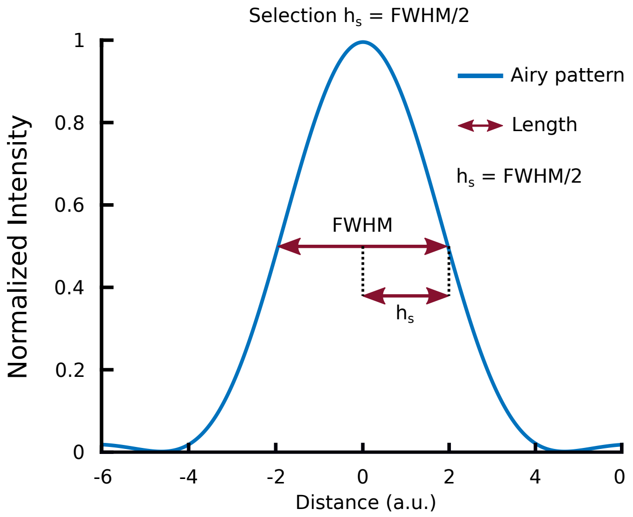

**Figure S8. Selection of spatial parameter** $h_s$ **related to** $FWHM$**.** Given a $PSF$ given by an Airy pattern, the $h_s$ is chosen as the half of the $FWHM$.

Comparative results of $MSSR^0$ applied over a Gaussian or a Bessel distribution are presented in Supplementary Figure S9 (the relationship between the $PSF$ and each type of distribution is discussed in Supplementary Note 1). $MSSR^0$ reduces the radius of a Gaussian distribution by about a half. The Bessel distribution is affected in a similar way, $MSSR^0$ reduces the radius of the main lobe of the distribution. Moreover, the concentric rings of the Bessel distribution are reduced in width and intensity and as consequence they can not be distinguished anymore. For this reason, near emitters will not be affected by Airy discs away of the central lobe.

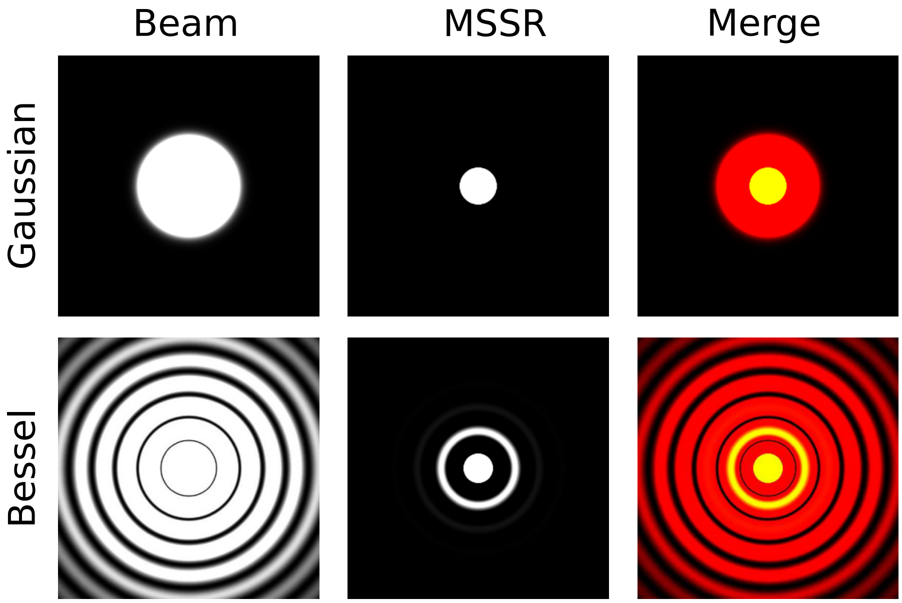

**Figure S9.** ***$MSSR^0$* applied to Gaussian and Bessel distributions.** First row: Gaussian beam and its corresponding $MSSR^0$ result. Second row: Bessel beam and its corresponding $MSSR^0$ result. By column from left to right: beam type, $MSSR^0$ result and merge of the previous images. All images have been threshold by the same level at 0.4% of the maximum intensity to show the concentric rings.

Supplementary Figure S10a shows an emitter simulated an Airy pattern using a wavelength of 488 nm and the corresponding result of $MSSR^0$ is shown in Supplementary Figure S10b. In both graphs, a Gaussian distribution is fitted to obtain the respective standard distribution. Supplementary Figure S10c shows the linear dependence of $\sigma$ and each modelled distribution at different wavelengths of the visible spectrum.

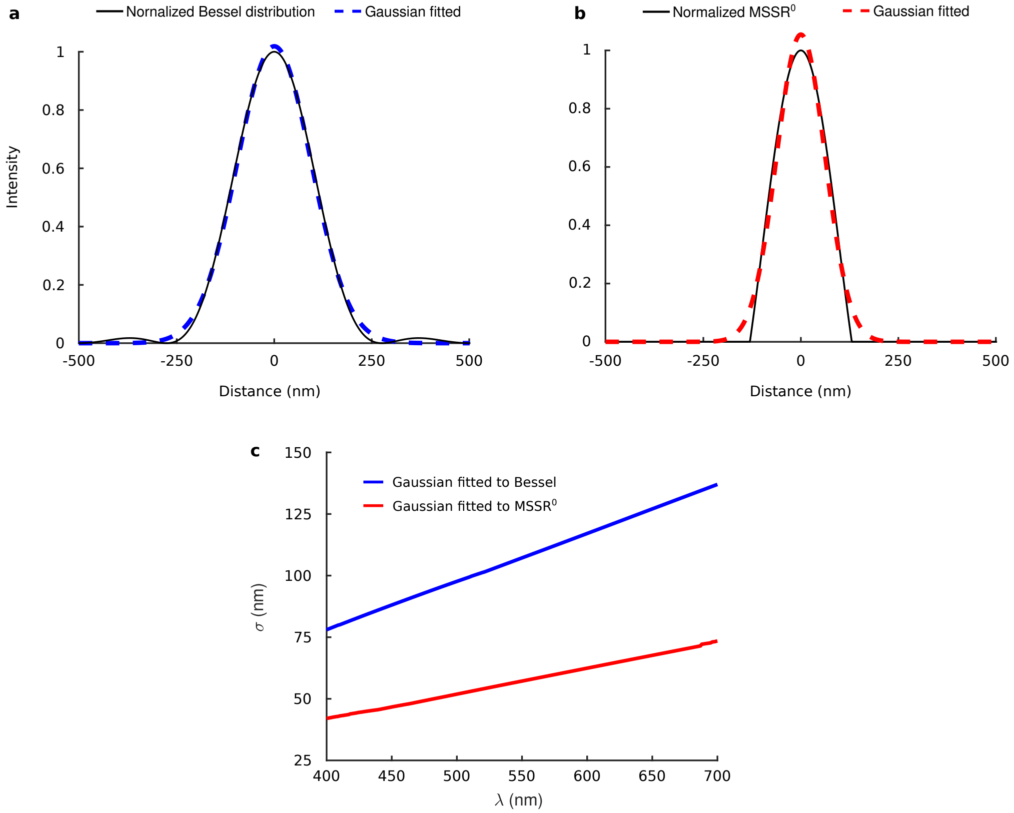

**Figure S10. Standard deviation of the Gaussian distribution fitted to Bessel distribution and the corresponding result of $MSSR^0$ at different wavelengths. a)** Gaussian fitted (discontinuous blue line) to Bessel distribution (continuous black line) of the emitter simulated with a wavelength of 488 nm. **b)** Gaussian fitted (discontinuous red line) to $MSSR^0$ (continuous black line) of the Bessel distribution obtained in a). **c)** Standard deviation of the Gaussian ($\sigma$) fitted to Bessel distribution (blue line) and the corresponding result of $MSSR^0$ (red line).

###### 5.1.2. MSSR of higher orders

Higher-order $MSSR$ ($MSSR^n$, $n > 0$) encompasses a sequence of iterative operations which increase the resolution of the image.

Since $FWHM$ of the emitter have been reduced from the original image $I$ to $MSSR^0$, the difference between $I$ and $MSSR^0$ is calculated, which results in a saddle point at the center of the emitter referred as Diff. Next, the complement of Diff is calculated and normalized. Note that intensities out of the interval $[-\sigma, \sigma]$ are different from zero, and causes possible artifacts. For this reason, an intensity weighting with the $MSSR^0$ image is required to remove undesirable artifacts. This gives $MSSR$ its iterative nature, increasing the resolution after each step, called *order* and denoted by $n$ (Supplementary Figure S11).

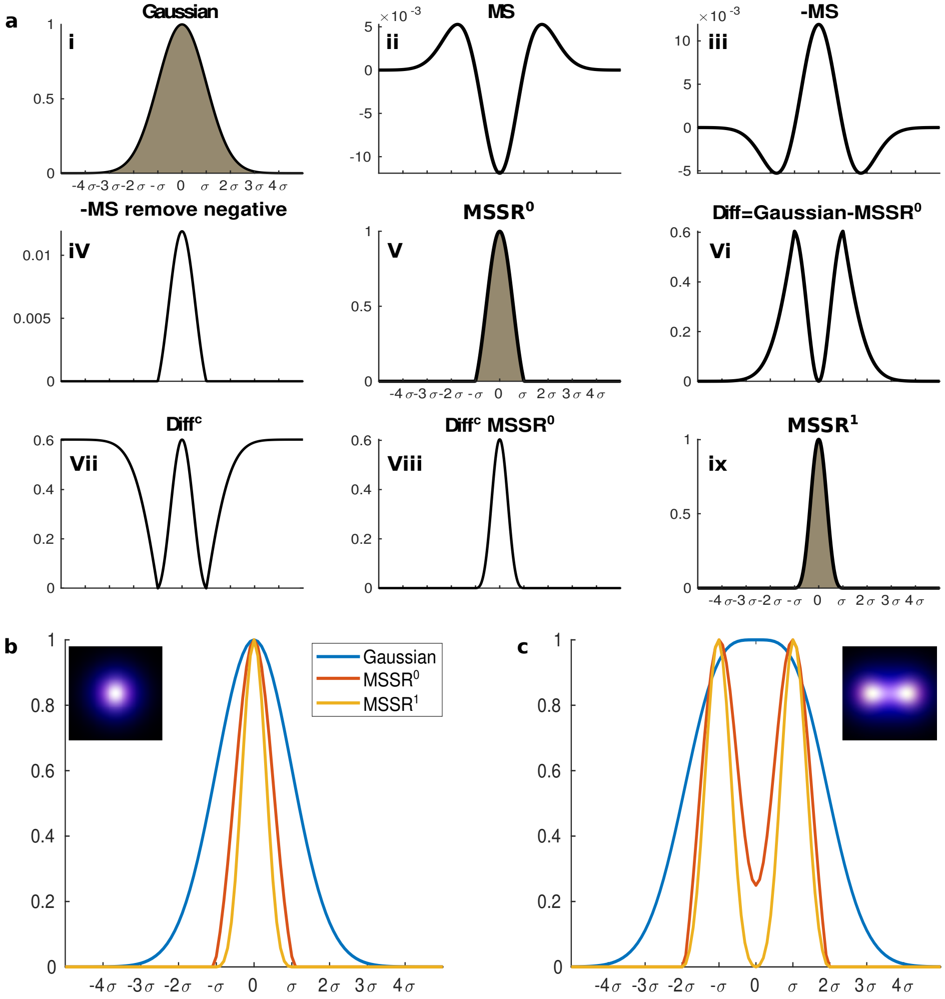

**Figure S11.** ***MSSR*** **reduces the Full Width Half Maximum (*FWHM*) of a Gaussian distribution as the *MSSR* order increases. a)** Steps of $MSSR$ procedure: analytical steps required for $MSSR$ of zero order (i-v) and for higher orders (vi-ix). The MS is applied to the initial fluorescence distribution (i) resulting in a MS graph (ii). The negative of MS is calculated to set MS minimum and maximum (iii). Negative values are set to zero (iv); positive values correspond to the main lobe of the $PSF$ (Supplementary Figure S7). $MSSR$ of zero order ($MSSR^0$) reduces the $FWHM$ by $1.4\sigma$ (v). The difference between the input image and the $MSSR^0$ creates a valley located between two maximum points at $\pm\sigma$ (vi). The complement is computed (vii) and intensity weighting of the result of $MSSR^0$ is performed (viii). Finally, a normalization step provides the $MSSR$ of order 1 ($MSSR^1$) (ix). **b)** Line profiles of a single emitter simulated as a Gaussian distribution before (blue) and after $MSSR^0$ (red) or $MSSR^1$ (yellow). **c)** Analogous case as in (b) but using two emitters located at $\pm\sigma$. Note that Gaussian distribution (blue stroke) is flat at the top center, indicating that emitters are unresolved. However, applying $MSSR^0$ forms a saddle point (valley) between emitters, hence they can be distinguished from one another. By applying $MSSR^1$ the height at the saddle point turns to zero and emitters become completely resolved.

The dependence of the $FWHM$ of a Gaussian distribution decreases as the number of iterations of the $MSSR$ increases (Supplementary Figure S12 a-c). $MSSR$ order is commonly set to a maximum of 2, given that part of the emitter's information is degraded with each iteration(Supplementary Figure S12).

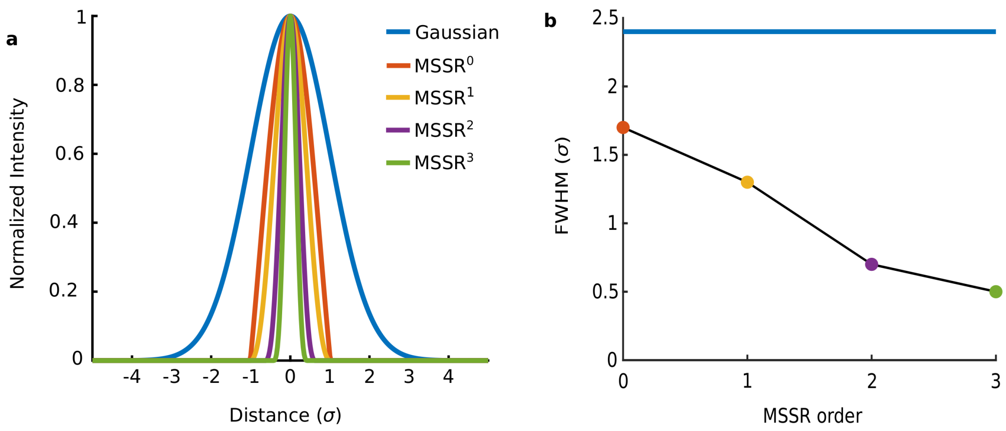

**Figure S12. FWHM decreases with increasing $MSSR$ order over a Gaussian distribution. a)** Comparison of Gaussian distribution and $MSSR$ of order 0 to 4. **b)** $FWHM$ for distributions presented in a).

Supplementary Figure S13 shows the procedure of thresholding and normalizing the joint distribution shaped by two adjacent emitters located at the Rayleigh limit, note that the dip decreases from a value of 0.8 to 0.57 (Supplementary Figure S13 a and b). Hence, it is tempting to conclude that there is a noticeable increase in resolution as the Rayleigh limit has been surpassed. By repeating the same procedure using a joint distribution shaped by two adjacent emitters located at the Sparrow limit there is no further gain of resolution as the dip remains constant, taking the value of 1 (Supplementary Figure S13 e and f). The Sparrow limit is associated to the maximum observable spatial frequency, hence, to the diffraction barrier. Noteworthy, processing the previous joint distributions with $sf - MSSR$ of any order leads to a decrease of the dip value (Supplementary Figure S13 e and f). The latter observation becomes striking when studying the case of $sf - MSSR^3$ processing which collapses the dip to zero for both Rayleigh and Sparrow conditions.

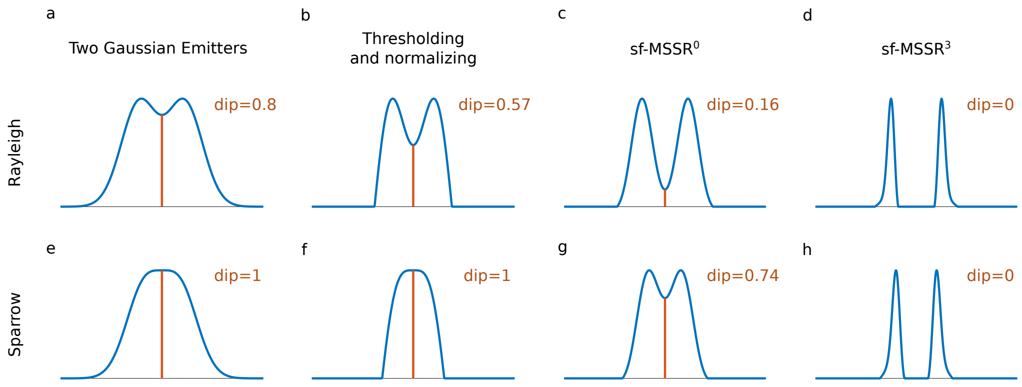

**Figure S13. Thresholding and normalizing processes are not enough to overcome the Sparrow limit.** This figure shows the sequential decrease in dip height between two intensity Gaussian distributions from simulated point emitters (a,e). Thresholding and normalizing (b,f) provides an initial reduction in dip height, which continues to decrease as MSSR of higher order is applied (c,d,g,h). Upper and bottom rows depict the relative distance between the peaks of both distributions when either the Rayleigh (top) or Sparrow (bottom) criteria are met. Worth noticing is that higher-order MSSR provides a complete dip collapse to zero, separating the distributions of both emitters completely.

$MSSR^0$ reliably provides spatial resolution gains, free of image analysis artifacts, down to the Rayleigh limit, hence, allowing the study of nanoscopic regimes at the boundaries of the

Sparrow limit (Supplementary Figure S14). $MSSR$ of higher orders ($MSSR^n$: $n > 0$) encompass further analytical operations (subtraction, complement, negative and intensity weighting) which allow the observer to distinguish between single emitters separated by ∼0.66 times the Gaussian $FWHM$ of the $PSF$ (1.55 times the PSF s.d. $\sigma$).

$MSSR$ aims to revert the effect of diffraction on optical microscopy so it must be considered as a deconvolution process. What makes $sf - MSSR^n$ unique is the fact that it restores spatial information down below the Sparrow limit. Supplementary Figure S14 shows that Wiener deconvolution partially deconvolves the effect of diffraction, but without a dramatic increase in spatial resolution. Interestingly, Richardson-Lucy deconvolution provides a noticeable increase in resolution at the boundaries of the Rayleigh limit but fails to restore spatial resolution down the Sparrow limit (Supplementary Figure S14). With such data in hand, we conclude that $sf - MSSR^n$ is the first deconvolution approach of optical microscopy that restores spatial information at the nano scales.

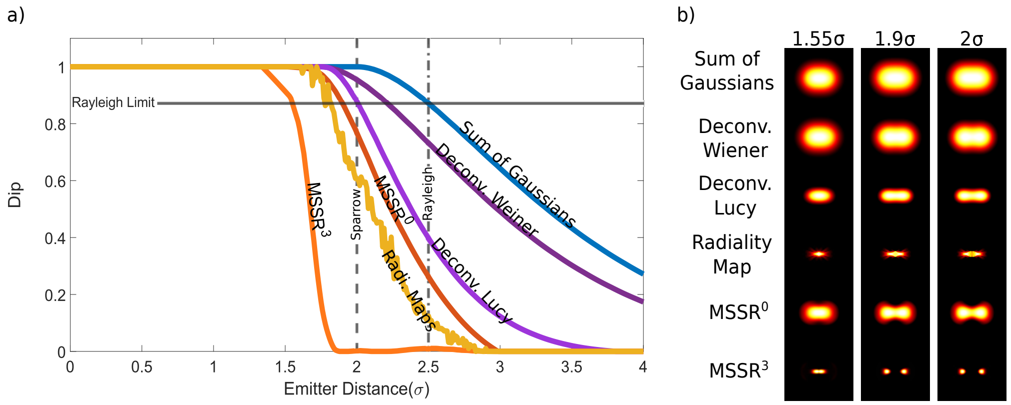

**Figure S14. Resolution increase provided by deconvolution methods, radiality maps and $sf - MSSR^n$. a)** All methods tested decrease the dip successfully. The horizontal line represents the Rayleigh limit in relation to the dip height (for the sum of Gaussians, 2.5$\sigma$). This value decreases the most when using radiality maps and MSSR, where the best result reaches Rayleigh limit = 1.55$\sigma$ (1.61 times smaller), provided by $MSSR$ of third order (orange line). **b)** Bidimensional representations of the reconstructed images provided by each method tested, for emitters separated at distances of 1.55$\sigma$, 1.9$\sigma$ and 2$\sigma$. Simulated emitters were generated with the gaussian distribution with a sigma of 121.10 nm ( 10 pixels) using the following parameters: Refractive index of medium = 1.33, Refractive index of oil = 1.515, NA = 1.4, wavelength = 610 nm, pixel size = 11.7 nm.

##### 5.2. Temporal analysis of MSSR

The computation of $MSSR$ over a temporal stack of diffraction-limited images $\mathbf{I}$, with $M$ frames collected at a static or pseudo-static scene, generates a temporal stack of $M$ super-resolved replicates.

A temporal analysis can be used to take advantage of the information stored at the temporal dynamics of the fluorophores, which allows for further resolution enhancement. This step consists in applying a pixel-wise temporal function ($PTF$) across the whole $MSSR$ stack in order to obtain a super-resolved image with higher spatial resolution and diminished noise effects (tab. S2).

Each $PTF$ has an advantage over the other depending on the nature of the fluorescence dynamics of the image stack to be processed. The selection of $PTF$ depends of the characteristic of blinking emitters along the stack (Supplementary Table S2). For example, $Var$ is useful to reveal nano-scaled details if there exists high pixel-wise temporal entropy (e.g. highly fluctuating signal due to fluorophores transiting into the triplet state). $Mean$ is better suited for experimental scenarios with low nanoscopic temporal dynamics, or for scenarios with low signal to noise ratio. $TPM$ can be used over an intermediate level of signal entropy. $CV$ takes into account both the blinking and the intensity of emitters (the latter not being considered with $Var$). Finally, $SOFI$ temporal functions [14] allow to shrink and refine structures with the risk of loss of continuity on low fluorophore density samples.

| PTF Notation | Name | formula | recommended usage |
| --- | --- | --- | --- |
| $Mean$ | Average | $\frac{1}{M}\sum_{i=1}^{M} x_i = \overline{x}$ | low blinking of fluorophores |
| $TPM$ | Temporal Product Mean | $\frac{1}{M}\sum_{i=1}^{M}(x_i x_j)^2$ | intermediate blinking of fluorophores |
| $Var$ | Variance | $\frac{1}{M}\sum_{i=1}^{M}(x_i - \overline{x})^2$ | high blinking of fluorophores |
| $CV$ | Coeffitient of Variation | $Var/Mean$ | high blinking of fluorophores |
| $SOFI_2$ | Auto-cumulant order 2 [14] | $\langle \delta x_i \cdot \delta x_{i+1} \rangle$ | high blinking of fluorophores |
| $SOFI_3$ | Auto-cumulant order 3 [14] | $\langle \delta x_i \cdot \delta x_{i+1} \cdot \delta x_{i+2} \rangle$ | high blinking of fluorophores |
| $SOFI_4$ | Auto-cumulant order 4 [14] | $\langle \delta x_i \cdot \delta x_{i+1} \cdot \delta x_{i+2} \cdot \delta x_{i+3} \rangle$ $- \langle \delta x_i \cdot \delta x_{i+1} \rangle \langle \delta x_{i+2} \cdot \delta x_{i+3} \rangle$ $- \langle \delta x_i \cdot \delta x_{i+2} \rangle \langle \delta x_{i+1} \cdot \delta x_{i+3} \rangle$ $- \langle \delta x_i \cdot \delta x_{i+3} \rangle \langle \delta x_{i+1} \cdot \delta x_{i+2} \rangle$ | high blinking of fluorophores |

$M$: number of frames of $MSSR(\mathbf{I})$,
$x_k$: $MSSR$ image at frame $k$,
$\langle \ldots \rangle$: average over time,
$\delta x_k = x_k - \langle x \rangle$: time lag fluctuation over $k$ frames.

**Table S2.** $MSSR$ **temporal functions and their recommended usage.**

### 6. SUPPLEMENTARY NOTE 6. INTERPOLATION ALGORITHMS

Part of the super-resolution image reconstruction process is an interpolation step that fills the gaps in the amplified image. Implemented in the current version of *MSSR* are two different interpolation algorithms: bicubic and Fourier interpolation. The theoretical foundations of both of these two approaches and their differences are detailed in this Supplementary Note. In addition, a comparison of the performance between these algorithms is provided

#### 6.1. Bicubic interpolation

The bicubic interpolation is an extension of the cubic interpolation, applied to a two-dimensional grid, where polynomial or cubic convolution algorithms are used. For each new pixel of the reconstruction image, the closest 16 surrounding pixels (in a 4x4 array) in the original image are considered (Supplementary Figure S15), a weighted average of these 16 pixels is obtained; the weights are related to each pixel's distance to the new one, given by:

$$v(x^{'}, y^{'}) = \sum_{i=0}^{3}\sum_{j=0}^{3} a_{i,j} A(x_i, y_i), \tag{S50}$$

where $a_{i,j} = \text{cubic}(\delta_x) * \text{cubic}(\delta_y)$, $v(x^{'}, y^{'})$ is the pixel value as a result of the interpolation, $A(x_i, y_i)$ is the original image pixel and $\delta_x$ and $\delta_y$ are the distances between $x^{'}, x$ and $y^{'}, y$, respectively. The cubic interpolation function has the form:

$$\text{cubic}(x) = \begin{cases} (-c+2)x^3 + (c-3)x^2 + 1 & \text{if } 0 \leq x < 1 \\ -cx^3 + 5cx^2 - 8cx + 4c & \text{if } 1 \leq x < 2 \\ 0 & \textit{otherwise} \end{cases} . \tag{S51}$$

where $c$ is a smoothing parameter (set to $c = \dfrac{1}{2}$ as an optimal working value) [15, 16].

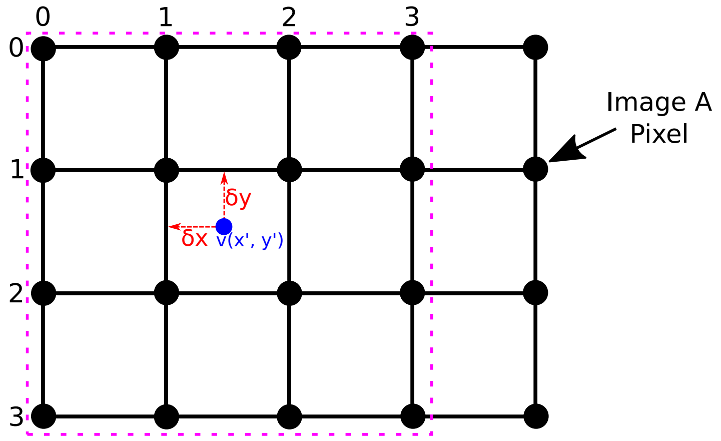

**Figure S15. Scheme of bicubic interpolation.** The black points in the grid represent the 16 near-most pixels in the original image to the new one in the reconstruction, located at $(x^{'}, y^{'})$ (blue point). $\delta_x$ and $\delta_y$ are the distances between the original pixels and the new one, on the $X$ and $Y$ axis respectively. Interpolation is performed using the formula S50 shown above.

##### 6.1.1. Mesh-like pattern originated from bicubic interpolation

Occasionally, reconstructions provided by bicubic interpolation can present a mesh-like pattern. When the interpolation process takes place, a gradient of intensities is generated between the original pixels and the ones that are generated during the process; this is because the interpolation function averages values which are close to the point that is being generated. For each new pixel, the closer it is to the original one, the more it will resemble it. This means that, if a new pixel is in an intermediate point between two pixels, its value will be affected by the information of both sides and will most likely be an average of them both. When this process is repeated over a stack with very similar pixel intensities, the new pixels (in intermediate positions) in each image will tend to retain likeness to each other (upper panel in Supplementary Figure S16c), and thus causing the variance to be rather low. In the upper panel in Supplementary Figure S16c, pixel values that match the dotted-blue lines have a greater dispersion; they correspond to the generated pixels which were the closest to the original ones during the interpolation process.

Supplementary Figure S16 shows an example of the result obtained from an image sequence processed with interpolation and how it is affected. A sequence of white images contaminated with EM-CCD noise (using a custom code written in R based on the theory and equations provided by Hirsch and co-workers in [17]) and amplification equal to 5 is shown in Supplementary Figure S16a; like structures can be easily perceived as the vertical and horizontal lines distribute evenly across the entire frame (lower panel in Supplementary Figure S16a). After careful analysis of the line profiles in Supplementary Figure S16c, it can be concluded that:

- The distance (pixels) between the mesh maxima is equal to the amplification value.
- These lines (mesh) correspond to the local minima, and the midpoint between them to the maxima.
- The distance between a local maximum and minimum is half the amplification value.

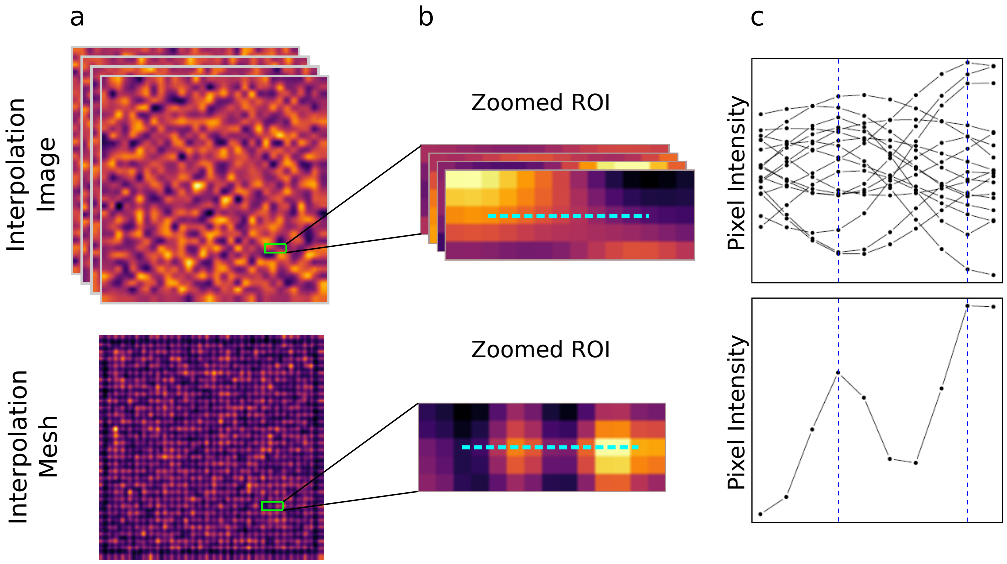

**Figure S16.** **a)** Interpolated white images stack of interpolation and its variance projection (where the mesh is clear). **b)** zoomed region of the images in a). **c)** Line profiles corresponding to the pixel values across the cyan dotted lines in the zoomed *ROI*s. The first fifteen frames are depicted in the upper graph in c). The blue dotted lines in c) represent the position of the maximum values of the mesh across the line profile.

###### *6.1.2. Mesh minimization algorithm*

To compensate for this 'meshing' effect, an information shift in two directions for each axis is performed, resulting in four total shifts, and thus generating four extra images (Supplementary Figure S17). The magnitude of the shift in either direction is equal to half the amplification value, as this corresponds to the distance from a given local maximum and its minimum. These four extra images along with the original are averaged and the result is the new compensated image.

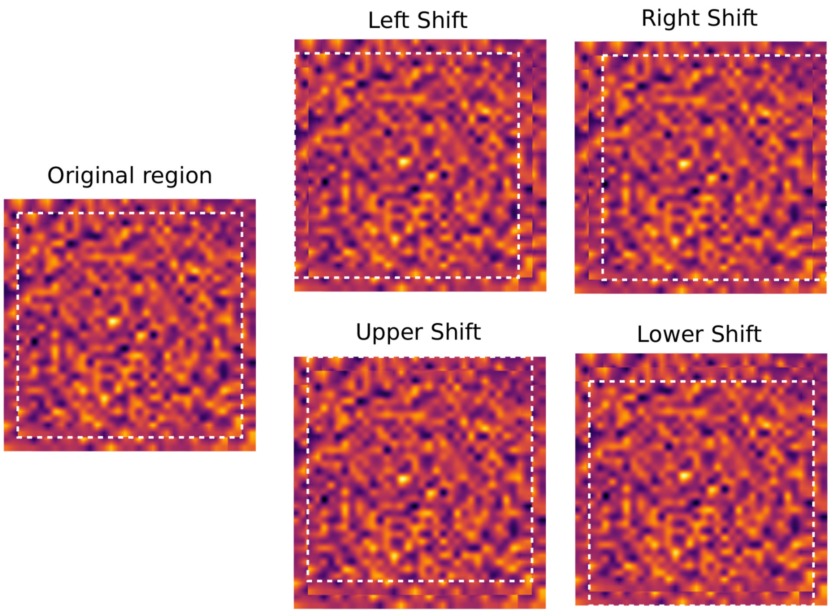

**Figure S17. Scheme of the 'meshing' compensation.** A total of four extra images are generated as a result of spatially shifting the pixel information across both sides of each axis.

###### *6.1.3. Comparison between uncompensated and compensated data*

In order to compare the performance of the bicubic interpolation with and without the mesh-minimization option and assess its effectiveness, a set of synthetic (Supplementary Figure S18-S20) and experimental (Supplementary Figure S21 and S22) data was used.

A stack of white images contaminated with EM-CCD noise and the synthetic data SynEx2 (https://sites.google.com/site/uthkrishth/musical) with and without EM-CCD noise were used to demonstrate the 'meshing' minimization effect during the (bicubic) interpolation process. A comparison of the results before and after the compensation is presented in Supplementary Figures. S18-S20.

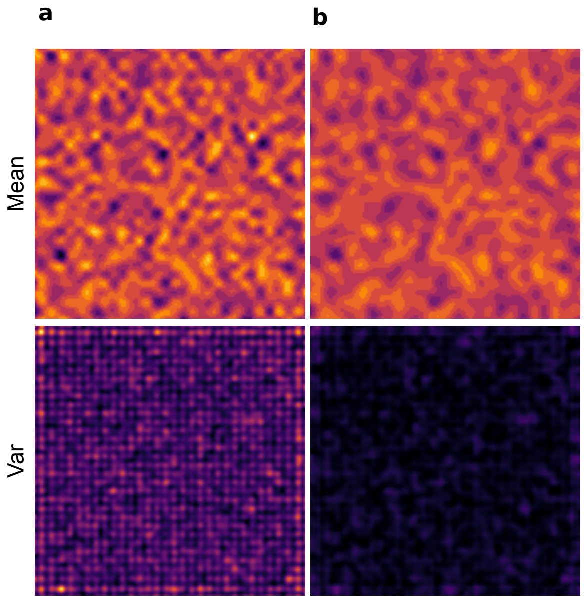

**Figure S18. Example of the result of bicubic interpolation and mesh compensation in a noisy image. a)** without the mesh compensation. **b)** with mesh compensation.

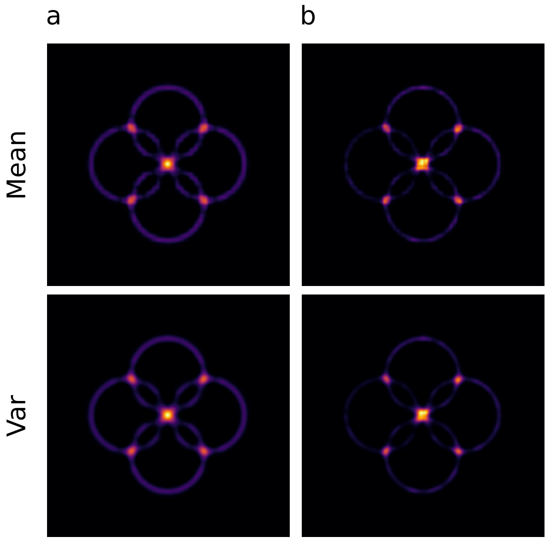

**Figure S19. Example of the result of bicubic interpolation and mesh compensation in an image without noise. a)** without the mesh compensation. **b)** with mesh compensation.

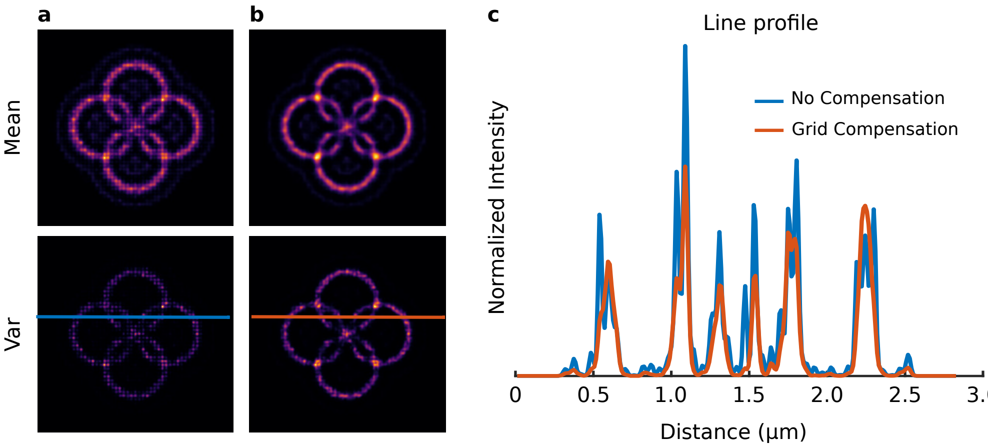

**Figure S20. Example of the result of bicubic interpolation and mesh compensation in a noisy image. a)** without the mesh compensation. **b)** with mesh compensation. **c)** Line profiles of the variance results from dashed lines in a) and b). The blue line in the images correspond to the values from a) and the red line to b).

The lack of noise in the synthetic data notably reduces the generation of visible artifacts (Supplementary Figure S19), whereas the presence of noise greatly increases their emergence (Supplementary Figure S18 and S20). The 'meshing' effect will always take place when real experimental data is analyzed, due to noise being intrinsically and inevitably introduced along the imaging process in any optical system.

We further assessed the effectiveness of the mesh-minimization algorithm using real experimental data. A comparison of the results before and after the compensation is presented in Supplementary Figures. S21 and S22. First, the *MSSR* analysis was performed on a single image (first image from stack, Supplementary Figure S21). Then, the *MSSR* analysis was performed on the entire stack along with three different temporal analyses (Supplementary Figure S21).

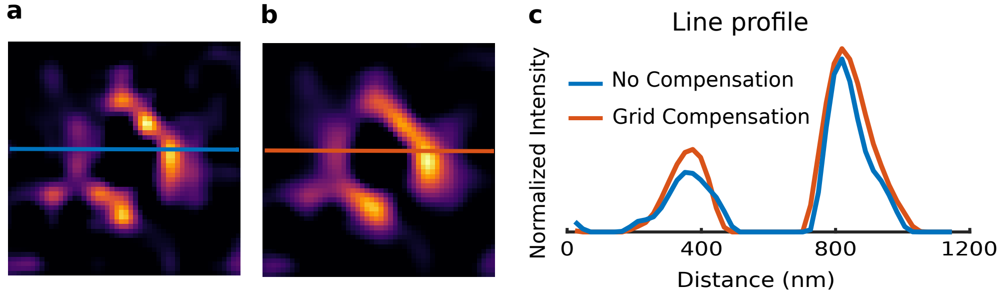

**Figure S21. Example of the mesh compensation over a** $MSSR$ **image obtained from a single noisy frame. a)** without the mesh compensation. **b)** with mesh compensation. **c)** Line profiles of the $MSSR$ results from a) and b), the blue line in the images correspond to the values from a), and the red line to b).

As can be seen, the use of $MSSR$ without compensation in a single image does not generate visible artifacts, but when the process is carried out on several images (stack) and a temporal analysis is applied, the mesh can emerge. Supplementary Figure S22 shows that the mesh pattern is only present in the Variance temporal analysis. However, even if the mesh pattern is not present, the reconstruction images may contain other artifacts, such as the four-sided polygon-like shape inside the toroids in Supplementary Figures. S21a and S22a. When the compensation is applied, the interior of the toroids adopts a more circular (expected) shape (Supplementary Figures. S21b and S22b).

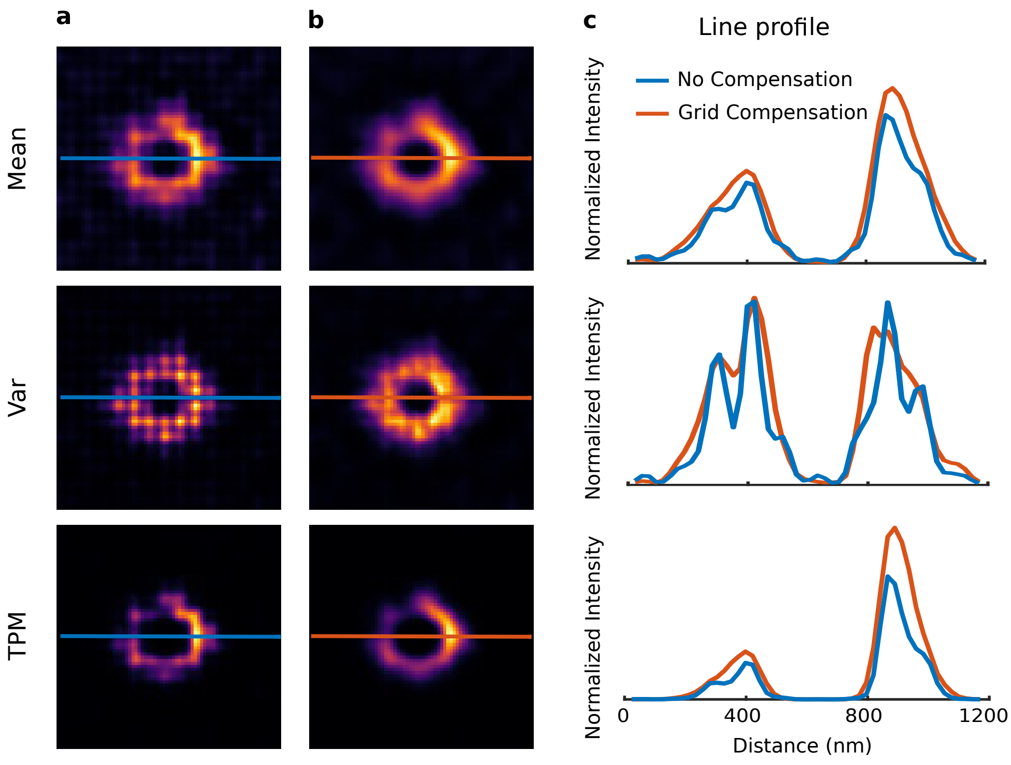

**Figure S22. Example of the result of** $MSSR$ **image and mesh compensation in a noisy image stack. a)** without the mesh compensation. **b)** with mesh compensation. **c)** Line profiles of the $MSSR$ results from a) and b), the blue line in the images correspond to the values from a), and the red one to b).

##### 6.2. Fourier interpolation

Fourier interpolation on images is an extension of the Fourier interpolation for 1-dimensional data and uses the Fast Fourier Transform.

The Fourier interpolation on images its made up by 5 basic steps.

1. apply Fast Fourier Transform ($FFT$) to translate an image to the frequency domain

2. apply zero-padding by adding rows and columns with value equal to zero.

3. frequency compensation by multiplying by the image factor magnification.

4. apply inverse Fast Fourier Transform ($iFFT$) to go back the image in the spatial domain.

5. compute the absolute values of the the previous step.

When translating an image from the spatial domain into the frequency domain, since both spaces share the same information and only its representation changes, no information is lost during the process. Zero-padding serves two purposes: a) preserve the original information of the image and b) ensure the reconstructed image has the desired amplified dimensions. The third step involves the compensation of the frequencies by multiplying by the image amplification factor (AMP parameter in the MSSR plugin). The reasoning behind this step is that, when the number of frequencies increases, this causes the information contained within the spatial domain to spread and stretch along a new period altered by the amplification factor, which is used to compensate the information in the amplified image. Finally, the application of the inverse $iFFT$ and the computation of its absolute value take place in order to obtain the final amplified, Fourier-interpolated image in the spatial domain. A diagram of the previous description is depicted on figure S23.

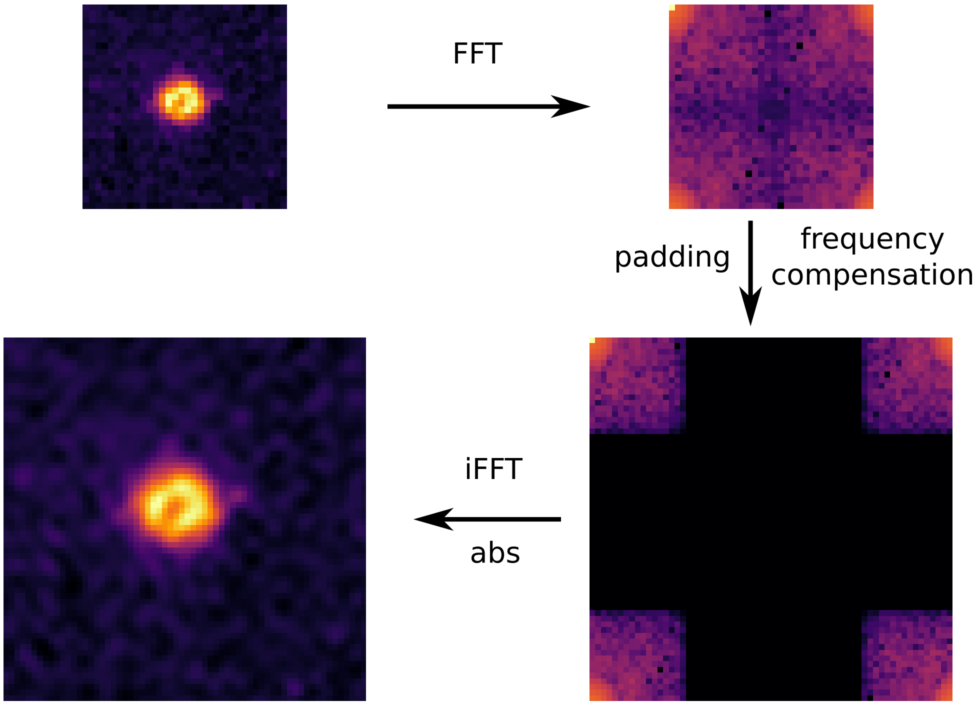

**Figure S23. Fourier interpolation in 2D.** Fourier interpolation is based on the application of direct and inverse Fast Fourier Transform, including zero-padding, frequency compensation and the computation of the absolute values of the image.

###### *6.2.1. Comparison of interpolation algorithms performance*

Both of the currently implemented interpolation algorithms are entitled to some advantages and disadvantages. Bicubic interpolation is less computationally demanding and generally provides shorter run times (a comparison of run times for both types of interpolation is provided in Supplementary Figure S24). Though, it is more prone to mesh-like artifact generation under certain experimental conditions. In most cases, Fourier interpolation does not require a mesh minimization procedure, although it might help mitigate background artifacts. Since it tends to provide better results when performing a temporal analysis, it is suggested to opt for Fourier interpolation over bicubic in such regard. Given that the mesh-like artifact arises only when a temporal analysis is performed (see Supplementary Note 6.1) bicubic interpolation is advised for single-frame analysis.

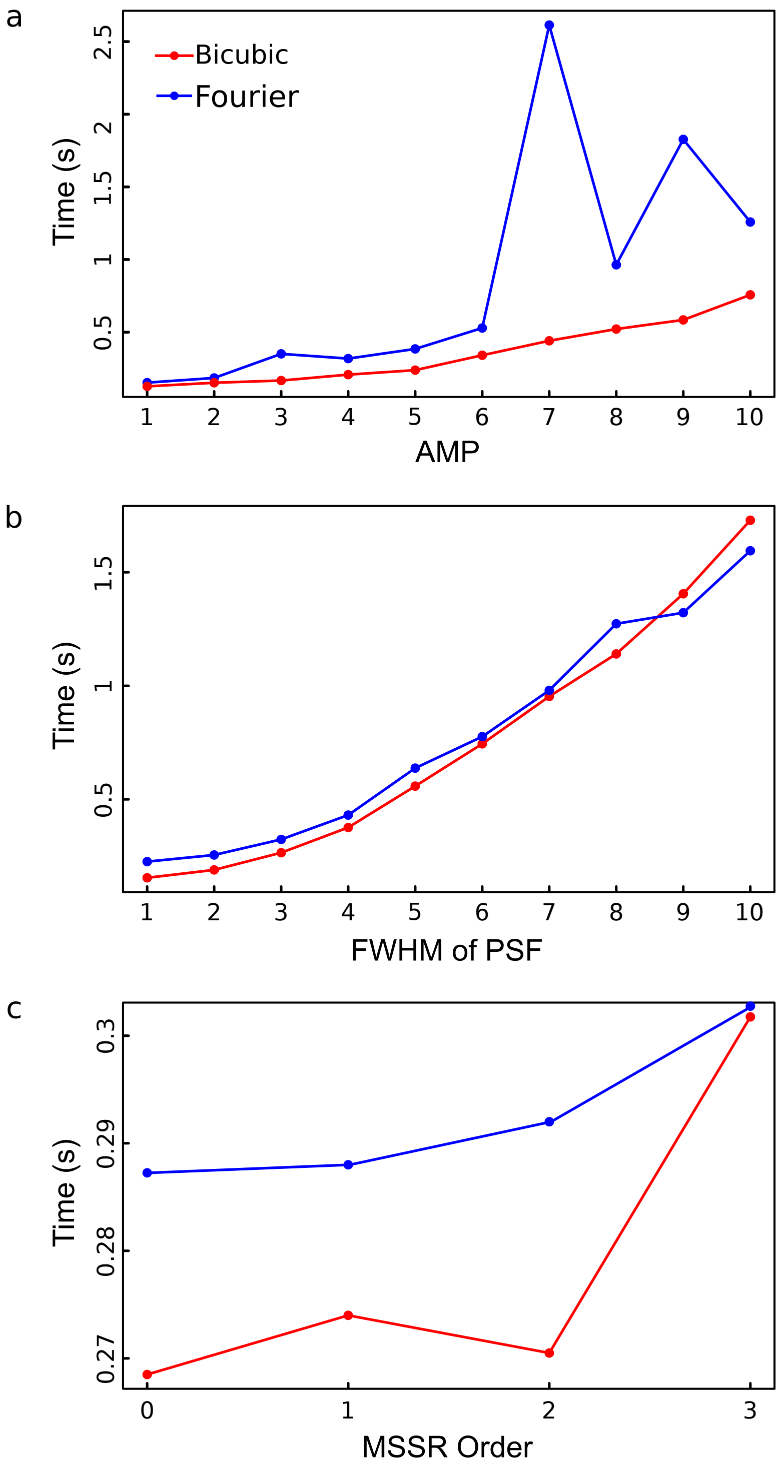

**Figure S24.** $sf - MSSR$ **processing times for bicubic and Fourier interpolations.** For each type of interpolation, the effect of the a) amplification value, the b) $FWHM$ of $PSF$ and the c) $MSSR$ order over the total run time was evaluated. All analyses were performed on a computer with an Intel Core i7-11370H CPU@3.30GHz using 16.0 GB of RAM, running Windows 10x64, with a NVIDIA GeForce RTX 3070.

Supplementary Figure S25 below shows a comparison of both interpolation methods for performing three different pixel-wise temporal functions over experimental laser lithography data (see *Online Methods* section *PSFcheck imaging*) for image reconstruction by applying temporal analysis. Future work will study the performance of both method for image reconstruction.

Since $FFT$ and $iFFT$ are used for Fourier interpolation, if an $AMP$ value different from of a power of two (i.e $AMP \neq 2^n$, $n$ integer) is applied, run times tend to be longer. In some cases, when $AMP$ is equal to a prime number, processing time extends further. For example, if $AMP = 7$ is applied, run time is 2613 ms, however, for $AMP = 10$ it decreases to $\sim$1215 ms (i.e., less than half the time). On the other hand, $MSSR$ parameters like $AMP$ and $FWHM$ of $PSF$ seem to have no significant differences between each other in terms of run time.

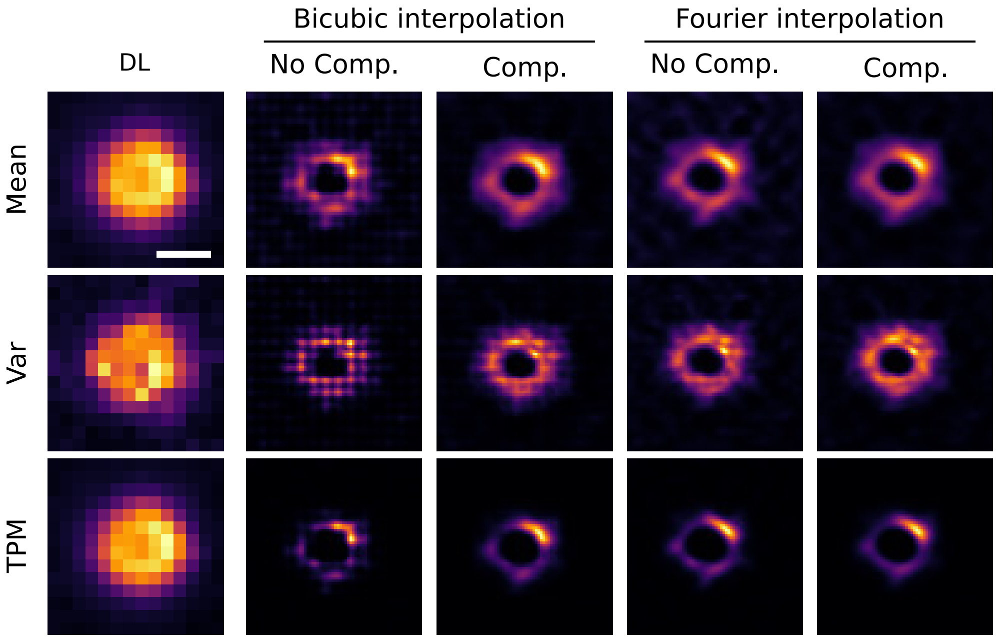

**Figure S25. Comparison between Fourier and bicubic interpolation for temporal analysis on experimental dataset.** From top to bottom: results of temporal functions of $Mean$, $Var$ and $TPM$. From left to right: first two columns are the results of bicubic interpolation with and without grid compensation, last two columns are are the results of Fourier interpolation with and without grid compensation. $MSSR$ parameters: $AMP = 5$, $FWHM$ of $PSF$ = 2, $order = 0$. Scale bar 500 nm.

### 7. SUPPLEMENTARY NOTE 7. MSSR PROCESSING TIME COMPARISON FOR DIFFERENT ANALYSIS PLATFORMS

Most of the currently available SRM algorithms (Supplementary Note 9) are implemented as $ImageJ$ plug-ins, and the majority of them report processing times of several minutes. Some of them have built-in multithreading or parallel $GPU$ capabilities, with the exception of $3B$.

In order to ensure maximum diffusion of $MSSR$ to the scientific community as a free and accessible tool, we implemented it in three different platforms: as an $ImageJ$ plug-in and as a package for $R$ and a function for $MATLAB$. We designed a simple GUI for $ImageJ$, while users with prior programming skills and the need for more statistical tools can take advantage of the analytical power of $R$ and $MATLAB$. All implementations can analyze single images or a set of images located in a specific user's system. Moreover, GPU parallel processing is an available option with $ImageJ$, especially for computationally demanding tasks.

We performed a comparative analysis of the $MSSR$ of all three platforms, evaluating the $MSSR$ order, $FWHM$ of $PSF$ and $AMP$ values over the total analysis time (Supplementary Figure S26). The data used for this study consists of a laser lithography image stack of 100 images of size 32x32 pixels [18] (see Materials and Methods). First, we evaluated the $MSSR$ parameters values for a single image from the stack. The observations indicate that $MATLAB$ implementation is the optimal.

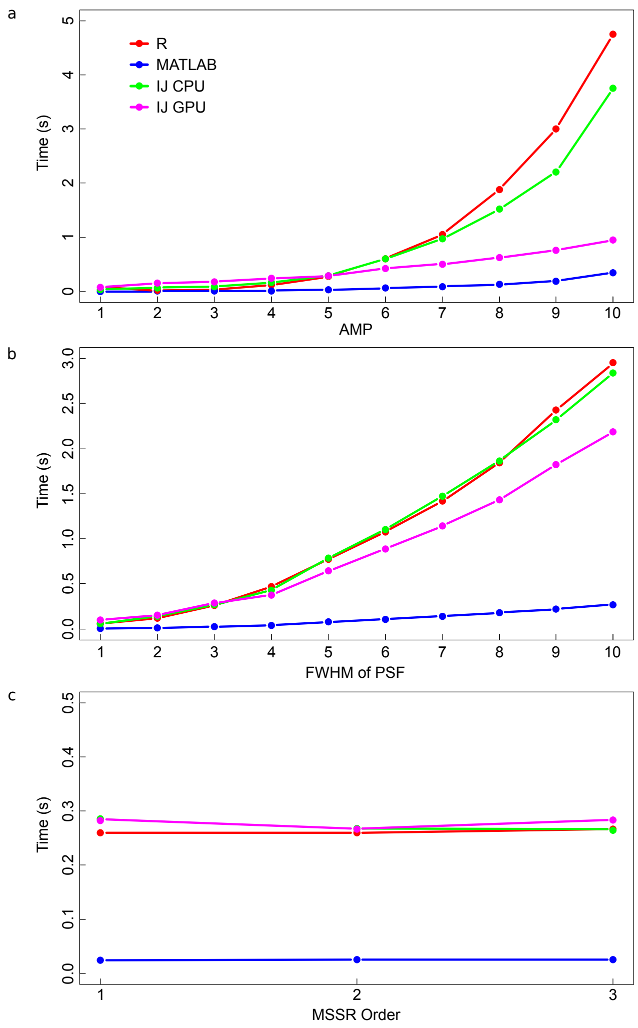

**Figure S26. Average $MSSR$ processing times for different platforms.** For each implementation, the effect of the a) amplification value, the b) $FWHM$ of $PSF$ value and the c) $MSSR$ order over the total run time was evaluated, while using GPU processing or not (if available). All analyses were performed on a computer with an Intel Core i7-7700K CPU@4.200GHz using 32.0 GB of RAM, running Windows 10x64, with a NVIDIA GeForce GTX 1070.

Next, a temporal analysis was performed using the previously obtained *MSSR* stack using optimal parameters for *MSSR* (amplification = 5, *PSF* = 3 and order = 1). The execution times obtained were proportional to those of the spatial analysis alone. In this case, the best performing platform was *ImageJ* with GPU processing enabled, showing a 30-fold improvement over *MATLAB* and a 15-fold improvement compared to CPU processing alone (Supplementary table S3).

| *MSSR* implementation | Time (s) |
|---|---|
| *R* | 22.69 |
| *MATLAB* | 2.82 |
| *IJ CPU* | 8.51 |
| *IJ GPU* | 1.31 |

**Table S3. Average *MSSR* temporal analysis processing times for different platforms.**

Even though *MSSR* can take several minutes to operate (compared to other *SRM* methods capable of increasing resolution within seconds, with the requirement of larger sets of data), it stands out among other approaches due to its ability to obtain a extended-resolved image with just one frame.

#### 8. SUPPLEMENTARY NOTE 8. SINGLE-FRAME NANOSCOPY FREE OF NOISE-DEPENDENT ARTIFACTS

The Resolution Scaled Pearson ($RSP$) coefficient and the Resolution Scaled Error ($RSE$) are global quality estimators for image reconstructions [19]. $RSP$ is an adaptation of the Pearson correlation coefficient for images, and provides a global quality assessment scored between the super-resolution image and the original one, with an ideal maximum value of 1 and a lower boundary of zero. $RSE$ is the root-mean-square-error computed between the super-resolution image and the original one; it measures the intensity differences of both inputs, where the ideal case is $RSE = 0$. We investigated the behaviour of these parameters depending on the stack size used for a $MSSR$ temporal analysis (Supplementary Figure S27).

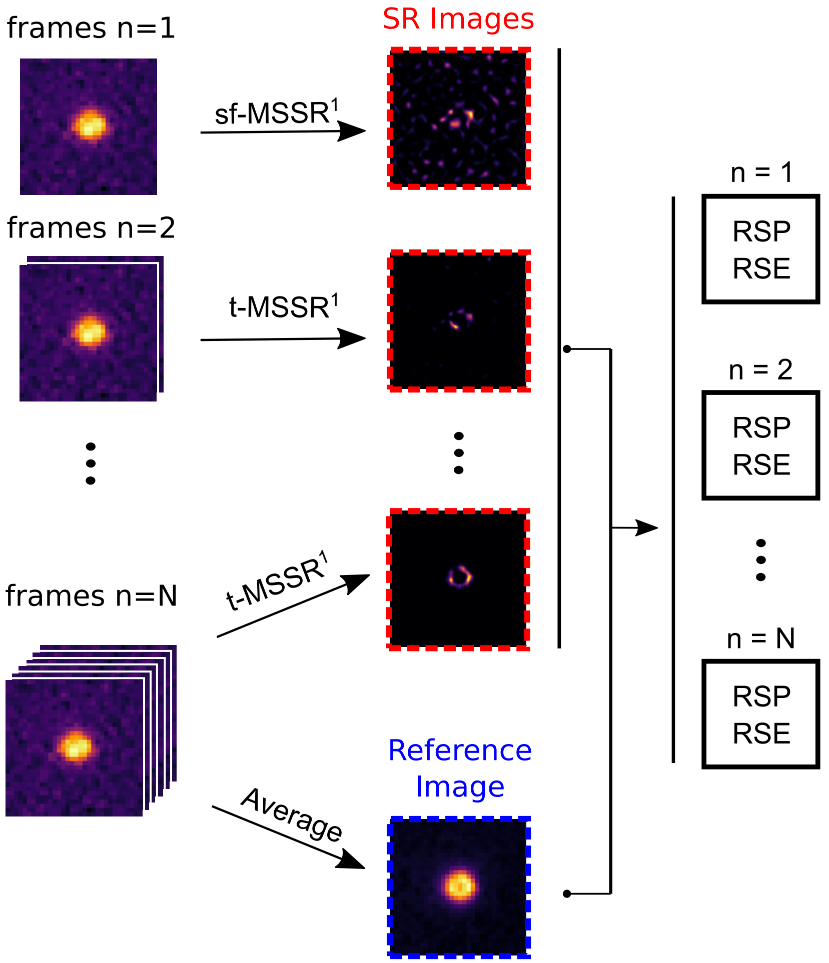

**Figure S27. Algorithm for the computation of $RSP$ and $RSE$ for super-resolved images.** Given a temporal stack of $N$ frames, $N$ super resolution images are generated by using 1, 2, 3, . . ., $N$ images as input for $t - MSSR^1$ (SR images, red). Considering the stack average to be a representative reference image, it was used to calculate the $RSP$ and $RSE$ for each of the $N$ reconstructions.

$MSSR$ operates in two stages: spatial information is first used to obtain a stack of extended-resolved frames, which are next integrated into a temporal analysis for further reconstruction quality enhancement through noise-related artifact minimization. Because digital noise is not correlated in time, its contribution to the actual signal diminishes as more frames are considered in a temporal analysis (*e.g.* averaging, for instance). However, data shows that when a certain number of frames analyzed is reached, no additional image quality is gained (Figure S28). As explained above, we generated 25 super-resolution images, each of which was obtained using 1, 2, 3, . . ., 25 sequential frames of the same field of view. The $RSP$ and $RSE$ were then computed by comparing each temporally-reconstructed image with the average of the diffraction-limited stack. The idea behind this analysis is that, although a reconstruction obtained with a limited amount of data might not completely reflect the true underlying nanoscopic structure, it does recover all the true sub-pixel information contained in the said data set. The results show that a $SNR$ lower than 5 (eq. S52) requires up to 25 images in order to provide a stable $RSP$ coefficient value. However, when the $SNR$ is high enough, an $RSP$ coefficient value of 1 is achieved with a single frame. Note that, for all $RSP$ curves, the maximum value is reached at no more than 5 images when the $SNR \geq 8.2$.

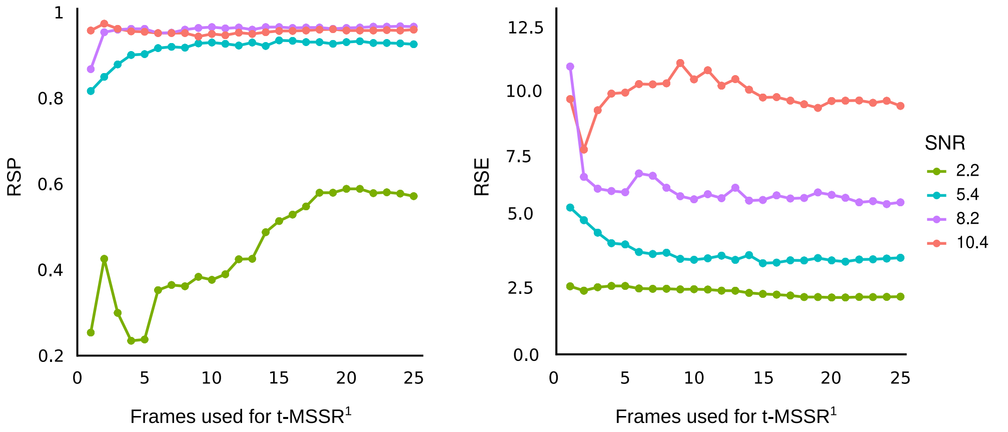

**Figure S28. Effect of $SNR$ and number of frames (temporal analysis) over $RSP$ and $RSE$.** $RSP$ (left) and $RSE$ (right) stabilize at about 25 and 5 frames for low and high $SNR$, respectively.

Based on these observations, it is advised to perform a temporal analysis over data with a $SNR$ anywhere between 3 and 8, according to the equation (https://camera.hamamatsu.com/jp/en/technical_guides/calculating_snr/index.html) [20]:

$$SNR = \frac{QE * S}{\sqrt{QE\,(S + I_b) + N_r^2}} = \frac{\text{Electrons}}{\sqrt{\text{Electrons} + \text{Read Noise}^2}}. \quad \text{(S52)}$$

In this equation:

- $S$ is the number of photons detected in a single pixel of a sCMOS camera.
- $QE$ is the quantum efficiency of the detector at the wavelength of the collected light; for this study $QE = 0.72$.
- $I_b$ is the background signal.
- $N_r$ is the camera read noise, measured in electrons (reported by the manufacturer in the specifications sheet; for our detector $N_r = 1e^-$.

To approximate $S$ and $I_b$ from the data, gray scale values (raw) were converted into average photons per pixel (https://camera.hamamatsu.com/jp/en/technical_guides/how_many_photons/index.html) [20], considering the equation

$$Photons_i = \frac{I_i - O_i}{G_i} * \frac{1}{QE}. \quad \text{(S53)}$$

where

- $I_i$ is the greyscale value of the raw image for pixel $i$.
- $O_i$ is the offset value for pixel $i$.
- $G_i$ is the gain for pixel $i$.

The offset and gain were calculated as previously reported [20].

From these data, it is possible to calculate the approximate number of photons that are detected per pixel. $S$ was considered as the average number of photons in the region where the sample of interest is located, while $I_b$ is the average signal value of the pixels that belong to the background of the image.

### 9. SUPPLEMENTARY NOTE 9. COMPARING MSSR WITH OTHER ANALYTICAL SRM METHODS

In the last decade, *SRM* development has tended towards the reduction of the needed data size to be processed to obtain a super-resolved image. Many computational methods rely on exploiting fluorophore blinking dynamics for the extraction of structural information through pixel-wise temporal operator computation. The result is a super-resolution reconstruction built from the analysis of hundreds to thousands of images collected at a static, or pseudo-static scene. However, the mandatory need of a temporal analysis to handle high spatial frequencies due to the detector's noise, restricts these methods' temporal resolution to the regime of seconds to minutes. To compare the super-resolving power of the single-frame and temporal analysis of *MSSR* with other *SRM* methods, it was tested and paired against three algorithms based on the single-molecule localization principle:

- *SRRF* (**S**uper-**R**esolution **R**adial **F**luctuation) encompasses a two-step analytical process. A single-frame analysis of local information stored within the fluorescence intensity domain provides nanoscopic resolution. Its principle is based on radiality maps theory which itself provides nanoscopic information, however, it is also sensitive to abrupt changes in local contrast when noise burdens the signal, a common scenario when studying single molecules. As result of that, it encompasses a temporal analysis aimed to reduce noise-related artifacts and to further gain resolution due to a statistical processing of a short sequence of radiality maps (about 100 images) [21]. Depending on the experimental design and the hardware architecture used, *SRRF* offers 50-150 nm of axial resolution.

- *ESI* (**E**ntropy-based **S**uper-resolution **I**maging) calculates the local- and cross-entropy to estimate the likelihood of emitters throughout an image sequence [22]. It is characterized by having effective noise-removal properties and requiring low-cost computational resources, but requires hundreds of frames for a single reconstruction. ESI is based on reconstructing super-resolved images from conventional image sequences containing random signal fluctuations and can achieve resolutions of up to $\sim$100 nm.

- *MUSICAL* (**MU**ltiple **SI**gnal **C**lassification **A**lgorithm) provides resolution enhancement by singular value decomposition to an image stack resulting on eigenimages and their respective eigenvalues [23]. Each eigenvalue describes a particular pattern on the imaged temporal stack. Signal is considered to be associated with large eigenvalues, whereas noise and background are associated with small eigenvalues. A predefined threshold divides the set of eigenimages into two ranges: space and null space. The projection of the point spread function over these spaces and their ratio is an indicator function where large values are related to the presence of fluorescence emitters and small values are related to background noise. It offers resolutions below 50 nm from the analysis of tens of images, which makes MUSICAL an effective approach for live-specimen imaging [24].

Each method's performance was evaluated at different *SNR* to test their ability to reliably underscore nanoscopic information with varying noise conditions. The quality of the reconstructions was assessed by means of computing global parameters of consistency between the SRM image and a reference image, which in this case is the corresponding diffraction-limited image.

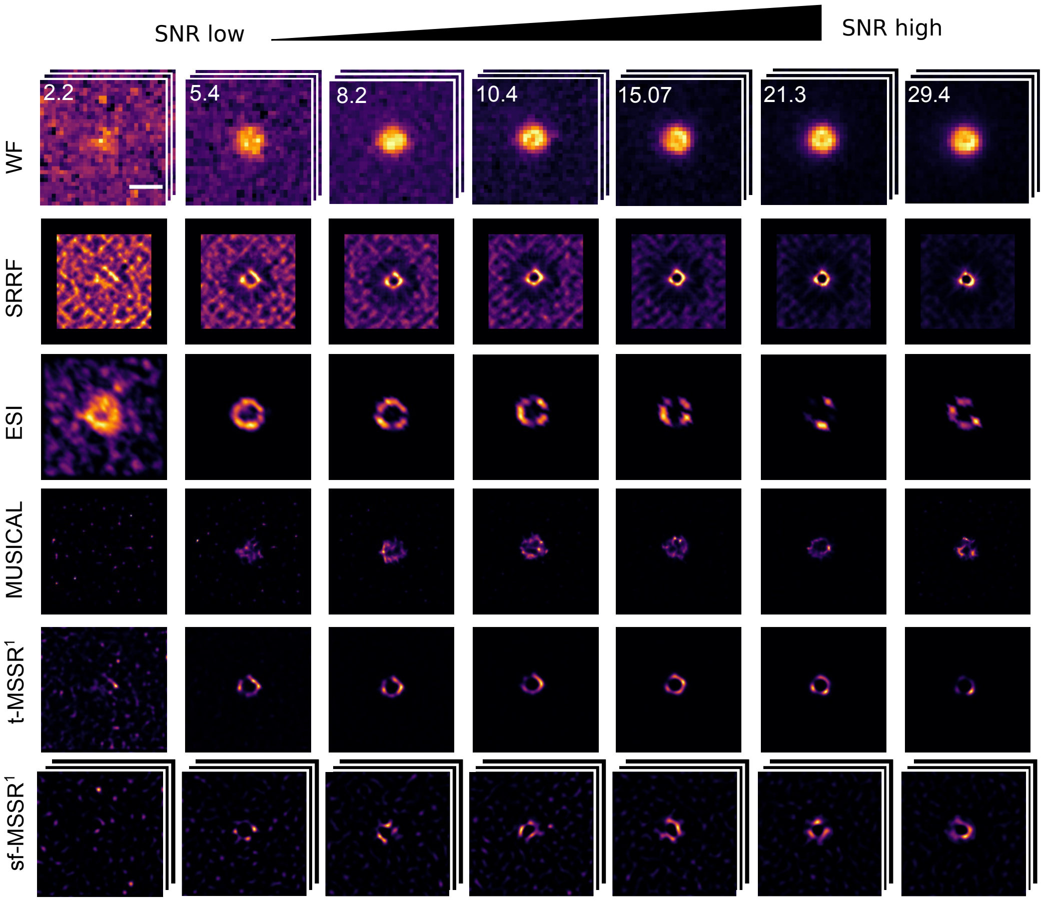

**Figure S29.** ***MSSR*** **is robust against background noise and introduces fewer artifacts when compared to other SRM methods**. Comparison of SRM images of the laser lithography sample generated by either $SRRF$, $MUSICAL$, ESI, or MSSR. The expected feature is a uniform fluorescent ring located at the center of the image with a dark background lacking fluorescence. The widefield row (WF, i.e. diffraction-limited) displays the temporal average of 100 frames. *ring radius* = 0.5, *radiality magnification* = 4, *axes in ring* = 6, rest of the parameters were taken by default; $ESI$ parameters: First iteration, *image in output* = 50, *bins for entropy* = 2, *order* = 0, Second iteration *image in output* = 25, *bins for entropy* = 2, *order* = 0; MUSICAL parameters: *emission wavelength* = 610 nm, *numerical aperture* = 1.4, *pixel size* = 117 nm, *subpixels per pixel* = 4, *threshold value* = -0.8 (chosen from the singular value plot calculated by the plugin), *alpha* = 4; $sf - MSSR$ parameters: $AMP$ = 5, $FWHM$ of $PSF$ = 4, *order* = 1; $t - MSSR$ parameters: $AMP$ = 5, $FWHM$ of $PSF$ = 4, *order* = 1, $PTF = TPM$.
Scale bar: 1 $\mu$m.

Qualitatively, fewer noise-related artifacts were introduced by $MSSR$. $MSSR$ and $SRRF$ provided SRM images with greater resemblance to the expected 'ring-like' structure imaged by SIM [25]. ESI and MUSICAL also unveiled sub-diffraction rings but with unexpected irregular or incomplete shapes. When the $SNR$ is high enough, all tested algorithms perform relatively well. Remarkably, with any tested $SNR$, $sf - MSSR^1$ provided reliable reconstructions of comparable quality to those approaches requiring a temporal analysis and resolved the underlying structure while introducing visually fewer artifacts than any of the other methods.

Since our data were acquired with a sCMOS camera, we tested the ability of a recently published sCMOS-specific noise correction algorithm [26] to improve quality of the super-resolution images, independently of the SRM method used. The effect of this noise-reducing step over the global quality of reconstruction was evaluated in terms of global image quality (Supplementary Figure S30). For the case of $SRRF$, $RSP$ values increased when the $SNR$ was low, while displaying visually fewer artifacts; the other methods showed no significant difference after denoising the images. This suggests that $SRRF$ is highly sensitive to noise, while $MSSR$, $ESI$ and $MUSICAL$ are less sensitive. Nonetheless, $MSSR$ and $SRRF$ were able to provide a more reliable ring-like structure than that of $MUSICAL$ or $ESI$. Surprisingly, $ESI$ displayed the highest $RSE$ values after denoising. Regardless of whether the input data was corrected or not, the $RSP$ and $RSE$

values for *MSSR* were very similar. Overall, *MSSR* did not perform much better after data was noise-corrected, while the other tested methods benefited greatly, showing a significant improvement in reconstruction quality. These observations indicate that *MSSR* highly resiliant to noise and can perform very well even in low light conditions.

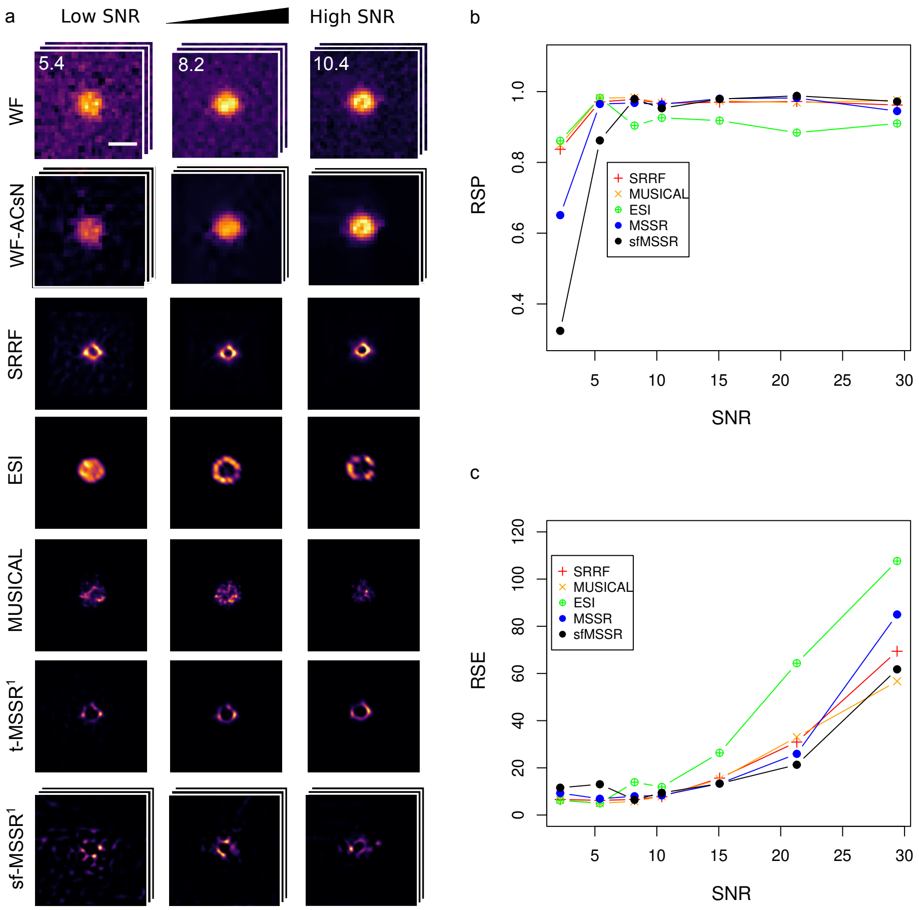

**Figure S30. MSSR provides no further gain in** $RSP$ **and** $RSE$ **after image denoising. a)** Image reconstructions for data with $SNR = 5.4$, 8.10 and 10.4 after noise reduction via ACsN. A representative stack of raw images (RAW) and its noise-cleaned version after ACsN (WF-ACsN) are shown. Subsequent reconstructions ($SRRF$, $MSRR^1$, $ESI$ or $MUSICAL$) were performed using 100 of these noise-cleaned images (WF-ACsN). Single Frame $MSSR$ row (sf-$MSSR^1$) shows a representative image for the analysis of a single frame (only spatial analysis) by $MSSR$ with the cleaned data. **b)** $RSP$ and **c)** $RSE$ values calculated for the super-resolution reconstructions in a) over a wider range of $SNR$, including the representative example of single frame MSSR. *ring radius* = 0.5, *radiality magnification* = 4, *axes in ring* = 6, rest of the parameters were taken by default; $ESI$ parameters: First iteration, *image in output* = 50, *bins for entropy* = 2, *order* = 0, Second iteration *image in output* = 25, *bins for entropy* = 2, *order* = 0; MUSICAL parameters: *emission wavelength* = 610 nm, *numerical aperture* = 1.4, *pixel size* = 117 nm, *subpixels per pixel* = 4, *threshold value* = -0.8 (chosen from the singular value plot calculated by the plugin), *alpha* = 4; $sf - MSSR$ parameters: $AMP$ = 5, $FWHM$ of $PSF$ = 4, *order* = 1; $t - MSSR$ parameters: $AMP$ = 5, $FWHM$ of $PSF$ = 4, *order* = 1, $PTF = TPM$.
Scale bar: 1 $\mu$m.

Finally, we use as input data a version of the images where each pixel is represented as the approximate number of photons that reached the detector at the time of acquisition (see Methods). The result of using this input data (photon images) is similar to that of using ACsN-cleaned data (Supplementary Figure S31). That is, calculating the average number of photons per pixel increases the *RSP* values for *SRRF* processing. This can be explained because the formula to obtain the photons per pixel is equivalent to the pre-correction step used by ACsN and other noise-correction algorithms prior to the noise reduction refinement. This suggests that by knowing the response function of the detector and thereby performing a pre-correction to the raw data, one can improve the quality of the images.

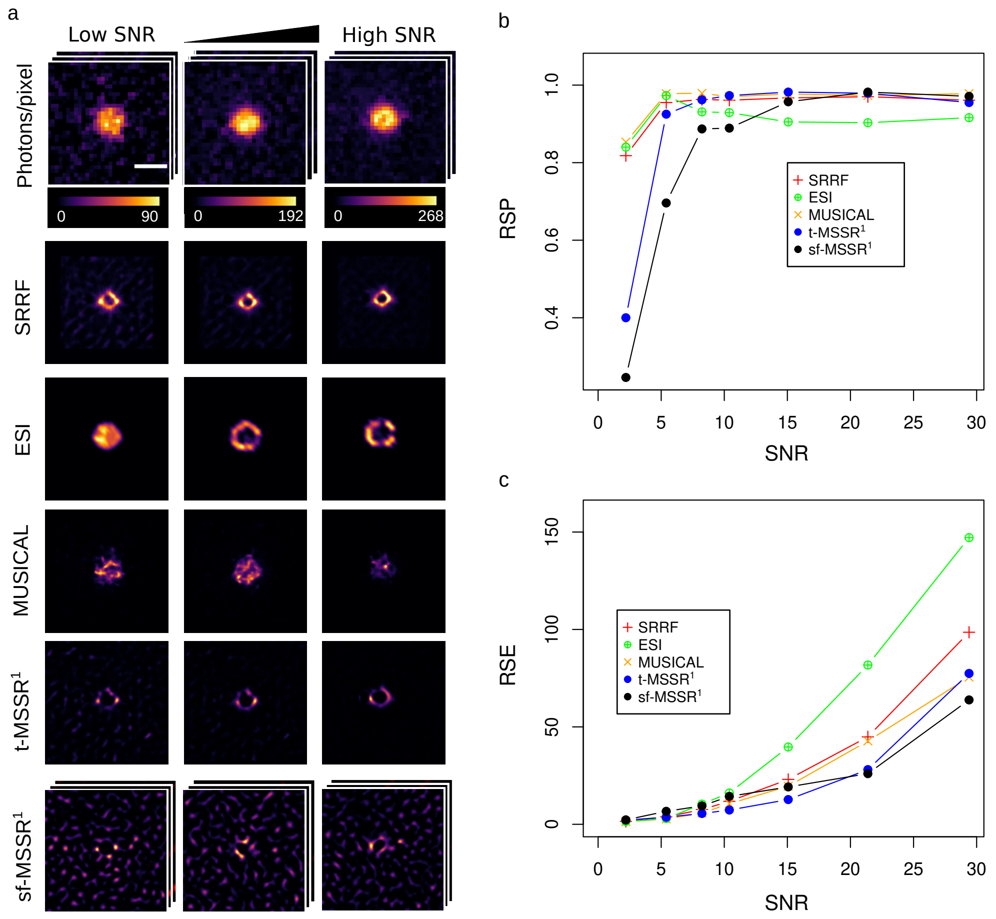

**Figure S31. Visual examples, $RSE$ and $RSP$ of reconstructions using image data in photons per pixel. a)** Image reconstructions for data with $SNR = 5.4$, 8.10 and 10.4 after pixel intensity was converted to photons per pixel (see Supplementary Note 8). Each image is displayed to show its full intensity range. Images converted to photons/pixel (first row) include a pseudo-color scale indicating the range of photons/pixel of each image. Their corresponding signal-to-noise ratio ($SNR$) is indicated on the upper left corner. Subsequent reconstructions ($SRRF$, $MSRR^1$, $ESI$ or $MUSICAL$) were performed using 100 of these converted images. Single Frame $MSSR$ row (sf-$MSSR^1$) shows a representative image for the analysis of a single frame (only spatial analysis) by $MSSR^1$ with the converted data. **b, c)** Resolution Scaled Pearson ($RSP$) coefficient (b) and Resolution Scaled Error ($RSE$) (c), computed for the super-resolution reconstructions in a). *ring radius* = 0.5, *radiality magnification* = 4, *axes in ring* = 6, rest of the parameters were taken by default; $ESI$ parameters: First iteration, *image in output* = 50, *bins for entropy* = 2, *order* = 0, Second iteration *image in output* = 25, *bins for entropy* = 2, *order* = 0; MUSICAL parameters: *emission wavelength* = 610 nm, *numerical aperture* = 1.4, *pixel size* = 117 nm, *subpixels per pixel* = 4, *threshold value* = -0.8 (chosen from the singular value plot calculated by the plugin), *alpha* = 4; $sf - MSSR$ parameters: $AMP$ = 5, $FWHM$ of $PSF$ = 4, *order* = 1; $t - MSSR$ parameters: $AMP$ = 5, $FWHM$ of $PSF$ = 4, *order* = 1, $PTF = TPM$.
Scale bar: 1 $\mu$m.

#### 10. SUPPLEMENTARY NOTE 10. EMPIRICAL DEMONSTRATION OF MSSR APPLICABILITY TO A WIDE RANGE OF FLUORESCENCE BIOIMAGING MODALITIES

The widespread use of fluorescence microscopy has led up to the development of a myriad of imaging protocols and acquisition systems. The *MSSR* principle stems from the analysis of local information of a single image. It provides nanoscopic resolution by means of solving an operator that incorporates information from both the spatial distribution of fluorescence (a spatial range covering the PSF) and range of local intensities (also known as the level of gray values) (Supplementary notes 3, 5 and 6). Such lower requirement for information makes *MSSR* suitable for its use at a wide range of applications of fluorescence microscopy, which are not necessarily limited to the realm of super-resolution microscopy.

To illustrate the versatility of the *MSSR* approach, its ability to provide nanoscopic information on a variety of biological and experimental systems (which have been previously reported by other SRM approaches) was tested. Common experimental scenarios of fluorescence microscopy are discussed after post-processing with *MSSR*; these include but are not limited to immunofluorescence imaging of fixed cells by epifluorescence microscopy (Supplementary Figure S32), live-cell imaging using permeable dyes with total internal reflection fluorescence microscopy (Supplementary Figure S33, S34), single particle tracking (Supplementary Figure S36), live-cell imaging of microtubule dynamics (Supplementary Figure S37), colocalization microscopy (Supplementary Figure S38), tissue live cell imaging - laser scanning confocal microscopy (Supplementary Figure S39).

##### 10.1. Nanoscopic organization of viral replication machineries

Viral replication is orchestrated at nanoscopic scales within machineries which perturb cellular architecture to ensure the production of viral progeny. The onset of rotavirus assembly occurs at viroplasms ($VP$s), which are protein coacervates shaped by both viral and cellular components that, when visualized by electron microscopy, resemble complex electrodense structures [27].

Supplementary Figure S32 shows diffraction-limited TIRF images of a rotavirus $VP$. Note that, regardless of the protein distribution analyzed, the $VP$ resembles diffraction-limited clumps of fluorescence shaped by viral proteins. The $NSP2$ protein, denoted by red channel, was selected to ensure the detection of $VP$s on infected cells and served as a point of reference to compare the relative distribution of the other viral proteins within the $VP$.

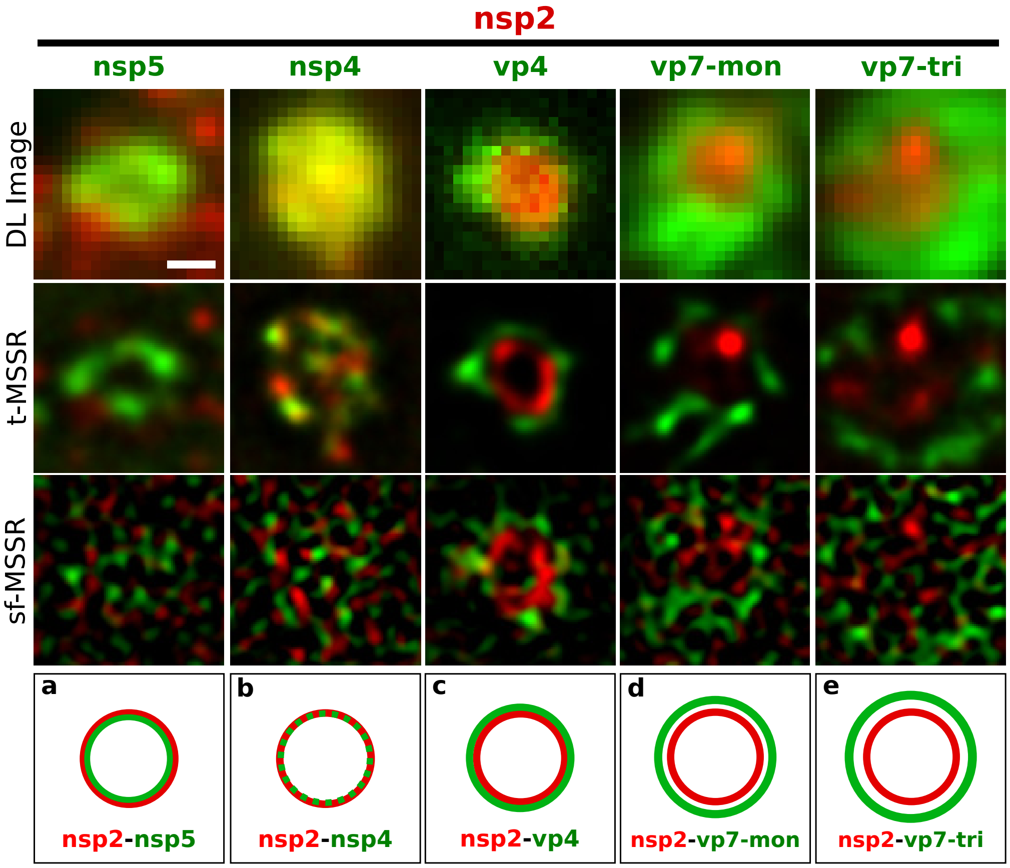

**Figure S32. Relative distribution of viral proteins in rotavirus viroplasms ($VP$s) visualized by $MSSR$ analysis.** RRV-infected MA104 cells (6 hpi) were fixed and processed for immunofluorescence microscopy. The $NSP2$ protein (denoted by the red channel for all panels) was used for $VP$ detection on infected cells and served as point of reference in order to compare the relative distribution of the other viral proteins in the $VP$ (indicated by the green title of each column). Rows from top to bottom: diffraction-limited (DL) images of the $VP$s; $t - MSSR$, reconstructions provided by a temporal analysis of 100 diffraction-limited images per $VP$; $sf - MSSR$, single-frame analysis reconstructions. a-e) Proposed models of the viral proteins distribution in the $VP$, based on the super-resolution data, as reported in [28]. $MSSR$ parameter: $AMP$ = 5, $FWHM$ of $PSF$ = 2.79 (green channel) and $FWHM$ of $PSF$ = 3.31 (red channel), $order$ = 1, $PTF = Mean$.
. Scale bar 500 nm.

In stark contrast with the observations made through visual inspection of the diffraction-limited images of the $VP$ (Supplementary Figure S32 – upper row), it appears that viral proteins are layered as concentric nanoscopic rings of proteins within the rotavirus $VP$ [28]. $NSP5$ and $NSP2$ act as core-forming scaffolding elements, which coexist at the inner layers of the $VP$ alongside with $NSP4$, followed by layers of $VP1$, $VP2$, $VP6$, $VP4$ and $VP7$ arranged in an inside-out fashion. These findings were discovered by using 3B-ODE SRM by means of analyzing a temporal stack of hundreds of diffraction-limited images with the help of using colocalization microscopy, TIRF imaging and a dedicated cluster for parallel computing (with a mean processing time of one day per super-resolution micrography) [28, 29].

With $MSSR$, we were able to visualize the nanoscopic organization of the rotavirus $VP$ by virtue of analyzing single diffraction-limited images in a matter of seconds using a consumer-grade computer. Viral proteins within the $VP$ milieu showed a layered organization, resembling the one observed by Garcés in [28]. Supplementary Figure S32 shows distributions of the viral proteins $NSP2$, $VP4$, $NSP5$, $NSP4$ and $VP7$ in its monomeric and trimeric form. The innermost core of the $VP$ was found to be enriched by $NSP2$, $NSP5$ y $NSP4$ proteins, followed by layers of $VP4$ in the middle and $VP7$ in its monomeric and trimeric form in the outermost side of the $VP$.

For the sake of comparison with the results obtained in [28] using $3B - ODE\ SRM$, the same $VP$s as in Supplementary Figure S32 were processed with both the single-frame ($sf - MSSR$) and pixel-wise temporal analysis ($t - MSSR$) strategies. Although the same concentric pattern of viral protein arrangement within the $VP$ was underscored using both approaches, differences in the layer structure were observed, such as the thickness and continuity of the layer. This is because, in $sf - MSSR$, the output image accounts only for the signal corresponding to the set of emitting fluorophores captured in the acquisition time interval of the input image analyzed, whereas the pixel-wise temporal analysis generates an ensemble averaged reconstruction with nanoscopic resolution.

##### 10.2. Live-cell extended-resolution imaging

*MSSR* was challenged for live cell extended and super-resolution imaging by means of studying the acrosomal exocytosis (*AE*), a unique secretory event which results from fusion events between the plasma membrane and a specialized vesicle called acrosome located in the sperm head [30].

The nanoscopic dynamics of the *AE* were visualized by following the spatio-temporal dynamics of SiR-actin, a fluorescent probe which preferentially fluoresces when bound to F-actin (green), and FM4-64, which underlines cellular membranes (magenta) (Supplementary Figure S33). In agreement with previous findings [31], *MSSR* facilitates the visualization of complex reorganization events at the plasma membrane, at the inner and outer acrosomal membranes and at the cortical actin cytoskeleton located underneath the acrosomal vesicle, as result of the *AE* ( Supplementary Figure S33).

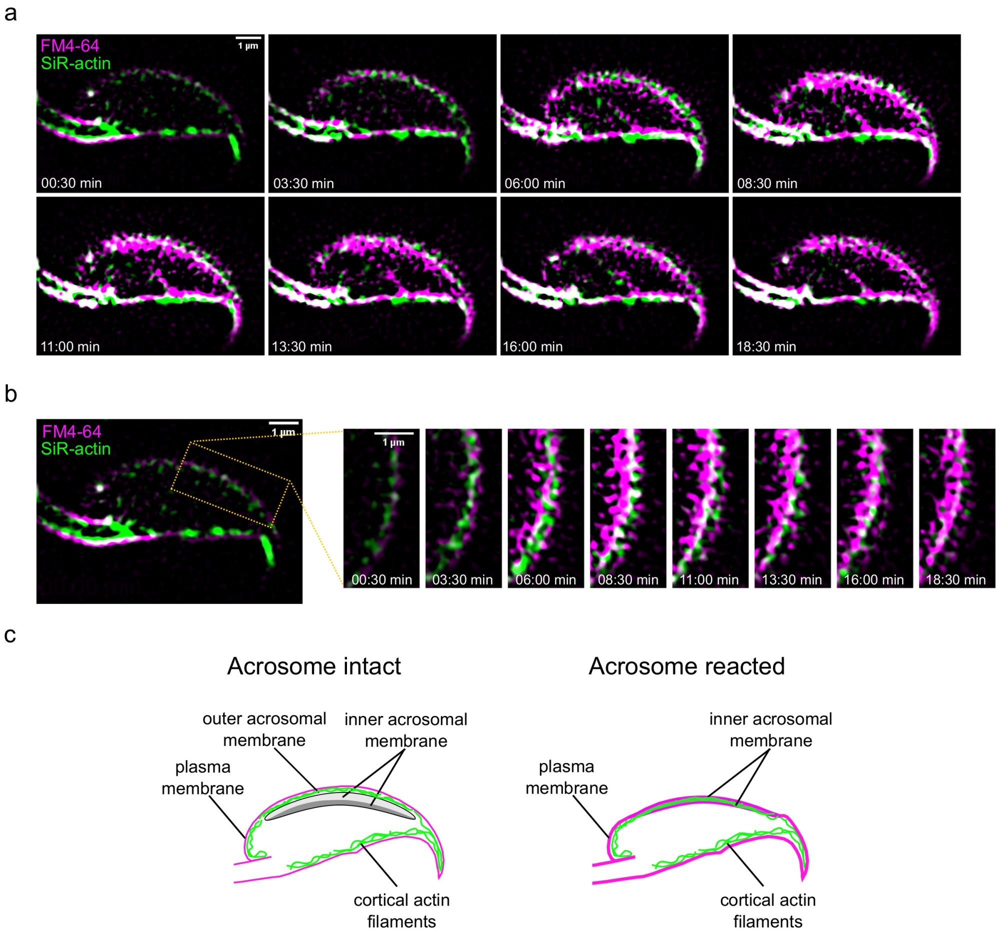

**Figure S33. Nanoscopic, single-frame, live-cell imaging of acrosomal exocytosis in mouse sperm.** F-actin (stained with SiR-actin) is shown in green and plasma membrane (stained with FM4-64) is shown in magenta. Capacitated mouse sperm were loaded with both dyes and immobilized on concanavalin A-coated coverslides. **a)** $sf - MSSR^0$ facilitates the visualization of the course of spontaneous acrosomal exocytosis. **b)** Higher magnification of sperm membrane experiencing fenestration events. **c)** Schematics of acrosomal exocytosis. At the beginning, only the plasma membrane can be seen through FM4-64 dye (left). The plasma membrane as well as the outer acrosomal membrane are lost as the acrosomal exocytosis develops. As a result, FM4-64 (present in the registration medium) now stains the inner acrosomal membrane (right). $sf - MSSR$ parameters: $AMP = 5$, $FWHM$ of $PSF = 5$, $order = 0$.

Several fenestration events were observed occurring at and within the plasma and acrosome membranes (Supplementary Figure S33a and Movies S4 and S5), showing complex millisecond dynamics (Supplementary Movies S6 and S7). At the beginning of the *AE*, the FM4-64 fluorescence was restricted to the plasma membrane which was visible above the F-actin cytoskeleton (Supplementary Figure S33a, Movies S5 and S6). At the end of the *AE*, the plasma membrane and the outer acrosomal membrane are lost, exposing the inner acrosomal membrane, hence providing new binding sites for FM4-64 (Supplementary Figure S33a, Movies S5 and S7) [32]. As a consequence, a notorious increase of FM4-64 fluorescence was recorded close to the cortical actin cytoskeleton (Supplementary Figure S33 b and c).

If we compare the $SRM$ images obtained from either the single-frame or temporal analysis (of the same pseudo-static scene) approaches, as in [31], the overall observations that arise are identical (compare Supplementary Figures. S33 and S34; compare Supplementary Movies S4 and S5 with Supplementary Movies S8 and S9). In either case, the rise in FM4-64 fluorescence, F-actin dynamics and membrane fenestration events can be observed. Nonetheless, an advantage of using the single-frame approach is that it implies much less sample manipulation and causes less phototoxicity, hence damage to the sample is reduced and the overall integrity of the experiment is preserved. Moreover, $sf - MSSR$ allows observation of milliseconds dynamics of an overall process that takes minutes to occur; this information is usually lost when hundreds of images are necessary to derive a single super-resolution micrography (Supplementary Figure S34) [31].

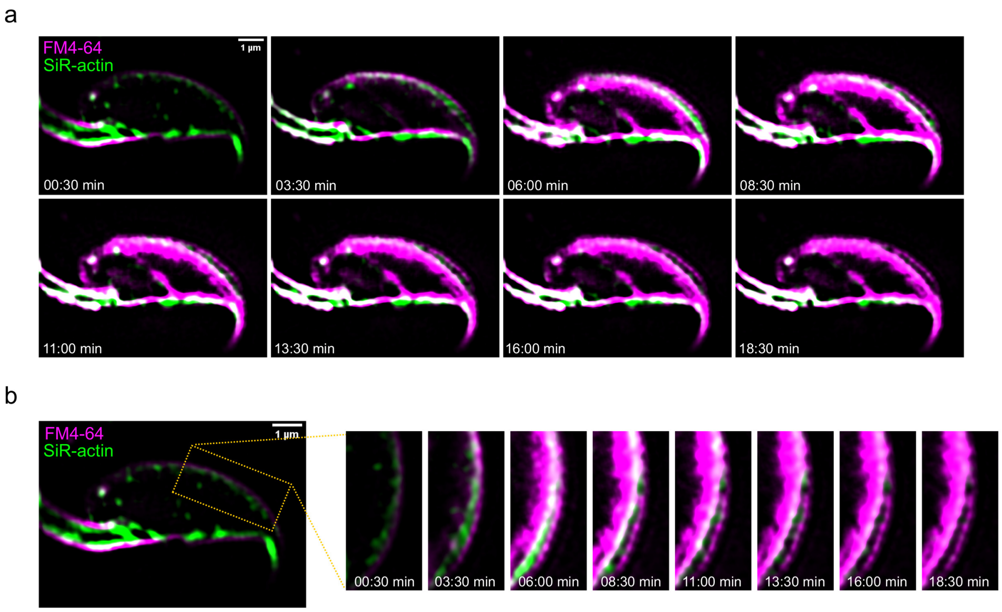

**Figure S34. Nanoscopic, single-frame, live-cell imaging of acrosomal exocytosis in mouse sperm.** F-actin (stained with SiR-actin) is shown in green and plasma membrane (stained with FM4-64) is shown in magenta. Capacitated sperm were loaded with both dyes were inmobilized on convanavalin A-coverslides for imaging. **a)** By using the 100 frame $t - MSSR^0$ analysis, it is possible to visualize the course of spontaneous acrosomal exocytosis. **b)** Higher magnification of sperm membrane fenestration. $t - MSSR$ parameters: $AMP = 5$, $FWHM$ of $PSF = 5$, $order = 0$, $PTF = Mean$.

In summary, $MSSR$ can unveil reliable nanoscopic information at the minimal speed requirements to record a single fluorescence scene.

##### 10.3. Single-particle tracking is enhanced by MSSR

Over recent years, single-molecule imaging together with super-resolution microscopy have contributed to our understanding on how molecular motion at the nanoscale level occurs [33–35]. Although single-molecule techniques have enabled us to acknowledge more details about the intricate biology of the cell such as movement of molecular motors or lipid diffusion dynamics at the cell membrane, there is still much to contribute to the fundamental understanding on how these different phenomena work at the cell extent.

Single Particle Tracking ($SPT$) has been successfully applied to a variety of different systems ranging from tracking the 2D motion of single lipids in well-defined [36–38], synthetic lipid bilayers up to the motion of cells in entire 3D organoids and embryos [39–41]. Despite that, $SPT$ is an imaging modality that requires special microscopy features that help to achieve exceptional temporal and spatial resolution in order to provide faithful data of the biological process of interest. Successful $SPT$ relies on the compliances in general aspects such as detection (good signal to noise ratio and frame ratio), localization (quantum efficiency, high-magnification, subpixel reconstruction, single-emitter localization) and linking the trajectories throughout the frames; the analysis of the resulting data can provide information about the motion of the set of labelled molecules that are being studied.

In the majority of live-cell imaging applications, tracking single particles demands short exposure times to capture and restore meaningful particle motion while avoiding cellular phototoxicity. Moreover, each fluorescent molecule has limited capabilities to produce photons. When the signal of a single fluorophore is received and digitalized by the camera, the photons arrive at random positions given by the distribution defined by the $PSF$. Therefore, the precision limit is given by the central limit theorem as $\sigma\sqrt{N}$ where $N$ is the number of photons, that is, by shot noise. Any tested $SPT$ algorithm performs reliably over an experimentally established $SNR$ lower boundary of 5 [42].

Apart from shot noise, the accuracy can worsen in the presence of background noise, delivered by either autofluorescence or detector's noise. Yet another problem arises from particle-to-particle proximity, which limits the ability to resolve them as single entities. $MSSR$ has shown denoising properties, providing nanoscopic information at very low $SNR$ (Figure 5 main document). To test whether $SPT$ can be improved by $MSSR$ preprocessing, our SRM method was scrutinized by the Single Particle Tracking Challenge ([42]; http://bioimageanalysis.org/track/).

Representative image datasets with exact ground truth data were used as a reference to assess the implications of preprocessing with $sf - MSSR^1$ before $SPT$, considering two main factors affecting tracking performance, namely, the number of particles within the imaging field and their signal relative to the noise (Supplementary Figure S35). Three classes of tracking algorithms were tested in TrackMate v7.6.1[43]: (i) $LAP$: the LAP framework for Brownian motion [44], (ii) $LM$: a linear motion tracker based on a Kalman filter [45, 46] and (iii)$NN$: a tracker based on Nearest neighbors (Supplementary Figure S36) [47–49]. Particles were identified using the $LoG$ detectors considering a particle diameter of 2 pixels. For $SNR = 2$, $LoG$ detection results were dominated by noise, hence, the histogram of detection quality was used to select a threshold that yielded the expected particle number of the dataset. For $SNR > 2$, the detection quality histogram was bimodal, the threshold was select at the dip between distributions. Tracking parameters were $LM$ (*initial search radius* = 10, *search radius* = 7, *max frame gap* = 3); $LAP$ (*max linkage distance* = 7, *max gap-closing distance* = 10, *max frame gap* = 3); $NN$ (*max search distance* = 10).

The tracking accuracy over raw or $sf - MSSR^1$ -preprocessed data was assessed in Icy 2.4 [50], as indicated in [51], following four indicators that range from 0 (worst) to 1 (best): $\alpha$ and $\beta$ indicate the overall degree of matching between ground truth and estimated trajectories; The Jaccard similarity between tracks quantifies the correspondence between the ground truth tracks and those returned by the tested tracking algorithm; Finally, the Jaccard similarity between detections quantifies how well the ground truth detections are matched by the particles detected by each tracking algorithm. Tracking performance was computed by pairing the ground truth tracks vs either $LM$, $LAP$, or $NN$ tracks, considering a maximum pairing distance between detections of 5. Supplementary Figures S35 and S36, show that, at any tested set of conditions ($SNR$, particle density, $SPT$ algorithm), $sf - MSSR^1$ significantly improves overall tracking performance.

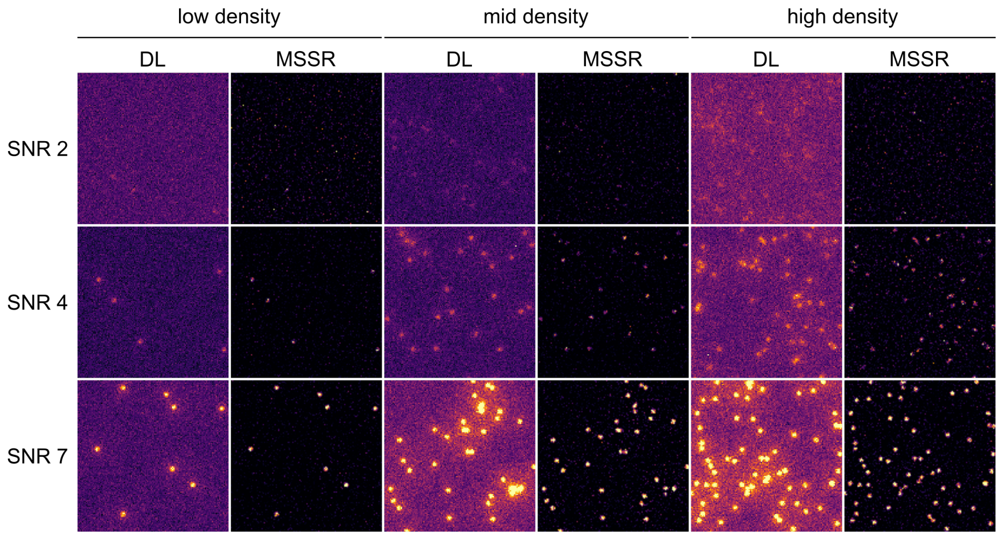

**Figure S35. Example images of the vesicles dataset before and after sf-MSSR1 processing.** $DL$ images and $sf - MSSR^1$ ($AMP = 1$, $FWHM$ of $PSF = 11.4$, $order = 1$) results for of diffusing vesicles from the "Single Particle Tracking Challenge" generated at three different $SNR$ : 2, 4, 7 and in three density levels of sub-diffraction particles: $low$, $mid$, $high$: 100, 500, 1000 particles per imaging field.

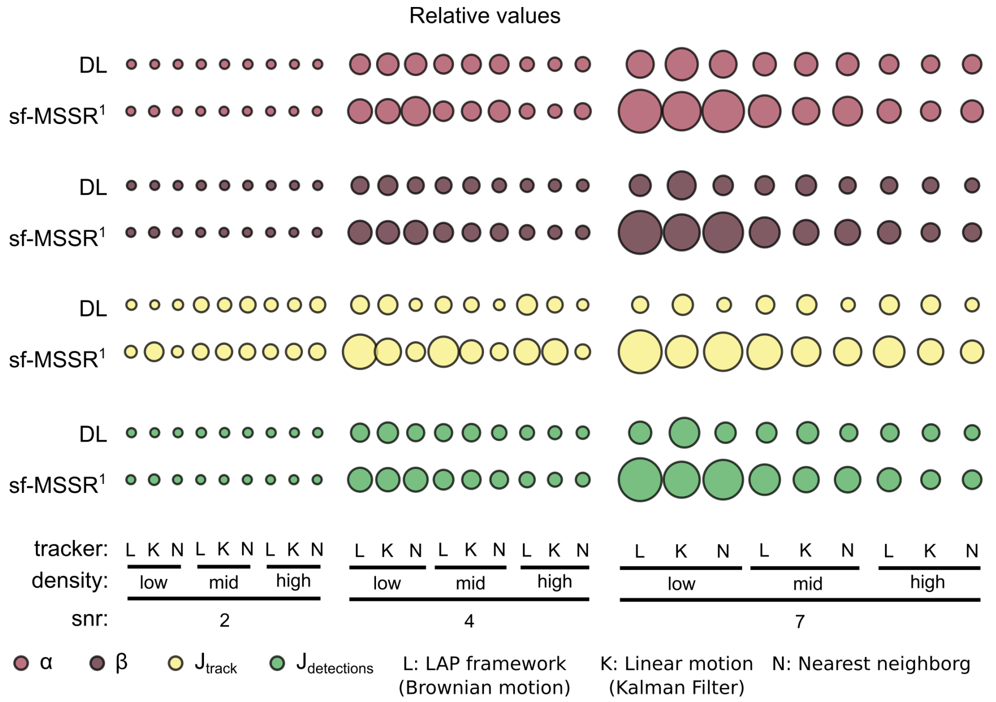

**Figure S36. sf-MSSR1 improves single particle tracking performance.** Tracking performance study using a data set of diffusing vesicles from the "Single Particle Tracking Challenge" generated at three different $SNR$: 2, 4, 7 and in three density levels of sub-diffraction particles: $low$, $mid$, $high$: 50-100, 100-500, 1000 particles per imaging field. The proficiency of three trackers *LAP framework*, *linear motion tracker*, *nearest neighbor* was tested over raw or pre-processed data with $sf - MSSR^1$ ($AMP = 1$, $FWHM$ of $PSF = 11.4$, $order = 1$). Tracking performance was assessed with the "tracking performance evaluation tool" deployed at Icy Icy 2.4 [50]. The size of the bubble represents the normalized tracking accuracy for: the overall degree of matching between ground truth and estimated trajectories ($\alpha$ and $\beta$), the Jaccard similarity between tracks ($Jt$), and the Jaccard similarity between detections ($Jd$). 'Big' indicate high estimates (ideal 1), whereas 'small' bubbles indicate poor estimates (worse 1).

##### 10.4. Live-cell imaging of microtubule dynamics

Microtubules are hollow polymer structures that comprise one of the three major components of the cytoskeleton in eukaryotes. These structures are involved a variety of cellular processes including organelle trafficking, cell migration, chromosome segregation and cell division, among others [52]. Microtubules undergo constant restructuring and switch dynamically between growing and shrinking phases, which is known as dynamic instability and is regulated by the hydrolysis of $GTP$ by $\beta$-tubulin [53].

Microtubule dynamics have been extensively investigated for their role in the regulation of several biological processes, primarily during cell division and differentiation. The highly conserved end-binding family of proteins ($EBs$) are master regulators of microtubule dynamics that aggregate at the tip in mammalian cells. These proteins recognize the minus and plus ends of microtubules, and are in charge of modulating and coordinating microtubule polymerization/depolymerization in live cells, as well as recruiting other proteins to interact with each other and orchestrate complex cytoskeleton-related cellular processes, such as cell and organelle shaping, migration and neuronal development [54–56].

Microtubules are polymeric cylindrical structures of about 25 nm [57] in diameter, which has hindered our ability to resolve them due to the diffraction limit of light. $EB$s are a widely used tool to image microtubule dynamics in live cells. However, despite their high specificity for labeling the tip of these cytoskeletal components, spatial resolution has been a limiting factor when attempting to track these comet-like moving particles. As discussed in the previous section, $SPT$ approaches have contributed towards our understanding of molecular motion at the nanoscale. However, our ability to faithfully track particles in a crowded molecular environment is commonly hindered by the overall contrast and $SNR$ of microscopy images. In the field of live-cell imaging, where low exposure times are desired in order to avoid cellular phototoxicity, $MSSR$ represents an auxiliary tool for the enhancement of the signal of fluorescence images, reducing the effect of background noise and increasing the contrast of moving particles.

As a proof of principle, live porcine kidney (LLC-PK1) cells stably expressing $EB3$ fused to mEmerald were imaged to investigate the growth dynamics of microtubules at the plus end. Supplementary Figure S37 shows a comparison between a diffraction-limited scene of live LLC-PK1 cells displaying microtubule dynamics and its $sf - MSSR$ extended-resolved counterpart. Note the visible gain in image contrast and background noise suppression, which greatly enhances the ability to track individual comets through time. Panel (a) shows a $2\times$ enlargement of a microtubule organization center ($MTOC$), which displays a clearer image of how microtubule polymerization emerges from this cellular structure when compared to the diffraction-limited counterpart. Similarly, panel (b) provides a zoomed view of microtubules polymerizing in a left-to-right sense at the edge of a cell, showing a considerable increase in contrast and comet resolution as the green fluorescent fusion of $EB3$ travels from the diffraction limited half to the $sf - MSSR$ extended-resolved side (see Supplementary Movies S10 and S11).

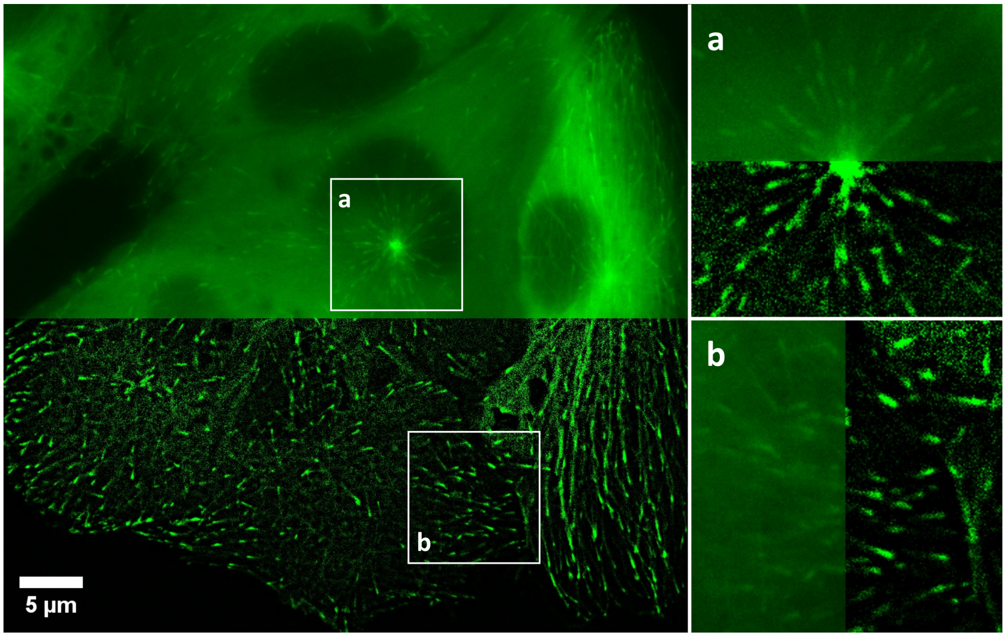

**Figure S37. Nanoscopic, single-frame, live-cell imaging of microtubule dynamics in LLC-PK1 cells.** LLC-PK1 cells stably expressing mEmerald-EB3 were cultured and imaged using a sCMOS sensor (Hamamatsu, C15440-20UP) and a 100×/1.47 NA oil-immersion objective in a Zeiss Celldiscoverer 7 microscope with $CO_2$ and temperature control set at 5% and 37°C, respectively. Fluorescence excitation was provided by a 488 nm laser at 1% laser power and emission light ($\lambda_{em}$ = 510 nm) was collected using a FITC filter. The comet-like structures depict the activity of plus-end-binding protein 3 (EB3) as it coordinates microtubule growth dynamics. Right side contains two 2× enlargements of the areas depicted by the white squares. **a)** A microtubule organizing center (*MTOC*), from which microtubule growth emerges. **b)** Microtubule dynamics at the edge of a cell with a left-to-right growth sense (see Supplementary Movie S10). Stable cell lines were generated and provided by Michael W. Davidson. $sf - MSSR$ parameters: $AMP = 3$, $FWHM$ of $PSF = 30$, $order = 0$.

##### 10.5. Dynamic colocalization with nanoscopic resolution

When looking for molecular interactions, a challenge for $SPT$ emerges as single molecules often have nanoscopic dimensions. Observing colocalization implies looking at sub-pixel resolution, hence interpretations must be elaborated bearing in mind restrictions due to PSF size and sampling. We applied $MSSR$ to DNA curtain assays to analyze interaction of DNA-bound fluorescent proteins. Here, a self-assembling polypeptide ($C - S_{10} - B$, labeled in green) binds to a DNA that has been pre-decorated with $CRISPR - dCas12a$ (labeled in magenta) via multiple $CRISPR - RNA$s ($crRNA$s). $C - S_{10} - B$ binds and diffuses along DNA but cannot move beyond stably bound $dCas12a$ (Supplementary Figure S38a). $MSSR$ increased the SNR and resolution of kymographs showing $C - S_{10} - B$ 1D diffusion on a dCas12a-delimited DNA stretch (Supplementary Figure S38b). During self-assembly, large clusters of $C - S_{10} - B$ form and colocalize with $dCas12a$ (Supplementary Figure S38c), as seen in Supplementary Figure S38d, $MSSR$ transformed blurred fluorescent areas into well-defined $C - S_{10} - B$ clusters and made colocalization of $C - S_{10} - B$ and dCas12a more apparent. $MSSR$ can help to recover meaningful information in scenarios of $SPT$ and nanoscopic dynamic colocalization.

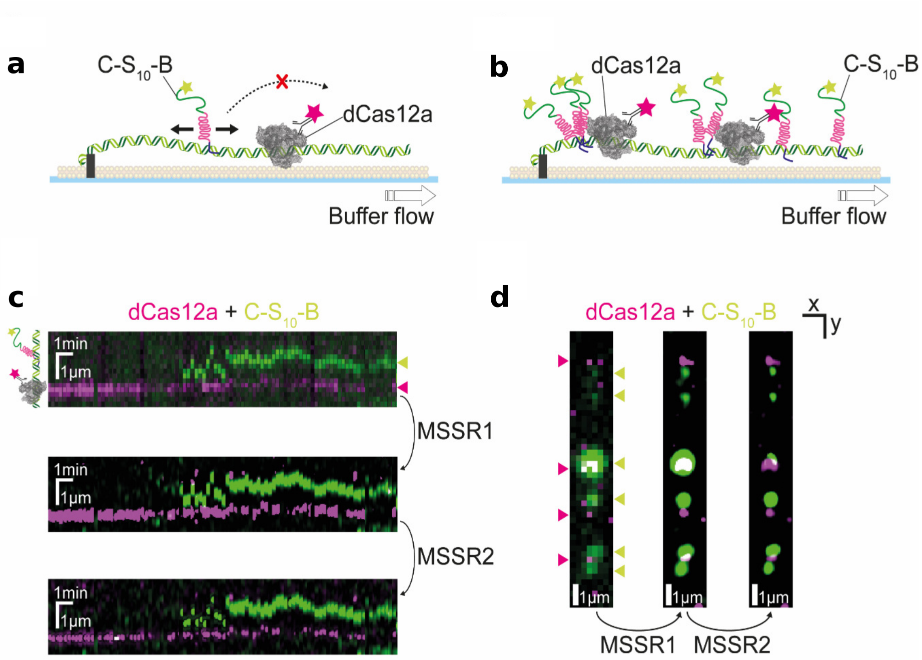

**Figure S38.** ***MSSR* denoises images and improves visualization of intermolecular dynamics on single molecule assays. a)** Linear diffusion of $C - S_{10} - B$ (Alexa488-labeled, green) on the buffer-extended DNA is constrained by stably bound $dCas12a$ (quantum dot-labeled, magenta) on a DNA molecule (unlabeled). **b)** Stalling of $C - S_{10} - B$ diffusion by $dCas12a$ causes co-localization of both molecules. **c)** Processing with one and two $MSSR$ iterations of a kymograph showing $dCas12a$ (immobile, magenta) and $C - S_{10} - B$ (sliding, green) on the DNA (unlabeled). **d)** Processing with one and two $MSSR$ iterations of a DNA molecule (unlabeled) with bound $C - S_{10} - B$ (green) and $dCas12a$ (magenta).

##### 10.6. Volumetric nanoscopy of deep samples

Transgenic *Arabidopsis thaliana* with plasmalemma [58] and nuclear [59] fluorescent markers are routinely used to study cell proliferation during root development in time-lapse experiments. To demonstrate its utility for multidimensional imaging, $MSSR$ was challenged to unveil nanoscopic structures in an *A. thaliana* roots as an example of a thick multicellular specimen, imaged with a confocal laser-scanning microscope ($CLSM$) and selective plane illumination microscopy ($SPIM$). To untangle the denoising properties of $MSSR$ from its spatial resolution enhancement capability, the $PSF$ was under-sampled (see Methods). Note that under such $CLSM$ scenario, mitotic structures cannot be optimally visualized (Supplementary Figure S39 a and b). However, after being post-processed with $MSSR$, the images displayed a great improvement in contrast and clarity which enabled the unveiling of hidden cellular structures (Supplementary Figure S39c).

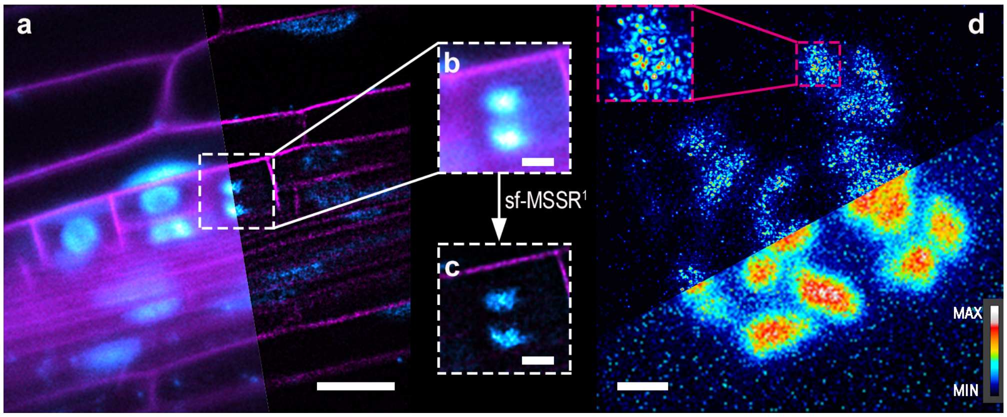

**Figure S39.** $sf-MSSR$ **processing of *A. thaliana* root cells images. a)** Image of early-stage lateral root primordium acquired with regular CLSM. The left side corresponds to the diffraction-limited image, while the right side is the result of $sf-MSSR^1$ image. **b)** Original $ROI$ took from the diffraction-limited image in a), highlighted by the white dashed square. **c)** the same $ROI$ as in b) after being processed by the $sf-MSSR^1$. **d)** Over-sampled and $sf-MSSR^1$ image of lateral root meristematic cell chromosomes seen by $H2B$ labeled with $RFP$. The image comprises the top and down sides, diffraction-limited and $sf-MSSR^1$ images. The inset shows a zoom (3×) $ROI$ highlighted by a pink dashed square. Scales bar in a = 20 $\mu$m, b and c = 5 $\mu$m, d = 1 $\mu$m. $sf-MSSR$ parameters: $AMP$ = 5, $FWHM$ of $PSF$ = 10, $order$ = 1. The final image in a), b) and c) is a Z projection of ten slices with SUM slices mode and the FIJI´s LUT used for red and green channels were "Cyan Hot" and "Magenta", respectively, while the image in "d" was made by a Z projection of 86 slices with MAX intensity mode and the LUT used was "Royal" of FIJI, for more details see methods.

As explained above, *A. thaliana* root cells were imaged with an under-sampled $PSF$, which led us to wonder if $MSSR$ could enable nanoscopic structure visualization with an over-sampled $PSF$. These new data were then processed with $MSSR$, which revealed heterogeneous chromatin nanodomains within the mitotic region (Figure S39d). These structures resemble cluster-like nucleosomes previously reported in mammalian cells [60, 61] and in *A. thaliana* leaf nuclei [62]. These images were acquired as $Z$-stacks, hence $MSSR$ provided nanoscopic resolution in three dimensions (Supplementary Figure S39d, mov. S10). Thus, MSSR allowed visualization of nanoscopic structures in pericycle derivatives cells, located deep inside the root. Interestingly, the resolution in this case was similar to that of an external root tissue as shown in SPIM images of *A. thaliana* epidermal root cells processed with MSSR, specifically of nuclei in root-hair and non-hair cells (Supplementary Figure S40).

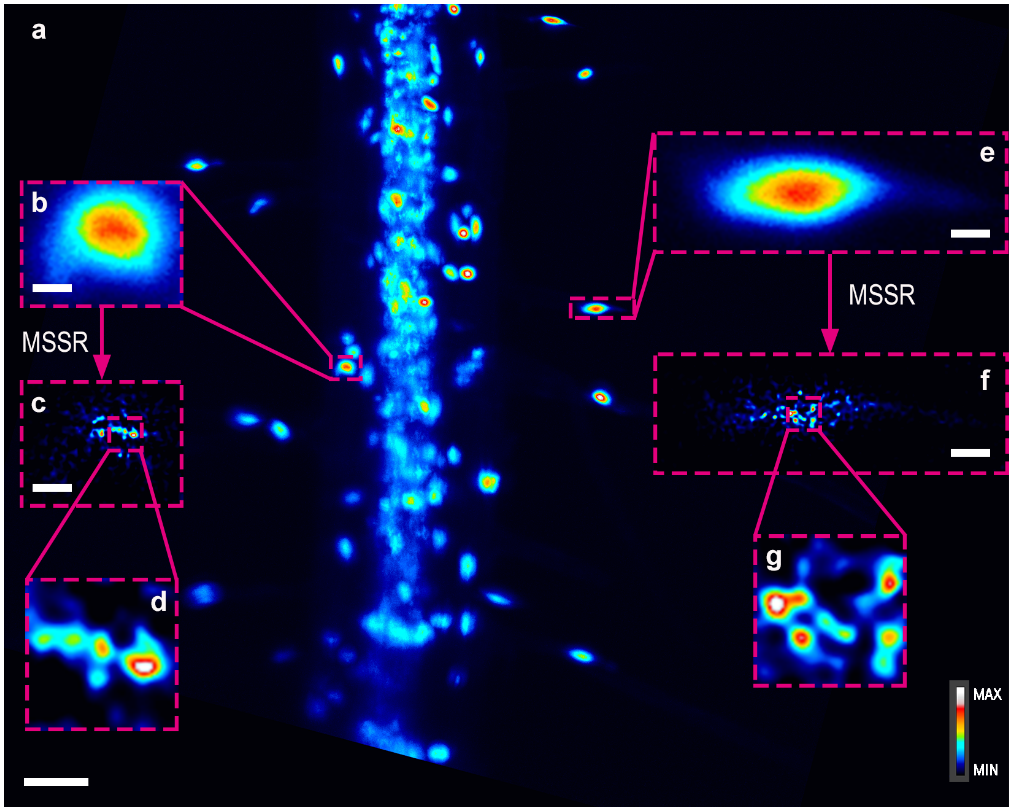

**Figure S40. SPIM images of *A. thaliana* root cells filtered by $sf - MSSR^1$. a)** A differentiated portion of a *p35s:H2B-RFP* transgenic line primary root acquired with SPIM . **b)** A *ROI* of non-root hair nucleus subtracted from the image in the panel a) indicated by the pink dashed square. **c)** The same *ROI* as in b) after being processed by $sf - MSSR^1$. **e)** a *ROI* of root hair nucleus subtracted from the image in the panel a) indicated by the right pink dashed square. **f)** The same *ROI* as in e) after being processed by $sf - MSSR^1$. **d)** and **g)** insets of magnified (3×) *ROI*s in c) and f), respectively, and indicated by the pink dashed squares. $sf - MSSR$ parameters: $AMP = 5$, a $FWHM$ of $PSF = 10$, $order = 1$. Scales bar in a) = 50 $\mu$m, b)-c) and e)-f) = 5 $\mu$m. The final images are Z projections of n slices with MAX intensity mode and the "Royal" LUT of FIJI. See Online Methods for more details.

#### 11. APPLICABILITY OF MSSR TO 3D MICROSCOPY

Since MS theory is not limited by the number of dimensions, the extension of MSSR to three-dimensional (3D) data can be achieved by adding a parameter $h_s$ that also includes the coordinates on the z axis. However, this extension is not trivial. The diffraction in the axial plane is greater than the lateral plane, so the parameter $h_s$ for the $z$-axis needs to be greater compared to the $h_s$ for the $xy$ axes. Hence, the value of $h_s$ for the z-axis will be related to the resolution limit of this axis. On the other hand, an important feature to get a reliable reconstruction when image magnification is performed, is that the number of pixels for each dimension must be increased proportionally.

Supplementary Figure S41 shows a comparison of a 3D reconstruction from a stack of epi-fluorescence images of bovine pulmonary artery endothelial (BPAE) cells. When a 2D image is processed by $sf-MSSR^1$, an improvement in spatial resolution and contrast is observed (Supplementary Figure S41 a, b). On the contrary, when a stack is used for a 3D representation (images taken at $z$-planes), the resolution obtained by $sf-MSSR$ along the z-axis is not as refined as in the $xy$-plane (Supplementary Figure S41 c, d). Nonetheless, a noticeable increase in contrast is attained in the overall data set (Supplementary Figure S41 d, see Supplementary Movies 13 and 14). To further extend the resolution of DL images in 3D, it is necessary to extend the current implementations of the $MSSR$ algorithm to account 3D information for $MSSR$ processing. Therefore, it is necessary to make a detailed study for $MSSR$ in 3D.

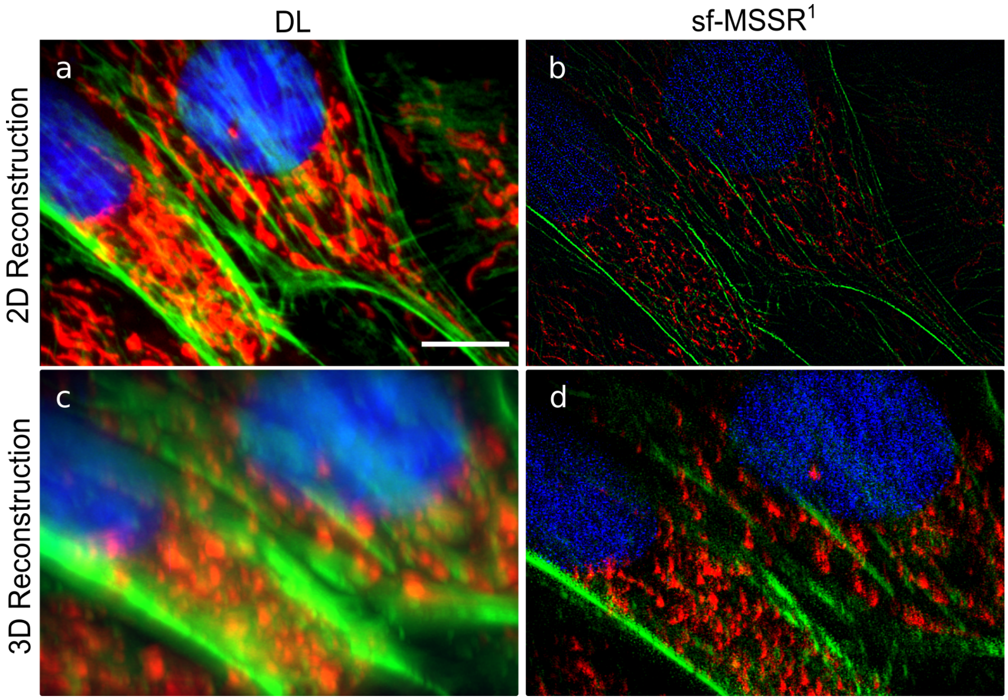

**Figure S41. Comparison DL images of $sf-MSSR$ reconstruction of epifluorescence BPAE cells for 2D and 3D images.** From top to bottom: 2D image for a selective plane and rotated view of the 3D reconstructions. From left to right: DL images of selective planes of the stack and $sf-MSSR$ results. Channels: mitochondria stained with MitoTracker$^{TM}$ Red CMXRos (red), F-actin was stained with Alexa Fluor$^{TM}$ 488 phalloidin (green) and the DNA of the nuclei were stained with DAPI (blue). Scale bar 10 $\mu$m. $sf-MSSR$ parameters: $AMP$ = 10, $FWHM$ of $PSF$ = 2.77, $order$ = 1. For 3D reconstruction the program napari was used with a modification in the voxel.

The $MSSR$ principle can be developed for explicit volumetric imaging by means of using an asymmetric kernel, which can be defined following the 3D lateral-axial aspect ratio of the $PSF$, but also, encompassing possible deformations of axial symmetry of the $PSF$ (due to optical aberrations introduced by the sample or the imaging system). Handling optical aberration in 3D light sheet microscopy is an active field of research (i.e., deconvolution or adaptive optics), where we want to contribute through the further development of $MSSR$ applications in 3D.

#### 12. EXTENDED SUPPLEMENTARY FIGURES

In order to validate the $CRISPR/Cas$ nanoruler system (Supplementary Figure S42a), the protein-DNA complexes were first imaged with atomic force microscopy ($AFM$) in dried conditions to corroborate the distances between DNA fluorophore-binding sites (Supplementary Figure S42b).

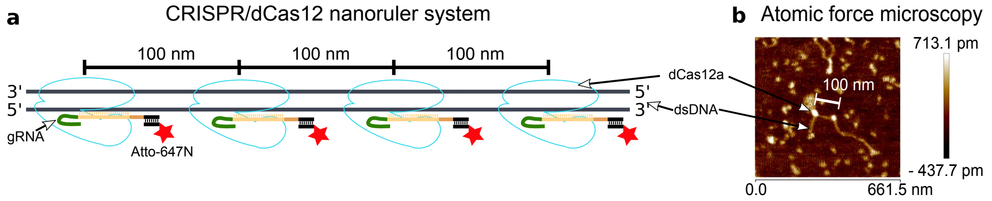

**Figure S42.** $CRISPR/Cas$ nanoruler system, which consists of a 1,537 bp double strand DNA ($dsDNA$) with four binding sites for $dCas12a$ set out apart regularly at lengths of 297 bp (~100 nm) over the $dsDNA$ chain. **a)** Graphical representation of the $CRISPR-PAINT$ nanoruler system. **b)** $CRISPR-PAINT$ nanoruler visualized by atomic force microscopy.

A MSSR temporal analysis of 100 images provided super-resolved images of the nanorulers with a binding-site distance range between 30 and 80 nm. This allowed to validate a lower experimental spatial resolution boundary of 40 nm (Supplementary Figure S43).

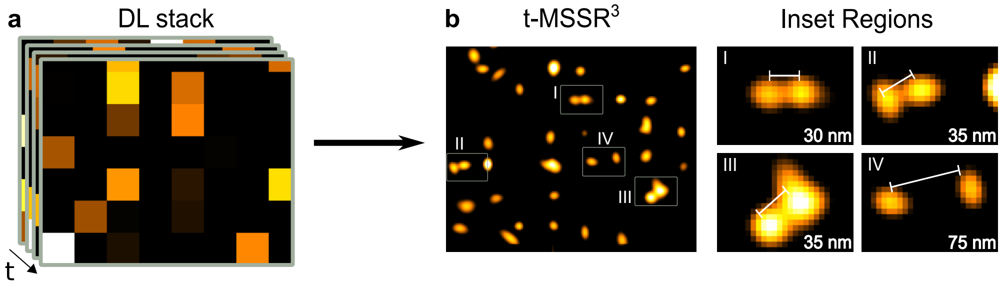

**Figure S43.** $MSSR$ was challenged to resolve a commercially available nanoruler ($GATTA\text{-}PAINT$, $GATTAquant$), in which three chemically bounded fluorophores are spaced by 40 nm each. a) Field of a $GATTA\text{-}PAINT$ nanorulers experiment resolved by $MSSR$. b) Four nanorulers were found in the field, with a distance of 30 nm, 35 nm, 35 nm and 75 nm respectively.

Analyzing the temporal dynamics of fluorescence is key to achieving the highest attainable gain in resolution by fluorescence fluctuation based super resolution microscopy (FF-SRM) approaches (reviewed in [14, 63]). $FF-SRM$ methods analyze higher order temporal statistics in order to discriminate between correlated (fluorescence signal) and uncorrelated (noise) information. $t-MSSR^n$ provides further gain in resolution by means of performing a pixel-wise temporal analysis and thus, $t-MSSR^n$ can be considered a FF-SRM approach.

Supplementary Figure S44 shows the pixel-wise temporal dynamics of two GATTA paint ranorulers imaged for 15 seconds at a 50 ms interval. In the provided examples (nanorulers 1 and 2), the properties of the fluorophores attached to the DNA binding sites can be retrieved by analyzing the signal fluctuations at two pixels (px1 and px2). Note that, by analyzing either px1 or px2, it is difficult to distinguish the temporal dynamics of individual PAINT binding sites. Notably, $sf-MSSR^3$ processing allows to untangle, in space and in time, the fluorescence dynamics at individual PAINT binding sites, where the fluorescence signal peaks due to the transient binding of a fluorescent labeled DNA strand to its corresponding binding site within the nanoruler.

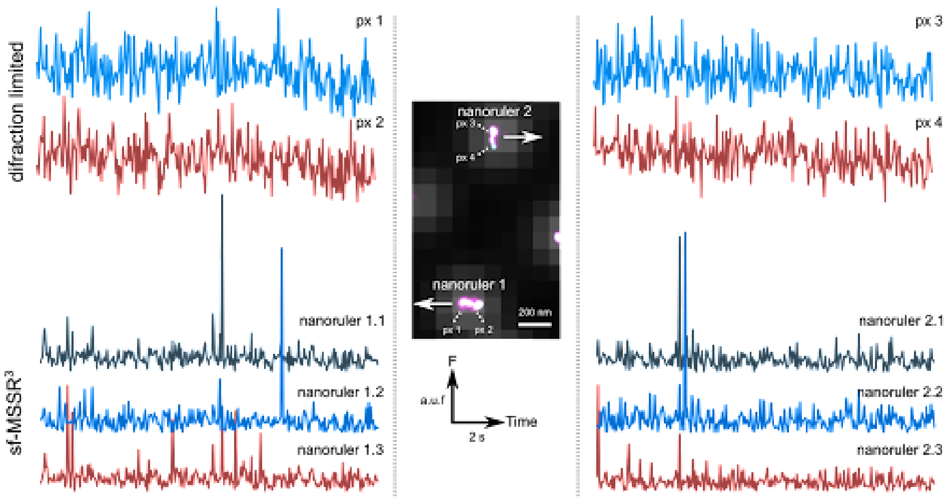

**Figure S44. Temporal fluctuation characteristics of two GattaPaint Nanorulers.** Overlay of DL image and SR reconstruction by $t - MSSR\ Var$ (middle). Two GattaPaint ATTO 550 nanorulers have been highlighted in white color containing three binding sites, each nanoruler is distributed in two pixels corresponding to the equivalent image DL. Upper strokes show the temporal fluctuation of pixels that match with the nanorulers, lower strokes show the temporal fluctuation of each binding site for the two nanorulers. The comparison of the dark times can be seen (unbound DNA-fluorescent strand), when the diffraction-limited data is analyzed against the data resolved by $sf - MSSR^3$.

#### 13. EXTENDED SUPPLEMENTARY MOVIES

- Nanoruler
  - Movie S1 (CRISPR/PAINT nanorulers resolved by MSSR)

    An image sequence of 100 frames of the CRISPR/PAINT nanoruler adquired by TIRF microscopy (diffraction-limited) was analyzed by $sf - MSSR$. Temporal analysis of $t - MSSR$ data by $PTF$ generates the super-resolved micrography. MSSR parameters: $AMP$ = 20, $FWHM$ of $PSF$ for GattaPaint ATTO 550 = 3.31, $order$ = 3.

- Free of noise-dependet artifacts
  - Movie S2 (Laser-written lithography pattern reconstruction at low $SNR$ 5.4).

    This video shows the progressive gain in resolution image when one or several images are used for reconstruction in a low $SNR$ set. The first image corresponds to a single frame analysis with $MSSR$ (no temporal analysis) while subsequent images result from the number of frames indicated on top. In this scenario, a single image is not sufficient for a reliable reconstruction, while at around 20 images used for temporal analysis the reconstruction recovers part of the expected shape of the fluorescent ring. However, there is a point where more images do not contribute to any further resolution enhancement, as seen in Supplementary Figure S28. $sf - MSSR$ and $t - MSSR$ parameters: $AMP$ = 4, $FWHM$ of $PSF$ = 2.37, $order$ = 1, $PTF$ = $TPM$.

  - Movie S3 (Laser-written lithography pattern reconstruction at high $SNR$ 21.3).

    As in Supplementary Movie 2, this video shows the progressive gain in resolution but, in this case, a single image is sufficient for a reliable reconstruction. Further images used in the temporal analysis are not required. This is also quantified in Supplementary Figure S28. $sf - MSSR$ and $t - MSSR$ parameters: $AMP$ = 4, $FWHM$ of $PSF$ = 2.37, $order$ = 1, $PTF$ = $TPM$.

- $MSSR$ extended-resolution and super-resolution videos for single frame and temporal analysis of mouse sperm during acrosome exocytosis.
  - Movie S4 (Super-resolution $sf - MSSR$ head).

    $sf - MSSR$ extended-resolution single frame analysis of sperm acrosome exocytosis. As this process develops, a rise in FM4-64 fluorescence (magenta) and different dynamics in F-actin showed by SiR-actin (green) are seen. $sf - MSSR$ parameters: $AMP$ = 5, $FWHM$ of $PSF$ = 5, $order$ = 0.

  - Movie S5 (Super-resolution $sf - MSSR$ fenestration).

    $sf - MSSR$ extended-resolution single frame analysis of sperm membrane fenestration. Prior to $AE$, the plasma membrane is seen above the cortical F-actin. Once the $AE$ occurs, the plasma membrane is fenestrated and the inner acrosomal membrane is now stained by FM4-64. As a result of this, F-actin localizes between membranes. $sf - MSSR$ parameters: $AMP$ = 5, $FWHM$ of $PSF$ = 5, $order$ = 0.

  - Movie S6 (Super-resolution $t - MSSR$ head of 100 frames).

    $t - MSSR$ super-resolution analysis of 100 temporal 'pseudo' replicates collected at the same scene of sperm acrosome exocytosis. As this process develops, a rise in FM4-64 fluorescence (magenta) and different dynamics in F-actin showed by SiR-actin (green) are seen. $t - MSSR$ parameters: $AMP$ = 5, $FWHM$ of $PSF$ = 5, $order$ = 0, $PTF$ = $Mean$.

  - Movie S7 (Super-resolution $t - MSSR$ fenestration of 100 frames).

    $t - MSSR$ super-resolution analysis of 100 temporal 'pseudo' replicates collected at the same scene of sperm membrane fenestration. Prior to $AE$, the plasma membrane is seen above the cortical F-actin. Once the $AE$ occurs, the plasma membrane is fenestrated and the inner acrosomal membrane is now stained by FM4-64. As a result of this, F-actin localizes between membranes. $t - MSSR$ parameters: $AMP$ = 5, $FWHM$ of $PSF$ = 5, $order$ = 0, $PTF$ = $Mean$.

- Movie S8 (Extended-resolution $sf - MSSR$ fenestration of 100 frames at t0).

  $sf - MSSR$ extended-resolution frame analysis of sperm membrane fenestration prior to $AE$. By using 'pseudo' replicates collected at the same scene before the $AE$ occurs, different dynamics of plasma membrane and cortical F-actin are seen at a millisecond scale. $sf - MSSR$ parameters: $AMP = 5$, $FWHM$ of $PSF = 5$, $order = 0$.

- Movie S9 (Extended-resolution $sf - MSSR$ fenestration of 100 frames at t13).

  $sf - MSSR$ extended-resolution single frame analysis of sperm membrane fenestration after $AE$. By using 'pseudo' replicates collected at the same scene after the $AE$ occurs, different dynamics of plasma membrane and cortical F-actin are seen at a millisecond scale. $sf - MSSR$ parameters: $AMP = 5$, $FWHM$ of $PSF = 5$, $order = 0$.

- $sf - MSSR$ extended-resolution movies of microtubule dynamics in LLC-PK1 cells.

  - Movie S10 ($sf - MSSR$ video of live LLC-PK1 cells expressing mEmerald-EB3).

    Extended-resolution video of microtubule dynamics in live LLC-PK1 cells stably expressing mEmerald-EB3. Right side are two $2\times$ enlargements of the areas depicted by the green squares. Time scale is 2 seconds per frame, playing at $30\times$ speed. $sf - MSSR$ parameters: $AMP = 3$, $FWHM$ of $PSF = 30$, $order = 0$.

  - Movie S11 ($sf - MSSR$ video of an apoptotic LLC-PK1 cell expressing mEmerald-EB3).

    Extended-resolution video of microtubule dynamics in an apoptotic LLC-PK1 cell stably expressing mEmerald-EB3. Membrane blebbing can be seen in the contour of the cell. Time scale is 2 seconds per frame, playing at $30\times$ speed. $sf - MSSR$ parameters: $AMP = 3$, $FWHM$ of $PSF = 30$, $order = 0$.

- Extended-resolution 3D reconstruction of Z-stack slices of *A. thaliana* root nucleosome.

  - Movie S12 (Extended-resolution 3D reconstruction of Z-stack slices of *A. thaliana* root nucleosome).

    3D representation of *A. thaliana* root nucleosomes labeled with RFP (p35s:H2B-RFP). The images were acquired with regular Olympus CLSM (see methods) and filtered with $sf - MSSR$. The 3D reconstruction was made by the 3D Viewer plugin of $FIJI$, with 86 slices Z stack with a step size of 100 nm. For more details see methods.

- Comparison of DL reconstruction vs $sf - MSSR^1$ reconstruction of epifluorescence BPAE cells for 2D and 3D images

  - Movie S13 (DL reconstruction of epifluorescence BPAE cells for 2D and 3D images).

    Selective plane and rotated view of epifluorescence BPAE cells. Channels: mitochondria stained with MitoTracker$^{TM}$ Red CMXRos (red), F-actin was stained with Alexa Fluor$^{TM}$ 488 phalloidin (green), and the DNA of the nuclei was stained with DAPI (blue). The video was generated with Napari, with the plug-in animation: wizard.

  - Movie S14 ($sf - MSSR^1$ reconstruction of epifluorescence BPAE cells for 2D and 3D images).

    Selective plane and rotated view of $sf - MSSR^1$ results of epifluorescence BPAE cells. Channels: mitochondria stained with MitoTracker$^{TM}$ Red CMXRos (red), F-actin was stained with Alexa Fluor$^{TM}$ 488 phalloidin (green), and the DNA of the nuclei was stained with DAPI (blue). $sf - MSSR$ parameters: $AMP = 10$, $FWHM$ of $PSF = 2.77$, $order = 1$. The video was generated with Napari, with the plug-in animation: wizard.

#### REFERENCES

1. K. Ahi, "Mathematical modeling of thz point spread function and simulation of thz imaging systems," IEEE Transactions on Terahertz Sci. Technol. **7**, 747–754 (2017).
2. A. Small, I. Ilev, V. Chernomordik, and A. Gandjbakhche, "Enhancing diffraction-limited images using properties of the point spread function," Opt. Express **14**, 3193–3203 (2006).
3. B. Zhang, J. Zerubia, and J.-C. Olivo-Marin, "Gaussian approximations of fluorescence microscope point-spread function models," Appl. Opt. **46**, 1819–1829 (2007).
4. K. Fukunaga and L. Hostetler, "The estimation of the gradient of a density function, with applications in pattern recognition," IEEE Transactions on Inf. Theory **21**, 32–40 (1975).
5. Yizong Cheng, "Mean shift, mode seeking, and clustering," IEEE Transactions on Pattern Analysis Mach. Intell. **17**, 790–799 (1995).
6. D. Comaniciu and P. Meer, "Mean shift analysis and applications," in *Proceedings of the Seventh IEEE International Conference on Computer Vision,* vol. 2 (1999), pp. 1197–1203 vol.2.
7. Weglarczyk, "Kernel density estimation and its application," ITM Web Conf. **23**, 00037 (2018).
8. D. Comaniciu, V. Ramesh, and P. Meer, "Real-time tracking of non-rigid objects using mean shift," in *Proceedings IEEE Conference on Computer Vision and Pattern Recognition. CVPR 2000 (Cat. No.PR00662),* vol. 2 (2000), pp. 142–149 vol.2.
9. D. Comaniciu and P. Meer, "Mean shift: a robust approach toward feature space analysis," IEEE Transactions on Pattern Analysis Mach. Intell. **24**, 603–619 (2002).
10. I. M. Khater, F. Meng, T. H. Wong, I. R. Nabi, and G. Hamarneh, "Super resolution network analysis defines the molecular architecture of caveolae and caveolin-1 scaffolds," Sci. Reports **8**, 9009 (2018).
11. J. Boulanger, C. Kervrann, P. Bouthemy, P. Elbau, J.-B. Sibarita, and J. Salamero, "Patch-Based Nonlocal Functional for Denoising Fluorescence Microscopy Image Sequences," IEEE Transactions on Medical Imaging **29**, 442–453 (2010).
12. F. J. Fazekas, T. R. Shaw, S. Kim, R. A. Bogucki, and S. L. Veatch, "A Mean Shift Algorithm for Drift Correction in Localization Microscopy," bioRxiv (2021). Publisher: Cold Spring Harbor Laboratory _eprint: https://www.biorxiv.org/content/early/2021/05/07/2021.05.07.443176.full.pdf.
13. M. Pawlowska, R. Tenne, B. Ghosh, A. Makowski, and R. Lapkiewicz, "Embracing the uncertainty: the evolution of SOFI into a diverse family of fluctuation-based super-resolution microscopy methods," J. Physics: Photonics **4**, 012002 (2021).
14. T. Dertinger, R. Colyer, G. Iyer, S. Weiss, and J. Enderlein, "Fast, background-free, 3d super-resolution optical fluctuation imaging (sofi)," Proc. Natl. Acad. Sci. **106**, 22287–22292 (2009).
15. R. Keys, "Cubic convolution interpolation for digital image processing," IEEE Transactions on Acoust. Speech, Signal Process. **29**, 1153–1160 (1981).
16. S. Reichenbach and F. Geng, "Two-dimensional cubic convolution," IEEE Transactions on Image Process. **12**, 857–865 (2003).
17. M. Hirsch, R. J. Wareham, M. L. Martin-Fernandez, M. P. Hobson, and D. J. Rolfe, "A stochastic model for electron multiplication charge-coupled devices – from theory to practice," PLOS ONE **8**, 1–13 (2013).
18. A. D. Corbett, M. Shaw, A. Yacoot, A. Jefferson, L. Schermelleh, T. Wilson, M. Booth, and P. S. Salter, "Microscope calibration using laser written fluorescence," Opt. Express **26**, 21887–21899 (2018).
19. S. Culley, D. Albrecht, C. Jacobs, P. M. Pereira, C. Leterrier, J. Mercer, and R. Henriques, "Quantitative mapping and minimization of super-resolution optical imaging artifacts," Nat. methods **15**, 263–266 (2018). Edition: 2018/02/19.
20. F. Huang, T. M. P. Hartwich, F. E. Rivera-Molina, Y. Lin, W. C. Duim, J. J. Long, P. D. Uchil, J. R. Myers, M. A. Baird, W. Mothes, M. W. Davidson, D. Toomre, and J. Bewersdorf, "Video-rate nanoscopy using sCMOS camera–specific single-molecule localization algorithms," Nat. Methods **10**, 653–658 (2013).
21. N. Gustafsson, S. Culley, G. Ashdown, D. M. Owen, P. M. Pereira, and R. Henriques, "Fast live-cell conventional fluorophore nanoscopy with ImageJ through super-resolution radial fluctuations," Nat. Commun. **7**, 12471 (2016).
22. I. Yahiatene, S. Hennig, M. Müller, and T. Huser, "Entropy-Based Super-Resolution Imaging (ESI): From Disorder to Fine Detail," ACS Photonics **2**, 1049–1056 (2015). Publisher: American Chemical Society.
23. K. Agarwal and R. Macháň, "Multiple signal classification algorithm for super-resolution fluorescence microscopy," Nat. Commun. **7**, 13752 (2016).

24. S. Acuna, F. Strohl, I. S. Opstad, B. S. Ahluwalia, and K. Agarwal, “Musij: an imagej plugin for video nanoscopy,” Biomed. Opt. Express **11**, 2548–2559 (2020).
25. M. G. Gustafsson, “Surpassing the lateral resolution limit by a factor of two using structured illumination microscopy.” J. microscopy **198**, 82–87 (2000). Place: England.
26. B. Mandracchia, X. Hua, C. Guo, J. Son, T. Urner, and S. Jia, “Fast and accurate sCMOS noise correction for fluorescence microscopy,” Nat. Commun. **11**, 94 (2020).
27. B. N. Fields, D. M. Knipe, and P. M. Howley, *Fields virology* (Wolters Kluwer Health/Lippincott Williams & Wilkins, Philadelphia, 2013).
28. Y. Garcés Suárez, J. L. Martínez, D. Torres Hernández, H. O. Hernández, A. Pérez-Delgado, M. Méndez, C. D. Wood, J. M. Rendon-Mancha, D. Silva-Ayala, S. López, A. Guerrero, and C. F. Arias, “Nanoscale organization of rotavirus replication machineries,” eLife **8**, e42906 (2019).
29. H. O. Hernández, P. Hidalgo, C. D. Wood, R. González, and A. Guerrero, “Parallelizing the Bayesian Analysis of Blinking and Bleaching for Super-Resolution Microscopy,” in *High Performance Computer Applications*, I. Gitler and J. Klapp, eds. (Springer International Publishing, Cham, 2016), pp. 356–366.
30. P. A. Balestrini, M. Jabloñski, L. J. Schiavi-Ehrenhaus, C. I. Marín-Briggiler, C. Sánchez-Cárdenas, A. Darszon, D. Krapf, and M. G. Buffone, “Seeing is believing: Current methods to observe sperm acrosomal exocytosis in real time,” Mol. Reproduction Dev. **87**, 1188–1198 (2020).
31. A. Romarowski, A. G. Velasco Félix, P. Torres Rodríguez, M. G. Gervasi, X. Xu, G. M. Luque, G. Contreras-Jiménez, C. Sánchez-Cárdenas, H. V. Ramírez-Gómez, D. Krapf, P. E. Visconti, D. Krapf, A. Guerrero, A. Darszon, and M. G. Buffone, “Super-resolution imaging of live sperm reveals dynamic changes of the actin cytoskeleton during acrosomal exocytosis,” J. Cell Sci. **131** (2018). Jcs218958.
32. C. Sánchez-Cárdenas, M. R. Servín-Vences, O. José, C. L. Treviño, A. Hernández-Cruz, and A. Darszon, “Acrosome Reaction and Ca2+ Imaging in Single Human Spermatozoa: New Regulatory Roles of [Ca2+]i1,” Biol. Reproduction **91** (2014). 67, 1-13.
33. D. Krapf and R. Metzler, “Strange interfacial molecular dynamics,” Phys. Today **72**, 48–54 (2019).
34. S. Sadegh, J. L. Higgins, P. C. Mannion, M. M. Tamkun, and D. Krapf, “Plasma membrane is compartmentalized by a self-similar cortical actin meshwork,” Phys. Rev. X **7**, 011031 (2017).
35. C. Manzo and M. F. Garcia-Parajo, “A review of progress in single particle tracking: from methods to biophysical insights,” Reports on Prog. Phys. **78**, 124601 (2015).
36. S. Jin, P. M. Haggie, and A. S. Verkman, “Single particle tracking of membrane protein diffusion in a potential: Simulation, detection, and application to confined diffusion of cftr cl-channels,” Biophys. J. **93**, 1079–1088 (2007).
37. C.-L. Hsieh, S. Spindler, J. Ehrig, and V. Sandoghdar, “Tracking single particles on supported lipid membranes: Multimobility diffusion and nanoscopic confinement,” The J. Phys. Chem. B **118**, 1545–1554 (2014). PMID: 24433014.
38. Z. Ye, X. Wang, and L. Xiao, “Single-particle tracking with scattering-based optical microscopy,” Anal. Chem. **91**, 15327–15334 (2019). PMID: 31751513.
39. F. Pampaloni, U. Berge, A. Marmaras, P. Horvath, R. Kroschewski, and E. H. K. Stelzer, “Tissue-culture light sheet fluorescence microscopy (TC-LSFM) allows long-term imaging of three-dimensional cell cultures under controlled conditions,” Integr. Biol. **6**, 988–998 (2014).
40. N. Yamashita, M. Morita, W. R. Legant, B.-C. Chen, E. Betzig, H. Yokota, and Y. Mimori-Kiyosue, “Three-dimensional tracking of plus-tips by lattice light-sheet microscopy permits the quantification of microtubule growth trajectories within the mitotic apparatus,” J. Biomed. Opt. **20**, 1 – 18 (2015).
41. M. Held, I. Santeramo, B. Wilm, P. Murray, and R. Lévy, “Ex vivo live cell tracking in kidney organoids using light sheet fluorescence microscopy,” PLOS ONE **13**, 1–20 (2018). Publisher: Public Library of Science.
42. N. Chenouard, I. Smal, F. de Chaumont, M. Maška, I. F. Sbalzarini, Y. Gong, J. Cardinale, C. Carthel, S. Coraluppi, M. Winter, A. R. Cohen, W. J. Godinez, K. Rohr, Y. Kalaidzidis, L. Liang, J. Duncan, H. Shen, Y. Xu, K. E. G. Magnusson, J. Jaldén, H. M. Blau, P. Paul-Gilloteaux, P. Roudot, C. Kervrann, F. Waharte, J.-Y. Tinevez, S. L. Shorte, J. Willemse, K. Celler, G. P. van Wezel, H.-W. Dan, Y.-S. Tsai, C. O. de Solórzano, J.-C. Olivo-Marin, and E. Meijering, “Objective comparison of particle tracking methods,” Nat. Methods **11**, 281–289 (2014).

43. J.-Y. Tinevez, N. Perry, J. Schindelin, G. M. Hoopes, G. D. Reynolds, E. Laplantine, S. Y. Bednarek, S. L. Shorte, and K. W. Eliceiri, "Trackmate: An open and extensible platform for single-particle tracking," Methods **115**, 80–90 (2017). Image Processing for Biologists.
44. K. Jaqaman, D. Loerke, M. Mettlen, H. Kuwata, S. Grinstein, S. L. Schmid, and G. Danuser, "Robust single-particle tracking in live-cell time-lapse sequences," Nat. Methods **5**, 695–702 (2008).
45. R. E. Kalman, "A new approach to linear filtering and prediction problems," J. Basic Eng. **82**, 35–45 (1960).
46. G. Welch and G. Bishop, "An introduction to the kalman filter," Tech. Rep. 95-041, University of North Carolina at Chapel Hill, Chapel Hill, NC, USA (1995).
47. J. C. Crocker and D. G. Grier, "Methods of Digital Video Microscopy for Colloidal Studies," J. Colloid Interface Sci. **179**, 298–310 (1996).
48. J. Mazzaferri, J. Roy, S. Lefrancois, and S. Costantino, "Adaptive settings for the nearest-neighbor particle tracking algorithm," Bioinformatics **31**, 1279–1285 (2014).
49. M. Patel, S. E. Leggett, A. K. Landauer, I. Y. Wong, and C. Franck, "Rapid, topology-based particle tracking for high-resolution measurements of large complex 3D motion fields," Sci. Reports **8**, 5581 (2018).
50. http://icy.bioimageanalysis.com.
51. [http://bioimageanalysis.org/track/.
52. X. Wu and J. A. Hammer, "Zeiss airyscan: Optimizing usage for fast, gentle, super-resolution imaging," in *Confocal Microscopy: Methods and Protocols*, J. Brzostowski and H. Sohn, eds. (Springer US, New York, NY, 2021), pp. 111–130.
53. R. Heald and E. Nogales, "Microtubule dynamics," J. Cell Sci. **115**, 3–4 (2002).
54. A. Akhmanova and C. C. Hoogenraad, "Microtubule plus-end-tracking proteins: mechanisms and functions," Curr. Opin. Cell Biol. **17**, 47–54 (2005).
55. D. van de Willige, C. C. Hoogenraad, and A. Akhmanova, "Microtubule plus-end tracking proteins in neuronal development," Cell. Mol. Life Sci. **73**, 2053–2077 (2016).
56. W. Li, T. Miki, T. Watanabe, M. Kakeno, I. Sugiyama, K. Kaibuchi, and G. Goshima, "Eb1 promotes microtubule dynamics by recruiting sentin in drosophila cells," J. Cell Biol. **193**, 973–983 (2011).
57. G. M. Cooper, *The cell : a molecular approach* (ASM Press; Sinauer Associates, Washington, D.C.; Sunderland, Mass., 2000).
58. N. Geldner, V. Dénervaud-Tendon, D. L. Hyman, U. Mayer, Y.-D. Stierhof, and J. Chory, "Rapid, combinatorial analysis of membrane compartments in intact plants with a multicolor marker set," The Plant journal : for cell molecular biology **59**, 169–178 (2009). Edition: 2009/02/26.
59. F. Federici, L. Dupuy, L. Laplaze, M. Heisler, and J. Haseloff, "Integrated genetic and computation methods for in planta cytometry," Nat. Methods **9**, 483–485 (2012).
60. J. Otterstrom, A. Castells-Garcia, C. Vicario, P. A. Gomez-Garcia, M. P. Cosma, and M. Lakadamyali, "Super-resolution microscopy reveals how histone tail acetylation affects DNA compaction within nucleosomes in vivo," Nucleic Acids Res. **47**, 8470–8484 (2019).
61. M. A. Ricci, C. Manzo, M. F. García-Parajo, M. Lakadamyali, and M. P. Cosma, "Chromatin fibers are formed by heterogeneous groups of nucleosomes in vivo." Cell **160**, 1145–1158 (2015). Place: United States.
62. K. Rutowicz, M. Lirski, B. Mermaz, G. Teano, J. Schubert, I. Mestiri, M. A. Kroteń, T. N. Fabrice, S. Fritz, S. Grob, C. Ringli, L. Cherkezyan, F. Barneche, A. Jerzmanowski, and C. Baroux, "Linker histones are fine-scale chromatin architects modulating developmental decisions in arabidopsis," Genome Biol. **20**, 157 (2019).
63. I. S. Opstad, D. H. Hansen, S. A. na, F. Ströhl, A. Priyadarshi, J.-C. Tinguely, F. T. Dullo, R. A. Dalmo, T. Seternes, B. S. Ahluwalia, and K. Agarwal, "Fluorescence fluctuation-based super-resolution microscopy using multimodal waveguided illumination," Opt. Express **29**, 23368–23380 (2021).
