## Supplementary material for "Nanoscopic resolution within a single imaging frame": MSSR User Manual

### MSSR: Mean-Shift Super Resolution ImageJ plugin

#### Operation Manual

Release 2.0.0, 27/05/2022

#### Disclaimer

*The MSSR plugin is open-source and is provided "as is" and without any warranty. It is not subject to copyright protection and is in the public domain. The authors assume no responsibility whatsoever by its use by other parties, and make no guarantees, expressed or implied, about its quality, reliability, or any other characteristic. Your access to and use of the MSSR plugin is at your own risk. The authors disclaim all liability for and are not responsible for any harm to your computer system, loss or corruption of data, or other harm that results from your access to or use of the plugin. The MSSR plugin is subject to FIJI/ImageJ use policies and disclaimer. For more details, see https://imagej.nih.gov/ij/disclaimer.html.*

### 1. Introduction

This document is designed to help you learn how to use the MSSR plugin through the FIJI/ImageJ GUI.

The fundamentals of the Mean-Shift Super Resolution (MSSR) algorithm are described in (Torres-García E. et al. *bioRxiv*, 2021). MSSR was developed to extract nanoscopic information from digital images limited by diffraction through either a single (*sf* – MSSR) or multi-frame analytical approach (*t* – MSSR). MSSR is an iterative algorithm that enhances image resolution and contrast per *n*-iteration step. We refer to this as *n*-order MSSR ($MSSR^n$, $n > 0$). $MSSR^n$ is compatible with both *sf* – $MSSR^n$ and *t* – $MSSR^n$ approaches.

There is a broad range of fluorescence microscopy and bioimaging applications in which MSSR can be used. These applications include but are not limited to:

• Immunofluorescence imaging of fixed cells by regular epifluorescence microscopy.
• Live-cell imaging using organic dyes, fluorescent proteins, quantum dots.
• Total Internal Reflection Fluorescence microscopy.
• Single-particle tracking.
• Single-molecule localization microscopy.
• Colocalization microscopy.
• Volumetric imaging at either single cell or tissues.
• Confocal laser-scanning microscopy.
• Selective Plane Illumination Microscopy.

In addition, MSSR is compatible with other super-resolution microscopy approaches, such as Structured Illumination Microscopy (SIM), Image Scanning Microscopy (ISM), Stimulated Emission Depletion (STED) microscopy, Super-Resolution Radial Fluctuations Microscopy (SRRF), Entropy-Based Super-Resolution Imaging (ESI), Multiple Signal Classification Algorithm for super-resolution fluorescence microscopy (MUSICAL) and Super Resolution Optical Fluctuation Imaging (SOFI). Usage of MSSR to process digital images not collected in the realm of fluorescence microscopy (e.g., phase-contrast images or electron microscopy images) must be conducted cautiously.

The MSSR plugin is intended to operate over either single images or image stacks of three dimensions, where the third dimension can be space or time. The MSSR algorithm operates over each image of a stack separately (*sf* – MSSR), providing a resolution increase down to $1.6 \sigma$ (when using higher orders of MSSR, i.e., when computing *sf* – $MSSR^n$).

*The Abbe's diffraction limit can be found at $2.5 \sigma$, where sigma indicates the distance between two emitters expressed as σ-times their individual standard deviation.*

A further resolution increase can be achieved through gathering information from the fluorescence dynamics by means of using a Pixel-wise Temporal Function (PTF) which, depending on the photophysical properties of the sample, can deliver nanoscopic detail down to $0.5 \sigma$. The use of a PTF with MSSR encompasses a temporal analysis (*t* – $MSSR^n$), where each individual image included on the temporal analysis is assumed to be collected from the same static or pseudo-static scene.

Hyperstacks (i.e., image stacks with more than three dimensions) must be first split into separate three-dimensional stacks before MSSR processing; doing otherwise might cause data rearrangement, which could introduce the risk of data misinterpretation.

#### 2. Downloading MSSR from GitHub

Navigate to https://github.com/MSSRSupport/MSSR to download the MSSR plugin file:

1. Click “MSSR\_1.0.0.jar” (Figure 1a).

*This file name is subject to change as newer versions of this plugin are released in the future.*

2. Click the download button (Figure 1b) and save the file in your system (Figure 1c).

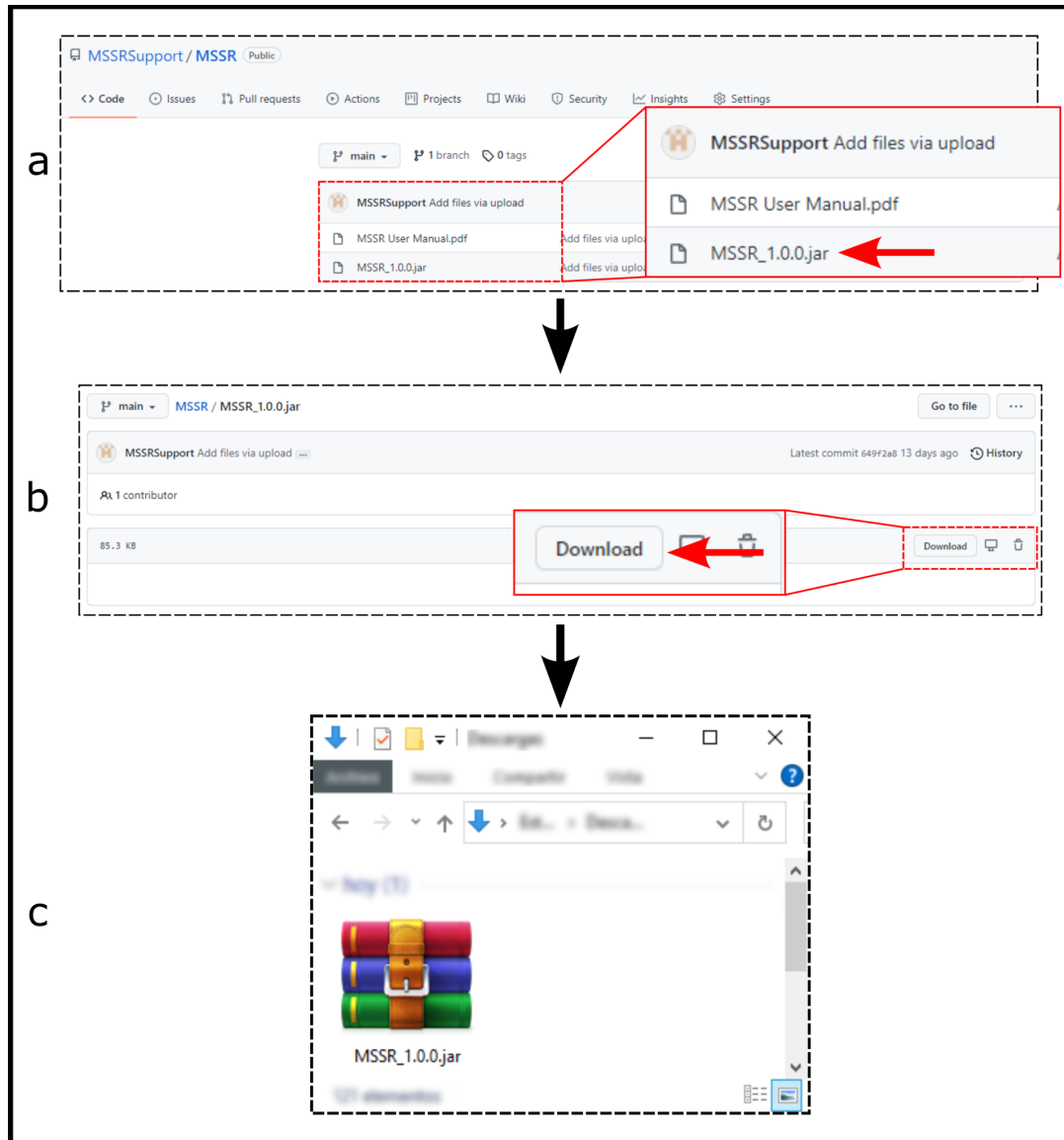

**Figure 1. GitHub download process.** a) Navigate to https://github.com/MSSRSupport/MSSR and click in the “MSSR\_1.0.0.jar” file (red arrow). b) Browse for the “download” option (red arrow) and click it. c) Save the file to a desired directory in your system.

##### 3. Installation

Prior to MSSR installation, the **latest version of FIJI** must be running on your computer (https://fiji.sc/). Additionally, the **CLIJ**, **CLIJ2** and **CLIJx** packages (https://clij.github.io/clij2-docs/installationInFiji) must be installed (Figure 2c).

Plugin installation of MSSR can occur in two alternative ways:

1. Through the installation option in FIJI (Plugins -> Install -> *MSSR\_1.0.0.jar*) (Figure 2c).
2. By directly placing the *MSSR\_1.0.0.jar* file in the specific FIJI *plugins* folder in your system (Figure 2b).

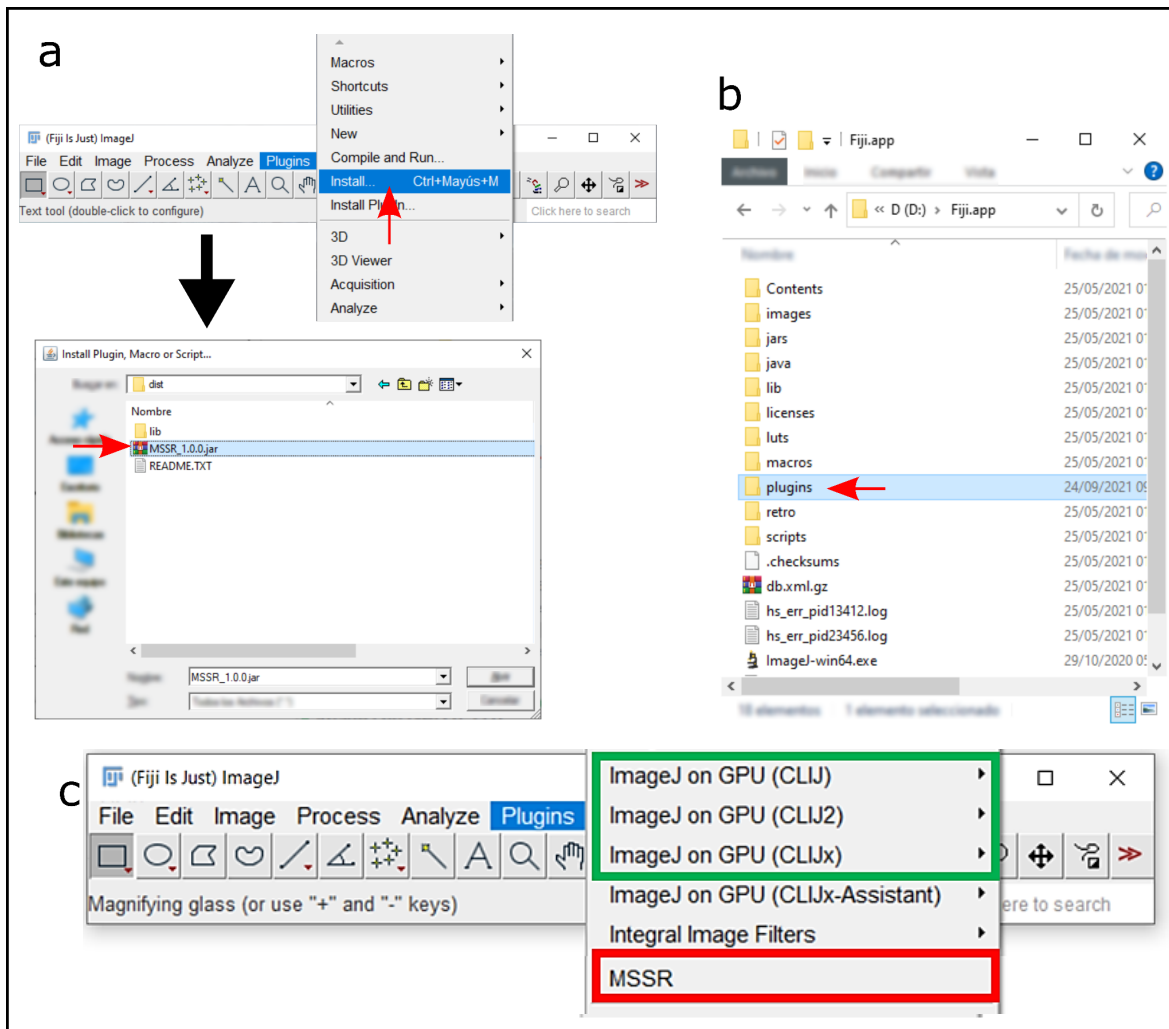

**Figure 2. MSSR plugin installation procedure.** Once downloaded from GitHub, install MSSR by either: **a)** navigating to the plugin installation option in FIJI and selecting the *MSSR\_1.0.0.jar* file. FIJI software must be restarted in order to complete the installation process; **b)** while FIJI is closed, navigate to the "Fiji.app" folder in your system (root directory of the FIJI software), access the "plugins" folder and paste a copy of the MSSR jar file there. Next, open FIJI to complete the installation. **c)** Once installed, the MSSR plugin will be accessible through the "plugins" menu in FIJI (red rectangle) within the currently installed plugins. The CLIJ packages must be installed in order to use the GPU processing feature of MSSR (green rectangle).

#### 4. MSSR plugin

Two options are available within the MSSR plugin (Figure 3a):

• MSSR Analysis - *Encompasses the major processing steps for either $sf - MSSR^n$ or $t - MSSR^n$.*
• Temporal Analysis - *Allows the user to perform an additional temporal analysis by selecting a desired PTF for $t - MSSR^n$ processing.*

In what follows the main parameters available for MSSR analysis computation will be explained, hence, description will be centered on the use of the **MSSR Analysis** tab of the MSSR plugin. A detailed explanation of the temporal analysis is provided in section 4.4 of this document.

Three parameters are needed for MSSR analysis (Figure 3b.1) (refer to section 4 of this document for a detailed description of each of them, as well as of each additional feature, mentioned below):

○ *AMP* – An upscaling factor for the resulting MSSR image size.
○ *FWHM* – The number of pixels that cover the Full Width at Half Maximum of the Point Spread Function (PSF) of the imaging lens.
○ *Order* – The number of MSSR iterations for the image resolution enhancement.

The plugin offers the option of computing $sf - MSSR^n$, or both $sf - MSSR^n$ and $t - MSSR^n$. The temporal analysis is enabled when selecting the option “MSSR Temporal analysis” (Figure 3b.3) where the user can choose one of five available PTFs: Mean, Variance (Var), Temporal Product Mean (TPM), Coefficient Variation, Auto-cumulant Function of order 2-4 (SOFI 2-4).

Additional features:

○ *Computation of FWHM* – provides an estimation of the imaging system’s Rayleigh criterion based on known optical parameters (Figure 3c).
○ *Interpolation type* – allows you to select between two types of interpolation to magnify the image. The default option for this parameter is ‘Bicubic’.
○ *Minimize Meshing* – Enable the mesh minimization algorithm which minimizes a ‘mesh’ pattern that commonly appears during the analysis as result of using a bicubic interpolation algorithm for digital upscaling (Figure 3b.2). The default option for this parameter is active.
○ *GPU Computing* – Enables GPU usage for computing for MSSR processing (Figure 3b.2).
○ *Intensity Normalization* – Allow a pixel-wise multiplication with the MSSR image (scaled from 0 to 1 in pixel values) with the magnified original image.
○ *Selecting Image*– Select a desired image or image stack for MSSR processing from the images which are already loaded in FIJI/ImageJ (Figure 3b.4).
○ *Batch Analysis* – Allow to automatically analyze all the images within a selected folder in the user’s computer (Figure 3b.4).

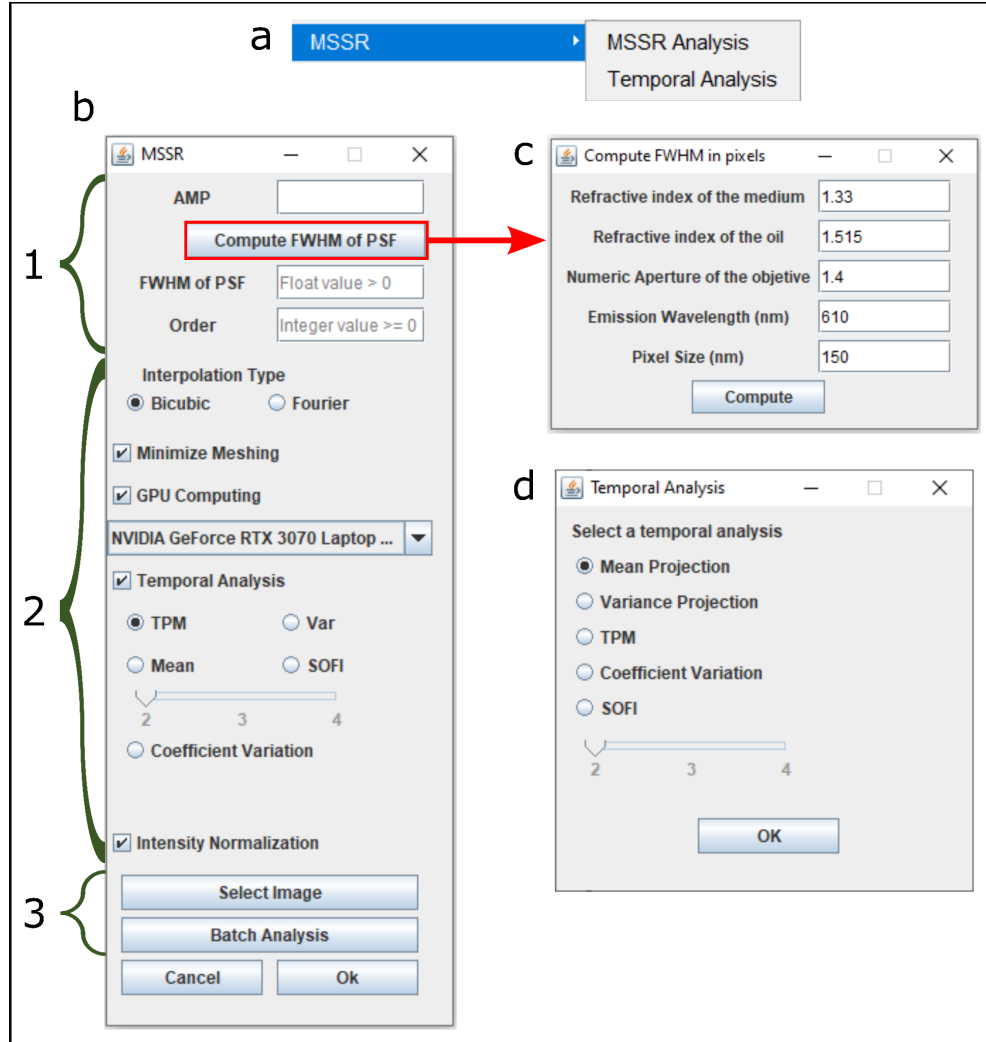

**Figure 3. Available tools and parameters within the MSSR plugin. a)** Choosing between the two available MSSR GUIs is needed instantly after opening the plugin. **b)** The main GUI for analysis where, first (1) the AMP, FWHM and Order parameters need to be specified. (2) The “Interpolation Type” option let you choose between two types of image interpolation (Bicubic or Fourier); The “Minimize Meshing” feature will compensate for the mesh-like artifact that emerges from the analysis; GPU Computing will improve computing speed; A Temporal Analysis can be optionally included if an open image stack is available. Once selected, choose between the four available PTFs. The “Intensity Normalization” feature will pixel-wise multiply the MSSR image (scaled from 0 to 1 in pixel values) with the magnified original image. (3) Next, input image selection, either one which is already open in FIJI or a set of images within a specific path in your computer, is done. Once selected, click the “Ok” button to start the MSSR processing. **c)** Automatic calculation of the FWHM of your specific optical system can be carried out by selecting the “Compute FWHM of PSF” option. **d)** Selecting the “Temporal Analysis GUI in A) will open a new window, where a PTF can be selected and applied to a previously generated sf-MSSR<sup>n</sup> stack.

#### 5. MSSR parameters and specification criteria

##### 5.1. AMPLIFICATION (AMP)

A magnification value that defines the digital zoom (upscaling) to be applied. This parameter takes integer numbers equal to or greater than 1.

Selecting an AMP = 1 will allow MSSR processing with further digital magnification. This option is recommended to be in theoretical or experimental scenarios where the PSF of the optical system is oversampled (i.e., at a pixel size down to 50 nm). The selection of AMP > 1 is recommended when the PSF of the optical system is sampled near or below the recommended Nyquist spatial frequency.

As an example, if the input image (diffraction-limited) has $N \times M$ dimensions with a pixel size of 100 nm, and an amplification value of 5 is used, the resulting extended-resolved image will then be of $(5*N) \times (5*M)$ dimensions with a pixel size of 20 nm ($5 \text{ px} = 100 \text{ nm}$) (Figure 4).

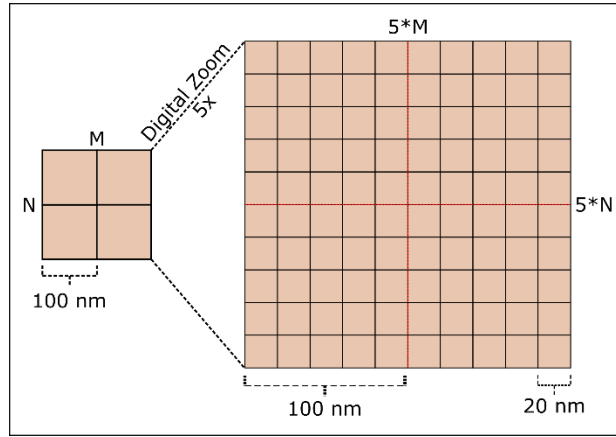

**Figure 4. Scheme of the effect of the amplification value.** Pixel size is defined as the physical distance one pixel covers in the sample. When choosing AMP > 1, said distance is reduced proportionally to AMP, i.e., the number of pixels covering the same distance in the sample is increased. In this example, after an amplification value of 5 is applied, a pixel with a 100 nm size is reduced to 20 nm.

To choose an AMP value, simply solve the next equation based on desired pixel size you want to achieve on the super-resolved image:

$$AMP = \frac{\text{Current Pixel Size}}{\text{Desired Pixel Size}} \quad [\text{Eq. 1}]$$

##### 5.2. FWHM OF PSF

The PSF describes the intensity distribution pattern resulting from the convolution of light traveling through an optical system. Approximately 86% of the light is harbored within the central disk of the Airy pattern, which is defined as the best-focused spot of light that a perfect lens with a circular aperture can make, limited by the diffraction of light. External disks are arranged concentrically, and their intensity decreases as a function of the distance from the center (Figure 5a). The FWHM of PSF parameter represents the number of pixels which fully cover the distance from the maximum of the central disk to the first minimum (innermost dark circle in Figure 5b).

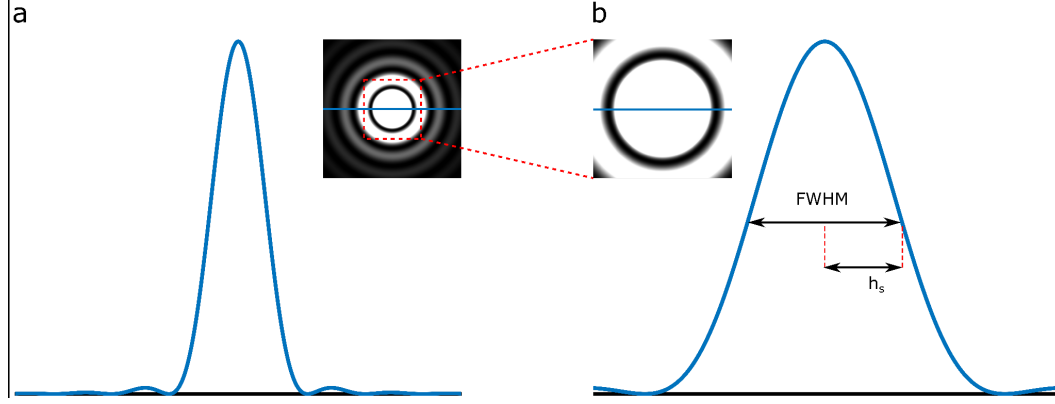

**Figure 5. Graphical description of the FWHM of PSF parameter for the MSSR plugin.** The Airy pattern shown describes a sectional representation of the PSF of the microscope. The FWHM of PSF parameter is a representation of the diameter of the main lobe of the PSF at the point of half its maximum magnitude (denoted as FWHM). This value is tightly related to the optical properties of the microscope and is an indicative of native resolution.

*FWHM* can be provided from direct experimental estimates of the PSF of the optical system, i.e., by computing the width of diffraction-limited objects such as isolated fluorescent beads, for instance. Alternatively, *FWHM* can be estimated from the following known properties of both the optical system used to generate the images and the specific sample under study:

• **Refractive index of the sample medium ($n_s$)** in which the sample is contained.
• **Refractive index of the oil ($n_o$)** placed between the lens and the coverslip.
• **Numerical aperture ($NA_{obj}$)** of the microscope's objective lens.
• **Emission wavelength ($\lambda_{em}$)** (in nanometers) of the fluorophore under study.
• **Pixel size ($p$)** of the input image (in nanometers).

*FWHM* can be computed automatically through the “Compute FWHM of PSF” option (section 3, Figure 3b.1), where the above mentioned parameters must be introduced. The algorithm provides an estimate of *FWHM* (in pixels), from the Rayleigh distance (distance to first minimum) as follows:

$$FWHM = 0.5746 * \frac{\lambda_{em}}{NA}. \quad [\text{Eq. 2}]$$

According to the pixel size the *FWHM* will be:

$$FWHM = FWHM/p. \quad [\text{Eq. 3}]$$

If there is a mismatch between the diffraction index of the sample ($n_s$) and the immersion oil ($n_o$), then the “Compute FWHM of PSF” algorithm performs a correction for the numerical aperture of the objective lens ($NA_{obj}$):

$$NA = \frac{n_s}{n_o} * NA_{obj}. \quad [\text{Eq. 4}]$$

For example, if the imaging lens is a 100X oil-immersion objective with $NA_{obj} = 1.4$, $n_o = 1.515$ and the sample is contained in water media ($n_s = 1.33$), then the corrected NA value is $(1.33/1.515) * 1.4 = 1.229$. In another example, a sample whose membrane labeled with a red fluorescent protein ($\lambda_{em} = 640 \text{ nm}$, $n_s$

= 1.33) is imaged through an oil-immersion objective with a NA =1.4, $n_o = 1.515$, at a final pixel size of 117 nm. Calculation of $FWHM$ using the “Compute FWHM of PSF” option yields $FWHM = 2.56$ pixels.

Another way to estimate a value for the FWHM of PSF is to analyze the frequency space to obtain the maximum observable spatial frequency ($K_0$) in an image, and calculate the inverse of this value, since $K_0 \approx 1/PSF_{FWHM}$. This can be done with the Image Decorrelation plugin for Imagej/Fiji.

##### 5.3. MSSR ORDER

The MSSR algorithm encompasses two main processing stages:

○ MSSR of zero order ($MSSR^0$)
○ MSSR of higher orders ($MSSR^n, n > 0$)

$MSSR^n$ is an iterative process intended to grant further image resolution and contrast enhancement per $n$-iteration step (order). The Order parameter is set to 1 by default. Higher orders provide higher resolution gain at the cost of fluorescence intensity decimation.

##### 5.4. INTERPOLATION TYPE

Part of the extended-, enhanced- and super-resolution image reconstruction process is an interpolation step that fills the gaps in the amplified image. Implemented in the current version of MSSR are two different interpolation algorithms: bicubic and Fourier interpolation. The theoretical foundations of both approaches and their differences are detailed in Supplementary Note 6 of this work, where a comparison of the performance between these algorithms is also provided.

##### 5.5. TEMPORAL ANALYSIS

Prior to a temporal analysis, MSSR operates over single fluorescence images. When given a stack, the result is a single-frame extended-resolved ($sf - MSSR^n$) stack of images. Once this first step is completed, a temporal analysis is optionally performed over the $sf - MSSR^n$ stack, through a Pixel-Wise Temporal Function (PTF), which further increases the resolution, resulting in a single, temporally super-resolved image ($t - MSSR^n$) (Figure 6 and Figure 8).

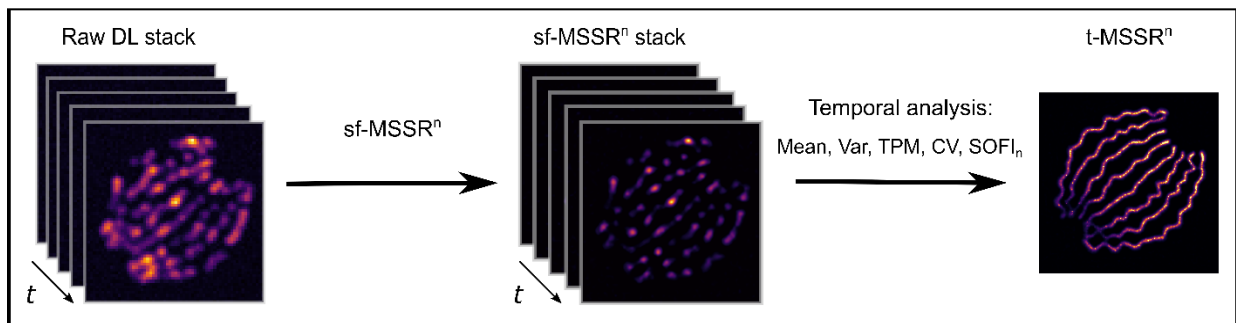

**Figure 6. Temporal analysis of MSSR.** First, a single-frame MSSR analysis over a diffraction-limited stack (left) yields a extended-resolved $sf-MSSR^n$ stack (middle). Next, when a PTF is applied through a temporal analysis, a super-resolved $t-MSSR^n$ frame is obtained.

The blinking nature of the fluorophores can be used as a criterion for the selection of the PTF that best fits your data (Figure 7, Table 1).

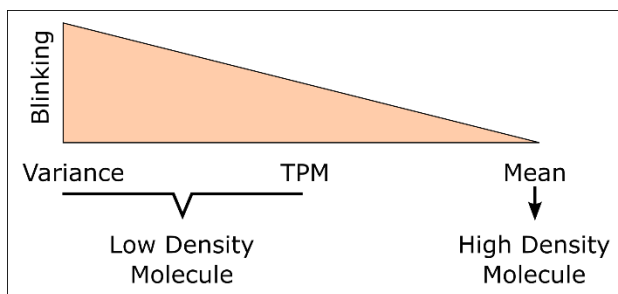

**Figure 7. Blinking nature and density of fluorophores as criteria for PTF selection.**

| PTF Notation | Name | Recommended Usage |
| --- | --- | --- |
| Mean | Average | Low blinking of fluorophores |
| TPM | Temporal Product Mean | Intermediate blinking of fluorophores |
| Var | Variance | High blinking of fluorophores |
| CV | Coefficient of Variation |  |
| SOFI<sub>2</sub> | Auto-cumulant order 2 |  |
| SOFI<sub>3</sub> | Auto-cumulant order 3 |  |
| SOFI<sub>4</sub> | Auto-cumulant order 4 |  |

**Table 1. Available PTFs for the MSSR temporal analysis and their recommended usage.** More rapidly-blinking fluorescently labeled samples will benefit the most from a SOFI temporal analysis, while more information about the fluorescence dynamics will be properly recovered with the Mean or TPM PTFs when blinking is low.

#### 5.6. MINIMIZE MESHING

Occasionally, reconstructions provided by bicubic interpolation can present a mesh-like pattern. When the interpolation process takes place, a gradient of intensities is generated between the original pixels and the ones that are generated; this is because the interpolation function averages values which are close to the point that is being generated. For each new pixel, the closer it is to the original one, the more it will resemble it. This means that, if a new pixel is in an intermediate point between two pixels, its value will be affected by the information of both sides and will most likely be an average of them both. When this process is repeated over a stack with very similar pixel intensities, the new pixels (in intermediate positions) in each image will tend to retain likeness to each other, and thus causing the variance to be rather low. This effect is commonly accentuated when SNR is relatively low. Due to noise being intrinsically and inevitably introduced during the imaging process in any optical system, this

‘meshing’ effect will always take place when real experimental data is analyzed. Therefore, the “Minimize Meshing” option is always enabled (default) when the MSSR plugin is opened, and its use is highly encouraged. Please refer to Supplementary Note 6 of this work for more details.

#### 5.7. GPU COMPUTING

Enabling this option allows the MSSR plugin to scan your system for available GPUs for MSSR analysis. This feature speeds up data processing time (depending on the GPU and the parameters used).

#### 5.8. BATCH (DIRECTORY ANALYSIS)

This feature allows to analyze all the images (separate files) contained in a specific, user-defined path in your system, using a set of previously defined global parameters for all images. When completed, this process generates a new folder inside the specified path, which contains all the resulting $sf - MSSR^n$ images (Figure 9). Note that, when enabled, this option will ignore any image stacks located within the specified path. *Directory analysis* is designed to only analyze as many single images as possible with the same set of parameters (AMP, FWHM, Order).

#### 6. MSSR usage

##### 6.1. EXAMPLE OF MSSR TEMPORAL ANALYSIS

If only a $sf - MSSR^n$ analysis was previously performed over a diffraction-limited stack, one can choose to additionally perform a temporal analysis without re-running the whole process. For the latter, choose the **Temporal Analysis** GUI from the plugin menu (Figures 3d and 8).

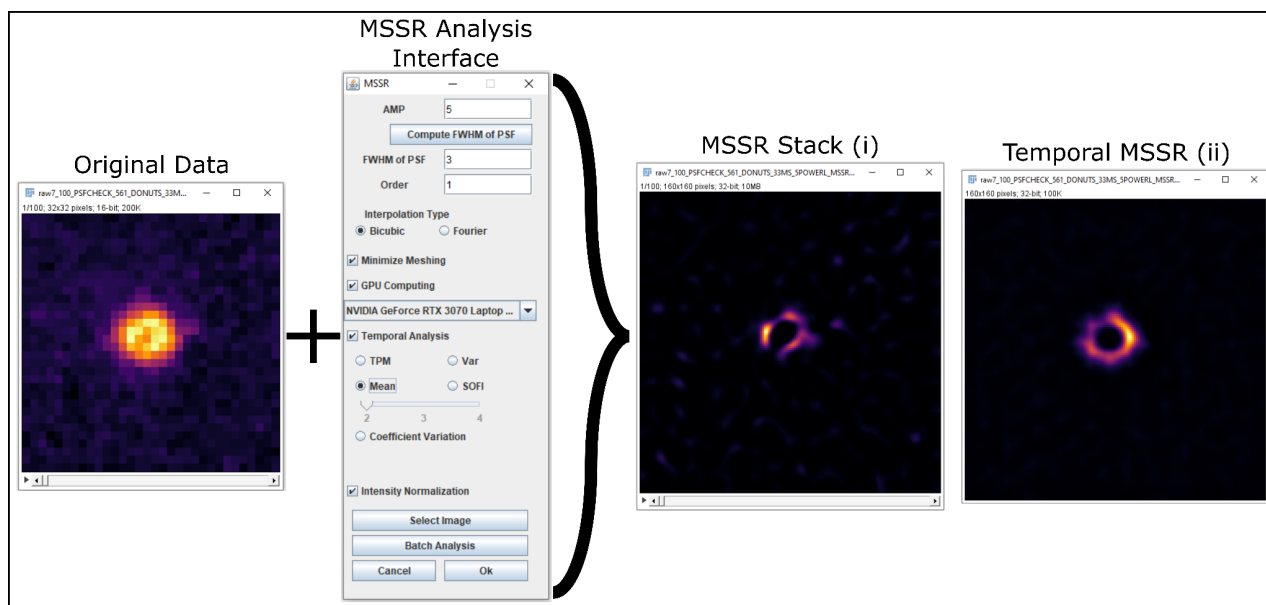

**Figure 8. Example of a MSSR temporal analysis.** Processing a temporal stack of diffraction-limited images through the “MSSR Analysis” GUI when enabling the “Temporal Analysis” option will generate two images: **(i)** a $sf$-MSSR<sup>n</sup> stack and **(ii)** a temporally super-resolved t-MSSR<sup>n</sup> frame. Parameters used in this example: AMP = 5, FWHM = 3, Order = 1, PTF=Mean, Meshing minimization = Yes, GPU computing = Yes.

#### 6.2. EXAMPLE OF MSSR BATCH (DIRECTORY) ANALYSIS

One can choose to perform an analysis on multiple tiff files located in the same folder on the computer, this analysis will take the parameters defined on the plugin to process all the files in the selected folder. The resulting images from this process will be saved in a new folder (named “MSSR”) created in the same location of the analyzed files. This is useful for analyzing data sets that can be processed under the same parameters (amplification, FWHM, order, etc.), and save the trouble of analyzing them one by one.

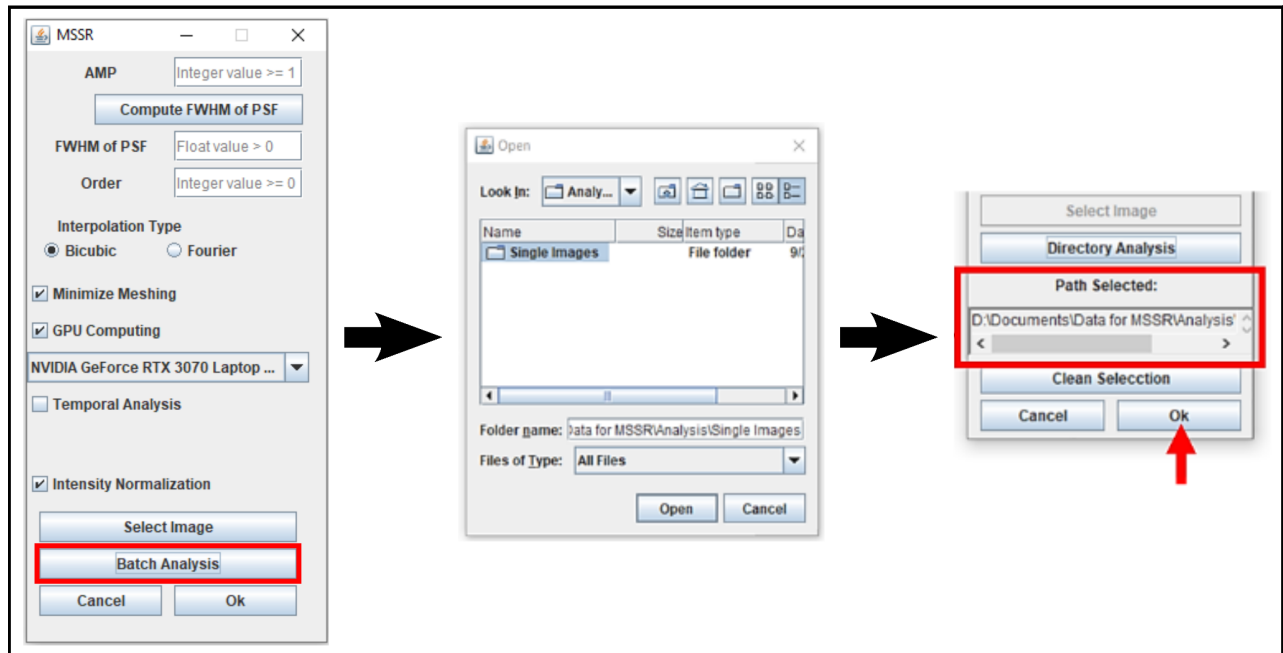

**Figure 9. Example of a MSSR batch (directory) analysis.** Selecting the “Directory Analysis” option (red rectangle, left) will open a new window, where the path to analyze is specified (it will be then displayed on the GUI) (middle); once specified, click “Open”. After all images are successfully analyzed, a folder named “MSSR” will be created within the selected path and will contain all the resulting extended- or enhanced-resolved sf-MSSR<sup>n</sup> images.
