## Supplementary material for "Nanoscopic resolution within a single imaging frame": Online Methods

Torres García E. et. al.,

### Online methods.

*Reagents.*

All chemicals were purchased from Sigma-Aldrich Chemical Co. (St Louis,MO) except otherwise indicated. SiR-actin was obtained from Cytoskeleton (Denver, CO) and FM4-64 was purchased from Thermo Fisher Scientific (Waltham, MA).

AFM Reagents. The Acidaminococcus sp dCas12a protein was expressed in *Escherichia coli* BL21 and purified by chromatography on Ni-NTA (Cytiva), HiTrap SP HP (Cytiva), and HiLoad Superdex 200 16/60 (Cytiva) columns according to recommendations (1) and determined purity through polyacrylamide gel electrophoresis. The 55 nt guide RNA (gRNA) was transcribed in vitro and then purified by TRIzol (Invitrogen) and verified integrity through denaturing urea polyacrylamide gel electrophoresis. The following list of antibodies was used in this study:

- Mouse monoclonal antibody VP4 (2G4) (Harry B. Greenberg, Stanford University. PMID:2431540).
- Mouse monoclonal antibody VP7 (M60) (Harry B. Greenberg, Stanford University. PMID:2431540).
- Mouse monoclonal antibody VP7 (159) (Harry B. Greenberg, Stanford University. PMID:2431540).
- Mouse polyclonal antibody NSP2 (Made by our laboratory, PMID: 9645203; RRID:AB_2802096).
- Rabbit polyclonal antibody NSP2 (Made by our laboratory, PMID: 9645203; RRID:AB_2802097).
- Rabbit polyclonal antibody NSP4 (Made by our laboratory, PMID: 18385250; RRID:AB_2802094).
- Rabbit polyclonal antibody NSP5 (Made by our laboratory, PMID:9645203; RRID:AB_2802098).
- Goat anti-rabbit Alexa 568 (Invitrogen, A-11011).

- Goat anti-mouse Alexa 488 (Invitrogen, A-10680).

*Animals.*

CD1 mature (10- to 12-wk old) male mice were used. Animals were maintained at 23°C with a 12-hours light - 12 hours dark cycle. Animal experimental procedures were approved by the Bioethics Committee of Instituto de Biotecnología, UNAM.

*CRISPR/Cas protein expression and purification.*

Nuclease-dead dCas12a from *Acidaminococcus* sp. fused to an N-terminal 6His-SUMO tag was expressed in Escherichia coli BL21. Cells were grown in Luria-Bertani broth at 37°C and transferred to 12°C when OD600 reached 0.8. After 1h, IPTG was added to a final concentration of 1 mM. After 24 h growth, cell pellets were collected by centrifugation and stored at -70°C until protein purification.

The pellet was thawed in a lysis buffer (20 mM Tris-HCl pH 8.0, 250 mM NaCl, 10 mM imidazole) and sonicated for 6 minutes in 5 s ON-25 s OFF intervals. This was followed by centrifugation at 35,000 g at 4°C for 35 min and filtration using membranes with 0.22μm pore-size. The cell-free extract was injected into a Ni-NTA column (Cytiva) and eluted with an elution buffer: 20 mM Tris-HCl pH 8.0, 1 M NaCl, 250 mM imidazole. The protein was mixed with the SUMO protease and dialyzed overnight at 4°C in dialysis buffer (50 mM phosphate buffer pH 6.0, 100 mM KCl, 5 mM $MgCl_2$, 10% glycerol, 2 mM DTT) to remove the 6His-SUMO tag. The cleaved protein was injected into a cation exchange HiTrap SP HP column (Cytiva) and eluted with a linear gradient from 0 to 50% IEX buffer (20 mM HEPES-KOH pH 7.2, 100 mM KCl, 5 mM $MgCl_2$, 10% glycerol, 2 mM DTT). The protein was further purified by size exclusion chromatography on a HiLoadSuperdex 200 pg 16/60 column (Cytiva) in storage buffer (20 mM HEPES-KOH pH 7.5, 500 mM KCl, 10% glycerol) and aliquots were stored at -70°C until use. Purity and identity of dCas12a were confirmed via polyacrylamide gel electrophoresis and western-blot against the 6xHis region on dCas12a.

*Production of crRNAs.*

The crRNA used in atomic force microscopy (AFM) experiments was produced by in vitro transcription of DNA templates previously amplified by PCR. The templates were produced using two self-complementary oligos (purchased from IDT).

Fw 5'-GAAATTAATACGACTCACTATAGGTAATTTCTACTCTTGTAGAT-3' and

Rv 5'-CCCTGGTCAACCAGGTGAACAAGGATCTACAAGAGTAGAAATT-3'.

In vitro transcription and crRNA purification were performed using HiScribe T7 (NEB) and RNA Cleanup kits (NEB), respectively. The crRNA pool for DNA curtains was produced according to the following procedure. Partially double stranded DNA templates for in vitro transcription were obtained by hybridizing a 24 nt long forward oligo encoding the promoter for T7 RNA polymerase:

5' GAAATTAATACGACTCACTATAGG,

with a pool of five 68 nt long reverse oligos (purchased from IDT):

5'-AUGAUGUUCUGCUGGAUAUGCACU-3',

5'-CCUGACACCGGACGGAAAGCUGAC-3',

5'-AAUGUCGGCUAAUCGAUUUGGCCA-3',

5'-GCUAGCAAUUAAUGUGCAUCGAUU-3' and

5'-AUGAACGCAAUAUUCACAAGCAAU-3',

which encoded the crRNA sequence and the region complementary to the T7 promoter. Forward and reverse oligos were annealed at 1.5:1 ratio (Fw:Rv) in 10 mM Tris-HCl, pH 7.5, 50 mM NaCl, 1 mM EDTA buffer, incubated at 75°C for 5 min and cooling to 25°C during 25 min. HiScribe T7 kit was used for in vitro transcription and the crRNA pool was purified with TRizol (Ambion).

*CRISPR/dCas12a nanoruler preparation.*

First, the CRISPR/dCas12a ribonucleoprotein complex was formed. On a 0.6 mL microcentrifuge tube, the following reagents were added (final concentration): 1X CRISPR action buffer (TRIS-HCl 200 mM, NaCl 500 mM, DTT 5 mM) and dCas12a

20 nM. The gRNA was pre-heated at 90°C for 1 minute and then let cool to room temperature. Upon gRNA cooling, it was added to the dCas12a to a final concentration of 30 nM. The CRISPR/dCas12a components were incubated for 20 minutes at room temperature. When the complex was formed, the target dsDNA was added and the microtube was incubated at 37°C for 1 hour so the CRISPR//dCas12a complex binds to the target sequences in the dsDNA and thus forms the CRISPR/dCas12a nanoruler.

*CRISPR/dCas12a nanoruler slide preparation for fluorescence microscopy.*

The sample imaging volume was delimited by a perforated double-sided tape attached to a coverslip and a slide on each side. The following reagents were prepared on an independent microtube: CRISPR/dCas12a nanorulers (described in the previous section), 20µg/mL Hoechst33342, and 5 nM ssDNA fluorescent probe PS3 [1] (5'-TCCTCCC-3'-ATTO 647N, Integrated DNA Technologies') and graded with dd$H_2O$. The mix was transferred to the perforated double-sided tape in the coverslip and covered with the slide. The following sequence represents the dsDNA used for CRISPR/dCas12a binding, in which **bold** sequences corresponds to the association sites and *italics* are the PAM sequence:

AATTCTTAGGCACCCTTCTTTTTCTTCTTCTTTTTCTTCTTTTTCTTAGCACCTTGGCCGGC
TCCAGCACCGGCTCCTTGACCAGCACCAGCACCAGCACCTTGGCCGGCTCCAGCACCGGCTC
CTTGACCAGCACCAGCACCAGCACCTTGGCCGGCTCCAGCACCGGCTCCTTGACCAGCACCA
GCACCAGCACCTTGGCCGGCTCCAGCACCGGCTCCTTGACCAGCACCAGCACCAGCACCTTG
GCCGGCTCCAGCACCGGCTCCTTGACCAGCACCAGCACCAGCACCAGCACCGGCTGGACCCT
GGTTTCCTGG*TTTA***CCTTGTTCACCTGGTTGACCAGGG**TTACCTGGCTGACCAGGGGAACCT
TGGTTACCTGGAGAGCCTTGTGAACCTGGGGATCCAGGTTGACCATTCTTTCCAGGGTTACCC
TGAGAACCTTGTGGACCGTTGGAACCTGGCTCACCAGGTTGTCCGTTCTGACCAGGTTGACC
AGGTTGACCTTCGTTTCCTGGTTGACCTGGATTACCTGGAGAACCCTTGTTACCGGGCTGTCC
TTGGTTACCAGGAGATCCTGGGTTACCTGGCTCACCGGCTGGACCCTGGTTTCCTGG*TTTA***CC**
**TTGTTCACCTGGTTGACCAGGG**TTACCTGGCTGACCAGGGGAACCTTGGTTACCTGGAGAG
CCTTGTGAACCTGGGGATCCAGGTTGACCATTCTTTCCAGGGTTACCCTGAGAACCTTGTGGA
CCGTTGGAACCTGGCTCACCAGGTTGTCCGTTCTGACCAGGTTGACCAGGTTGACCTTCGTTT
CCTGGTTGACCTGGATTACCTGGAGAACCCTTGTTACCGGGCTGTCCTTGGTTACCAGGAGAT
CCTGGGTTACCTGGCTCACCGGCTGGACCCTGGTTTCCTGG*TTTA***CCTTGTTCACCTGGTTG**
**ACCAGGG**TTACCTGGCTGACCAGGGGAACCTTGGTTACCTGGAGAGCCTTGTGAACCTGGG
GATCCAGGTTGACCATTCTTTCCAGGGTTACCCTGAGAACCTTGTGGACCGTTGGAACCTGGC
TCACCAGGTTGTCCGTTCTGACCAGGTTGACCAGGTTGACCTTCGTTTCCTGGTTGACCTGGA

TTACCTGGAGAACCCTTGTTACCGGGCTGTCCTTGGTTACCAGGAGATCCTGGGTTACCTGGC TCACCGGCTGGACCCTGGTTTCCTGG*TTTA***CCTTGTTCACCTGGTTGACCAGGG**TTACCTGG CTGACCAGGGGAACCTTGGTTACCTGGAGAGCCTTGTGAACCTGGGGATCCAGGTTGACCAT TCTTTCCAGGGTTACCCTGAGAACCTTGTGGACCGTTGGAACCTGGCTCACCAGGTTGTCCGT TCTGACCAGGTTGACCAGGTTGACCTTCGTTTCCTGGTTGACCTGGATTACCTGGAGAACCCT TGTTACCGGGCTGTCCTTGGTTACCAGGAGATCCTGGGTTACCTGGCTCACCGGGTGCACCAG CACCGAGACCACAAGCTTCAGCTTCTCTCTTCTCGAGAGAT 3’.

*GATTA-PAINT and CRISPR/dCas12a nanoruler sample imaging.*

The GATTA-PAINT 40 RG nanoruler was provided as a single slide ready for imaging (GATTAquant DNA nanotechnologies). It has three fluorophores at a separation of 40 nm between them (ATTO 542/ATTO 655) and 80 nm between the furthest. Imaging was performed on an Olympus IX-81 inverted microscope using total internal reflection fluorescence (TIRF) illumination with a penetration depth of 200 nm (Olympus, cellTIRF Illuminator). Images were collected with an iXon 897 EMCCD camera (Model No. DU-879-CS0-#BV). A set of 300 frames were acquired at an exposure time of 50 ms per image, excitation laser of 488 and 561 nm with full laser power (23.1 mW measured at the back focal plane of the lens), and an effective pixel size of 160 nm in the object plane (Olympus UApo N 100X / 1.49 numerical aperture, oil-immersion). For MSSR only the first 100 frames were analyzed.

The CRISPR/dCas12a nanoruler sample was visualized on the same imaging setup as the GATTA-PAINT 40RG nanoruler, except that a 20 ms as acquisition time was employed. Nearby emitters were automatically identified from t-MSSR$^3$-Var images using the Maximum Finder function of FIJI/ImageJ. Maxima were accepted only if their intensity value (DL, digital levels) was higher than a threshold value (prominence = 1900 DL), in comparison with the intensity values from the ridge to a higher maximum. The coordinates of the identified local maxima (emitter’s location) were computed from 16 regions of interest (1.5 $\mu m^2$ each) and exported to R to further quantify the intermitter’s distances considering a worm-like chain model [2]. Briefly, the intermitter distances were computed for any pair of identified local maxima within the same t-MSSR$^3$-Var image. The CRISPR/dCas12a nanoruler system was design to with four binding sites for dCas12a distributed uniformly every 297 bp (equivalent to ~ 100 nm), hence, two emitters are considered to be part of the same dsDNA if their intermitter distance is shorter than the accumulated distance of four binding sites (300 nm). All measured intermitter distances were pooled on a single histogram and fitted in the context of Gaussian mixture models. Fitting was performed in R with the normalmixEM routine of mixtools with parameters $\mu$: $\{\mu_1 = 100, \mu_2 = 200, \mu_3 = 300\}$ nm, and $\sigma = \text{sqrt}(\mu)$.

*AFM visualization.*

The ribonucleoprotein particle (RNP) was assembled from dCas12a and crRNA (1:1.5 molar ratio) in AFM buffer (20 mM Tris-HCl pH 8.0, 100 mM NaCl, 15 mM $MgCl_2$, 1 mM DTT) at 37°C for 20 min. The DNA template (1,500 bp) with four dCas12a target sites was added to the mix at 40:1 (RNP:DNA) molar ratio and incubated for 1 h at 25°C. The sample was diluted 5-fold to a final concentration of 1 nM DNA and deposited on a freshly cleaved mica for 10 min, followed by rinsing with 0.5 mL filtered milli-Q water and air-drying. Images were acquired with an atomic force microscope (NanoScope V, Bruker) on ScanAsyst-Air mode at room temperature and 1,024 samples/line. Images were processed with NanoScope Analysis Software v1.89.

*DNA curtain assay.*

DNA from bacteriophage λ (λDNA) (NEB) was mixed with biotinylated oligos complementary to the cohesive ends of λDNA in reaction buffer for T4 DNA ligase (NEB), incubated at 70°C for 15 min and cooled down to 15 °C, over 2 h. T4 DNA ligase was used for overnight ligation at room temperature. After ligase inactivation with 2 M NaCl, the biotinylated DNA was purified on a Sephacryl S-1000 size exclusion column (GE Healthcare).

The flowcell was passivated with a lipid solution (1.954% DOPC, 0.04% DOPE-mPEG2k and 0.006% DOPE-biotin) in buffer (10 mM Tris-HCL pH 8, 100 mM NaCl) for 30 min at room temperature. The flowcell was washed with BSA buffer (40 mM Tris-HCl pH 8, 2 mM $MgCl_2$, 0.2 mg/mL BSA) and incubated for 10 min. The biotinylated DNA in BSA buffer was injected into the flowcell and non-tethered DNA was washed out. BSA buffer supplemented with 100 mM NaCl, 5 mM $MgCl_2$, 2 mM DTT was used for imaging.

Ribonucleoprotein particles were prepared by mixing dCas12a with the crRNA pool at 1:10 molar ratio in buffer (20 mM Tris-HCl pH 8.0, 100 mM NaCl, 5 mM

$MgCl_2$, 2% glycerol, 2 mM DTT) at 37°C for 30 min. The complex (10 nM) was injected into the flowcell for DNA binding during 30 min at room temperature. Labeling of dCas12a was achieved with anti-FLAG antibodies conjugated to quantum dots (QD705). C-S10-B was labeled with maleimide-Alexa488 at a single N-terminal cysteine.

Images were acquired at 60X with an inverted Nikon Ti-E microscope with 488 nm excitation laser. Emission was split with a 638 nm dichroic beam splitter (Chroma) and captured by two EM-CCD cameras (Andor iXon DU897). Images were processed with FIJI [3].

*Structured-illumination microscopy.*

The GATTA-SIM 140B nanoruler was provided as a single slide ready for imaging (GATTAquant DNA nanotechnologies). Spreads of germ cell chromosomes were performed according to Faieta [4]. In brief, testes were removed from euthanized animals, decapsulated, macerated in high-glucose MEM and mixed. The suspension was left to settle, and the supernatant was spun down at 7200 rpm for 1 min. The pellet was resuspended in 0.5 M sucrose and the suspension was added to slides coated with 1% paraformaldehyde in 0.015% Triton X-100 and incubated for 2 h in a humidified chamber at room temperature (RT). At the end of the incubation, slides were rinsed twice in 1:250 Photo-flo Kodak professional (no. 1464510) in water and allowed to air dry. Surface chromosome spreads were either immediately processed for immunofluorescence or stored at −80°C for up to 6 months. SYCP3 was stained using a primary antibody from Santa Cruz SC-74569 (SYCP3 D1) and a secondary antibody anti-mouse Alexa 568 (A11004 Thermo). Imaging was performed using an Elyra 7 microscope (Zeiss). Image acquisition was made with a 60x 1.4 NA oil immersion objective and a 1.4x lens as extra magnification. Image reconstruction was done in ZEN Black with default parameters.

*Rotavirus cell infection and immunofluorescence.*

The cell culture and infection were performed as described previously [6]. Briefly, MA014 cells (American Type Culture Collection; ATCC:CRL-2378.1;

RRID:CVCL_3846) grown on glass coverslips were infected with rotavirus RRV (obtained from Harry B. Greenberg, Stanford University) at a multiplicity of infection (MOI) [6]. At six hours post infection the cells were fixed with and processed for immunofluorescence. Finally, the coverslips were mounted onto the center of glass slides with a STORM buffer mounting medium (1.5% glucose oxidase + 100 mM β-mercaptoethanol).

*Optical setup for Rotavirus replication machinery visualization.*

All images were kindly provided by Garces and collaborators [6]. Briefly, images of the rotavirus viroplasms were acquired on an Olympus IX-81 inverted microscope configured for total internal reflection fluorescence (TIRF) excitation (Olympus, cellTIRFM illuminator) using a critical angle such that the evanescence field had a penetration depth of 200 nm. The fluorophores Alexa Fluor 488 and Alexa Fluor 568 were excited with light of 488 nm and 568 nm respectively, using a laser-modulation protocol as described previously in [6]; the optical setup consists of an Olympus UApo N 100x 1.4 NA, oil-immersion objective lens, with an extra 1.6x intermediate magnification lens. The images were acquired by an EMCCD camera (iXon 897, Model No: DU-897E-CS0-#BV; Andor) at a frequency of 20 fps and effective pixel size of 100 nm at the object plane. MSSR processing was performed considering the following parameters: AMP = 5, PSF = 3, Order = 1. GPU parallel computing was enabled, and 100 images were used with t-MSSR-*Mean*.

*Live imaging of sperm acrosomal exocytosis and F-actin dynamics.*

The non-capacitating medium (NC) used was a modified Toyoda–Yokoyama–Hoshi (modified TYH) which contains 119.3 mM NaCl, 4.7 mM KCl, 1.71 mM $CaCl_2.2H_2O$, 1.2 mM $KH_2PO_4$, 1.2 mM $MgSO_4.7H_2O$, 0.51 mM sodium pyruvate, 5.56 mM glucose, 20 mM HEPES and 10 µg/ml gentamicin. For capacitating conditions 15 mM $NaHCO_3$ and 5 mg/ml BSA were added (CAP).

Animals were euthanized and cauda epididymal mouse sperm were collected. Both cauda epididymis were cut at multiple sites and placed in 500 µl of NC. After 15 min incubation at 37°C the epididymis were removed. Sperm were pre-incubated for 10 minutes in the presence of 100 nM SiR-actin in NC. Once loaded, sperm were incubated for another 60 minutes in CAP, the concentration of SiR-actin was 100 nM during the whole experiment.

Sperm were immobilized on concanavalin-A (1 mg/ml)-coated coverslips. The chamber was then filled with a recording medium (NC) containing 100 nM SiR-actin and 0.5 µM FM4-64. 100 images were obtained every 30 seconds for 20 minutes using the NanoImager S microscope (Oxford Nanoimaging Ltd), equipped with a 100X, 1.4 NA, oil-immersion objective (Olympus). For SiR-actin excitation, a 640 nm laser was used and for FM4-64 excitation a 561 nm laser was used. Effective pixel size at object plane = 117 nm.

*Plant Material and Growth conditions.*

*Arabidopsis thaliana* seeds were surface sterilized, germinated and grown in 0.2× Murashigue and Skoog medium (prepared based on Linsmaier and Skoog medium L477; PhytoTechnology Laboratories, Lenexa, KS, USA), pH 5.7, supplemented with vitamins (0.1 mg l-1 pyridoxine, 0.1 mg l-1 nicotinic acid), 1% sucrose, and 0.8% agar. The plants were grown in a chamber at 21°C, 16/8 h light/dark photoperiod and a light intensity of 105 µmol photons $m^{-2}s^{-1}$.

*Volumetric imaging of A. thaliana root cells.*

Confocal imaging of the double transgenic line, an F1 of a cross between plasmalemma pUBQ10::NPSN12-YFP [7] and nuclear p35S:H2B:RFP [8] marker lines, was performed with a Zeiss Axiovert 200M microscope equipped with a C-APO ×63, 1.2NA objective (Oberkochen, Germany) and a coupled confocal system with a 488-nm laser source, a filter cube with 525/45 nm and 630/92 nm bandpass filters for yellow and red fluorescent protein emission, respectively, and a linear motor travel XY Stage and a Z-axis piezo stage with controllers (Thorlabs, Inc. Newton, NJ, USA). The XY pixel size of 404 nm and Z step size of 500 nm were implemented. The final image in Figure S30 a, b and c is of a Z-projection of ten slices with SUM slices mode and the FIJI LUT used for red and green channels were “Cyan Hot” and “Magenta”, respectively.

A nuclear marker line, p35S:H2B:RFP, was imaged with an inverted Olympus FV1000-IX81 confocal microscope equipped with a LUMFLN×60, 1.3NA S objective. The 543 nm laser was used to excite RFP and emitted light was filtered with BA560-660. The oversampled XY pixel size of 41 nm and Z step size of 100 nm were implemented. The final image in Figure 29d was made by a Z-projection of 86 slices with MAX intensity mode and the LUT used was "Royal" of FIJI [3].

Selective plane illumination microscopy (SPIM) imaging of *A. thaliana* root cells was performed over a transgenic primary root expressing p35s:H2B-R with an in-house SPIM system inspired on the OpenSPIM project [9], with some setup modifications of the original design. Briefly, the illumination path consists of a C-flex laser combiner providing laser excitation sources at 405, 488, 561, 638 nm (Hubner Photonics, Cobolt Series 01-06, DPL for 561 and MDL for 405,488,638), which is coupled to the SPIM optics through a single multimode laser guide (Fiber optic with FC-APC output). Laser light is focused on a horizontal plane shaped via cylindrical lens (Thorlabs ACY254-050-A, f=50.0m ±1%, Ø 25.4mm, AR Coating: 350 - 700 nm). The focal plane of the cylindrical lens is imaged by a telescope in the back focal plane of the illumination objective (Olympus UMPLFLN10XW, 10X water immersion, NA=0.3mm WD=3.5mm). The excitation light-sheet is confined within the imaging area by a slit (Thorlabs VA100/M, Adjustable Mechanical Slits, Internal thread =2.4mm), which is placed in the center of the telescope. The resulting light-sheet has a beam waist of about 3 μm in the focal plan of the illumination objective. The detection unit consists of a 20X water immersion objective (Olympus UMPLFLN20XW, NA=0.5mm WD=3.5mm), a tube lense (Ø 60mm x 104mm), a multi-bandpass emission filter set (Semrock, FF01-446/523/600/677-25 BrightLine, Ø 25mm x 3.5mm), and a sCMOS camera 82 % peak QE (Hamamatsu ORCA-Flash4.0 V2 - Camlink 100fps). A 3D printed water filled imaging chamber (internal volume = 22x22x30mm without objectives) embodies the illumination objective and the detection objective aligned at 90°, a custom-made 3D printed sample holder and the sample.

Fluorescence excitation of the H2B-R-RFP expressing root cells was provided via the 561 nm laser light using stroboscopic illumination. SPIM volumetric imaging was achieved by mounting the sample on a four dimensional (XYZ, and Y rotation) motorized stacked stage (Picard Industries, USB 4D Stage, linear range = 9mm, Includes Sample-Arm). Computer control of stroboscopic illumination, image acquisition and sample translation were provided by the OpenSPIM plugin 64-Bits of μmanager (v.1.4 for windows) [10]. Images were collected at a final pixel size of 0.325μm, a z-step of 1.524μm and a rotation step of 1.8°. MSSR processing was performed considering the following parameters: AMP = 10, PSF = 2, Order = 0.

*Live-cell imaging of LLC-PK1 cells microtubule dynamics.*

LLC-PK1 cells stably expressing mEmerald-EB3 were cultured and imaged using an ORCA-Fusion back-thinned sCMOS camera (Hamamatsu, C15440-20UP) and a 100X/1.47 NA oil-immersion objective (Plan-Apochromat, Zeiss) in a Zeiss Celldiscoverer 7 microscope with $CO_2$ and temperature control set at 5% and 37°C, respectively. Fluorescence excitation was provided by a 488 nm laser at 1% laser power and emission light ($\lambda_{em}$ = 510 nm) was collected using a FITC filter. Image collection was done using the ZEN 3.2 (blue edition) acquisition software, with an exposure time of 100 ms, 2 $s^{-1}$ frame rate and a 43 nm pixel size. Stable cell lines were generated and provided by Michael W. Davidson [11,12].

*STED microscopy.*

Immunofluorescence imaging was carried out using a STEDYCON mounted on an upright Zeiss microscope in confocal or STED modes. Samples were imaged with a Zeiss 100x 1.46 NA objective, 20 nm pixel size, 5 us pixel dwell time, 15-line accumulations and a pinhole of 64 um**.** Immunofluorescence was performed as in [13]. Primary antibodies used were anti-H3K27me3 (Abcam ab6002) and anti-H3K27ac (Active Motif, 39034), both at 1:200 dilution. Secondary antibodies used were anti-mouse labeled with STAR Red and anti-rabbit labeled with STAR Orange. STED laser powers were 3% of the 640 nm and 775 nm 96.5% for the STAR red channel, whereas for the STAR Orange channel was 7.8% of the 561 nm laser and 100% for 775 nm.

*ArgoLight "Argo-SIM" test slide.*

Confocal images were acquired on a Zeiss LSM880 inverted microscope using a Plan-Apochromat 63x/1.4 Oil immersion objective, exciting the micropattern with the laser 405 nm and detecting the fluorescence in the range 420-480 nm. The pixel size was 0.044 micrometers. The same area has been acquired using the Airyscan detector with the same settings for laser power, detector gain, image format, bit depth, and line averaging, using a selective optical filter BP 420-480 + LP 605, and processing the images with the Airyscan algorithm set on strength parameter 6.

*PSFcheck imaging.*

We employed an immobile fluorescence pattern as a calibration sample for SRM [5]. The fluorescent patterns were fabricated using direct laser writing via infrared ultrashort-pulses that create regions of autofluorescence in a two-part epoxy mixture polymer sandwiched between a coverslip and a microscope slide (PSFcheck) [4]. One of the patterns present on a PSFcheck calibration slide consists of a 3D array of small diffraction limited shell features separated 10 μm from each other. The fluorescent thickness of a shell is small compared to the PSF so that the average FWHM across several features was calculated to be of 208 nm in a SIM microscope. We imaged this shell pattern using widefield fluorescence excitation on a NanoImager-S (Oxford Nanoimaging Ltd), equipped with a 100X, 1.4 NA, oil-immersion objective (Olympus). The PSFcheck sample was excited with a 561 nm laser and the emitted fluorescence acquired in the Emission Filter 2: Band 1 575-616.5 and recorded on an sCMOS Hamamatsu Orca Flash 4.0 V3. Acquisition time = 33 ms, effective pixel size at object plane = 117 nm.

*Automatic Correction of sCMOS-related Noise (ACsN).*

ACsN is a noise correction method for sCMOS images that uses a principle of similarity between patches within the same image to characterize noise using 3D filtering [19]. The camera noise together with the signal from the incident photons can be represented by a distribution whose standard deviation is approximated based on the frequency thresholds of the modulation transfer function (MTF). This threshold is calculated from the optical parameters of the system. The ACsN application was used in Matlab 2020a. The input parameters were 1.4 of numerical aperture, wavelength of 610 nm, and pixel size of 117 nm, without video filter and with parallel computation.

*ORCA Flash 4.0 V3 sCMOS detector characterization.*

The fixed noise patterns of a sCMOS detector characterize the offset, variance and gain of each pixel in so-called calibration maps. The offset and variance are the average and variance, respectively, of the digital pixel-wise values that result from a video where no photons hit the detector. The gain is a multiplicative value of the signal when photons are detected. To characterize the maps of the sCMOS ORCA Flash 4.0 V3 detector, a code was written in the R programming language that implements the previously described calibration [20].

A video of 60 thousand images was taken without illumination on the NanoImager-S microscope (Oxford Nanoimaging Ltd) with no sample or laser turned on. In addition, a uniform fluorescent sample was used to recreate a uniform illumination of the detector and 5 videos of one thousand images each were taken. Each video had an average number of photons, chosen between 20 and 200 photons per pixel as described by Huang et al.

*SNR calculation for raw data.*

From the stacks of 100 images limited by diffraction, the average number of electrons per pixel was estimated for each image based on the following equation:

$$electrons_i = \frac{I_i - O_i}{G_i}$$

Where O and G are the offset and gain maps, respectively. The signal-to-noise ratio (SNR) was calculated with the following equation:

$$SNR = \frac{QE*S}{\sqrt{QE*\left(S+I_b\right)+N_r^2}} = \frac{electrons}{\sqrt{electrons+readout\ noise^2}},$$

where S are the photons per pixel and $I_b$ is the signal in the background. A quantum efficiency (QE) of 0.72 (to calculate photons) and reading noise $N_r$ = 1e- were used. S was considered as the average of photons in the region where the fluorescent ring is located, while $I_b$ is the average value of the pixels that belong to the background of the image.

*Entropy-based Super Resolution Imaging (ESI).*

ESI is a super resolution microscopy method that calculates the entropy of a sequence of fluorescence images and generates a magnified image that contains the actual information of the fluorophores. ESI is available as a plugin for ImageJ [14]. The ESI implementation allows you to create a super resolution image with a magnification of 2x the original size of the input images. The algorithm was iterated twice to achieve a magnification of 4. The input parameters are the number of final images in the output data; the number of bins per entropy, that is, the number of bins in the intensity histogram values for the entropy; and the order of the central moment.

For the first iteration of the algorithm the sequence of 100 images was used as input data and the parameters were used: 50 images in result, 2 bins for entropy and order 0. The second iteration used the 50 images resulting from the first iteration with parameters: 25 images in result, 2 bins per entropy and order 0. The ESI plugin returns the specified number of images and the average image. The average image from the second iteration is the image that is used for subsequent analyzes.

*Multiple Signals Classification Algorithm (MUSICAL).*

MUSICAL is a super resolution method, implemented as a plugin for ImageJ [15], which improves resolution by singular values decomposition of a set of images taken from the same scene. This decomposition results in a collection of eigen-images and their respective eigen-values where each eigen-image characterizes a specific pattern present in the image and the eigen-value associated with that pattern is a statistical measure of the presence of that pattern in the underlying image. The signal from fluorophores in the scene is associated with patterns whose eigenvalues are large, while noise and background are associated with patterns whose eigenvalues are small. A predefined threshold divides the eigen-image set into range space (signal) and null space (noise).

The sequence of 100 images of PSFcheck was used with parameters: 610 nm as emission wavelength, 1.4 numerical aperture, 1 in the magnification of the objective (digital size of the pixel is known), and 117 nm of pixel size, and 4 subpixels per pixel . The threshold value was -0.8 and was chosen from the singular value plot calculated by the plugin.

*Super Resolution Radial Fluctuations (SRRF).*

SRRF overcomes the theoretical limit of diffraction since the emission of a fluorophore has a radial symmetry that can be detected by calculating the convergence of the gradient on a magnified version of the diffraction-limited image [16]. The degree of convergence for each sub-pixel is captured on a radiality map. In this first step, each diffraction-limited image has its corresponding radiality map, while in the second step, these maps are analyzed with a temporal function that improves the final resolution.

SRRF is implemented in an ImageJ plugin [17]. The parameters used for this algorithm were: 0.5 in ring radius, magnification of 4 and 6 axes in the ring. The rest of the parameters were taken by default.

*Super-resolution Quantitative Image Rating and Reporting of Error Locations (SQUIRREL).*

SQUIRREL is an algorithm, implemented as an ImageJ plugin [18], that calculates the global Error (RSE) and Pearson correlation (RSP) values of the Superthe of Super Resolution against its reference limited by diffraction. The RSP and RSE values were calculated using as reference the average image of the diffraction limited images used in the super resolution analysis for each case. These global resolution indexes are a measure of how confiable is the reconstruction related to the reference image.

*Simulation of fluorescent emitters.*

All 1D and 2D simulated emitters used for the examples shown in the main manuscript (Figures 1-2) and the Supplementary Information (Figures S6-10) were generated in Matlab. The Gaussian and Bessel distributions of emitters were generated using the Gaussian ($G_{PSF}$) and Bessel ($B_{PSF}$) PSF, respectively, following the formulas:

$$G_{PSF} = exp\left(-\frac{(x-x_c)^2+(y-y_c)^2}{2\sigma^2}\right)$$

Where σ is the standard deviation, $(x, y)$ are the generated coordinates and $(x_c, y_c)$ is the center of the distribution.

$$B_{PSF} = I_0\left(2\frac{besselj(1,\nu)}{\upsilon}\right)^2$$

Where $I_0$ is the maximum intensity of the distribution, υ is dimensionless distance and $besselj(1, \nu)$ is the Bessel function of first kind with dimensionless parameter υ. To achieve enough spatial detail for visualization of the Gaussian emitter distribution, σ = 10 pixels was used in a square grid of size 81x81 pixels ($x$ = -4σ:4σ, $y$ = -4σ:4σ), with a step size of 1 pixel.

Given that the generated Bessel PSF is undefined by zero division at the center ( $x = 0, y = 0$), its value is set to maximum intensity $I_0$ at this location. The dimensionless distance $v$ was computed following $\nu = k\frac{NA}{n}q$, where λ is the emission wavelength, $NA$ is the numerical aperture, $n$ is the refractive index of the medium, $q$ is the radial distance to the distribution center and $k$ is the wavenumber, given by $k = \frac{2\pi}{\lambda}$.

For 1D emitters, $q = x$ and a 1D grid ranging from $-10^{-6}:10^{-6}$ was used. For 2D emitters, $q = \sqrt{x^2 + y^2}$ and a 2D grid of size $x = -10^{-6}:10^{-6}$, $y = -10^{-6}:10^{-6}$ was used. Note that, in either case, a step size is of $1\ pixel = 1\ nm = 10^{-9}\ m$ was used.

*Dip computation.*

Since the Gaussian distribution is fitted with good accuracy to a Bessel pattern, its use is sufficient to simulate the emitters. First, two Gaussian distributions with centers positioned at different locations along $(-x_c, 0)$ and $(x_c, 0)$ were simulated (using $\sigma = 10$ pixels). The distributions were then added and the dip was computed as the intensity value at the center of the resulting distribution. The dip values in Figures 1c and 2b of the main document were calculated by increasing the distance between the two emitters' distribution centers from 0 to 4σ.

*Image Decorrelation (ImDecorr).*

ImDecorr is an algorithm to compute resolution in a single super-resolved image [21]. Its principle is based on partial phase autocorrelation by applying a mask filter and calculating cross-correlation coefficients in Fourier space. Its implementations are available in Matlab and a plugin for ImageJ. It is a fast, friendly and easy-to-use tool free of user optical parameters to analyze both real data. In this work we used the plugin version of this algorithm.

*Single particle Tracking.*

Single particle tracking was performed on simulated images generated in Fiji/ImageJ. Three different levels of SNR : 2, 4, 7 and three density levels of sub-diffraction particles: low, mid, high: 100, 500, 1000 particles per imaging field were used to simulate the images. Three classes of tracking algorithms were tested in TrackMate v7.6.1 [22]:

(i) LAP: the LAP framework for Brownian motion [22],
(ii) LM: a linear motion tracker based on a Kalman filter [23, 24] and
(iii) NN: a tracker based on Nearest neighbors [25–27].

Particles were identified using the Laplacian of Gaussian (LoG) detectors, where 2 pixels were used as diameter of particles. For SNR = 2, LoG detection results were dominated by noise, hence, the histogram of detection quality was used to select a threshold that yielded the expected particle number of the dataset. For SNR > 2, the detection quality histogram was bimodal, so the threshold was selected at the dip between distributions.

Parameters for tracking algorithms were:
LM: initial search radius = 10, search radius = 7, max frame gap = 3.
LAP: max linkage distance = 7, max gap-closing distance = 10, max frame gap = 3.
NN: max search distance 10 to the nearest neighbor.

Tracking performance was assessed with the “tracking performance evaluation tool” deployed at Icy Icy 2.4 (http://icy.bioimageanalysis.com).

### References

1 Schueder, F. et al. “An order of magnitude faster DNA-PAINT imaging by optimized sequence design and buffer conditions,” Nat. Methods 16, 1101–1104 (2019).

2 Wang, H. & Milstein, J. N. Simulation Assisted Analysis of the Intrinsic Stiffness for Short DNA Molecules Imaged with Scanning Atomic Force Microscopy. PLOS ONE **10**, 1–11, (2015).

3 Schindelin, J. et al. Fiji: an open-source platform for biological-image analysis. Nat. Methods **9**, 676–682, (2012).

4 Faieta, M. et al. A surge of late-occurring meiotic double-strand breaks rescues synapsis abnormalities in spermatocytes of mice with hypomorphic expression of SPO11. Chromosoma **125**, 189–203, (2016).

5 Corbett, A. D. et al. Microscope calibration using laser written fluorescence. Opt. Express **26**, 21887–21899, (2018).

6 Garcés S, Y. et al. Nanoscale organization of rotavirus replication machineries. elife **8**, e42906, (2019).

7 Federici, F., Dupuy, L., Laplaze, L., Heisler, M., and Haseloff, J. Integrated genetic and computation methods for in planta cytometry, Nat. Methods **9**, 483–485 (2012).

8 Geldner, N., Dénervaud-Tendon, V., Hyman, D. L., Mayer, U., Stierhof, Y.-D. and Chory, J. Rapid, combinatorial analysis of membrane compartments in intact plants with a multicolor marker set, The Plant J. **59**, 169–178 (2009).

9 Pitrone, P., Schindelin, J., Stuyvenberg, L. et al. OpenSPIM: an open-access light-sheet microscopy platform, Nat Methods **10**, 598–599 (2013).

10 “Download micro-manager latest release:” https://micro-manager.org/Download_Micro-Manager_Latest_Release.

11 Rizzo, M. A., Davidson, M. W., & Piston, D. W. (2009). Fluorescent protein tracking and detection: fluorescent protein structure and color variants. Cold Spring Harbor Protocols, 2009(12), pdb-top63.

12 Huang, F., Hartwich, T., Rivera-Molina, F. *et al.* Video-rate nanoscopy using sCMOS camera–specific single-molecule localization algorithms. *Nat Methods* 10, 653–658 (2013).

13 Boškovi ́A., Bender, A., Gall, L., Ziegler-Birling, C., Beaujean, N., & Torres-Padilla, M.-E. Analysis of active chromatin modifications in early

mammalian embryos reveals uncoupling of h2a.z acetylation and h3k36 trimethylation from embryonic genome activation. Epigenetics **7**, 747–757 (2012).

14 Yahiatene, I., Hennig, S., Müller, M., & Huser, T. (2015). Entropy-Based Super-Resolution Imaging (ESI): From Disorder to Fine Detail. *ACS Photonics*, *2*(8), 1049–1056.

15 Agarwal, K., & Machá, R. (2016). Multiple signal classification algorithm for super-resolution fluorescence microscopy. *Nature Communications*, *7*.

16 Gustafsson, N., Culley, S., Ashdown, G. *et al.* Fast live-cell conventional fluorophore nanoscopy with ImageJ through super-resolution radial fluctuations. *Nat Commun* 7, 12471 (2016).

17 https://github.com/HenriquesLab/NanoJ-SRRF

18 Culley, S., Albrecht, D., Jacobs, C. *et al.* Quantitative mapping and minimization of super-resolution optical imaging artifacts. *Nat Methods* 15, 263–266 (2018).

19 Liu, S., Mlodzianoski, M., Hu, Z. *et al.* sCMOS noise-correction algorithm for microscopy images. *Nat Methods* 14, 760–761 (2017).

20 Mandracchia, B., Hua, X., Guo, C. *et al.* Fast and accurate sCMOS noise correction for fluorescence microscopy. *Nat Commun* 11, 94 (2020).

21 Descloux, A., Grußmayer, K.S. & Radenovic, A. Parameter-free image resolution estimation based on decorrelation analysis. *Nat Methods* 16, 918–924 (2019).

22 Tinevez, J. *et al.* TrackMate: An open and extensible platform for single-particle tracking. Methods 115 80–90 (2017).

23 Jaqaman, K. *et al.* Robust single-particle tracking in live-cell time-lapse sequences. Nat. Methods 5, 695–702 (2008).

24 Kalman, R. E. A new approach to linear filtering and prediction problems. J. Basic Eng. 82, 35–45 (1960).

25 Welch, G. and Bishop, G. An introduction to the kalman filter. Tech. Rep. 95-041, University of North Carolina at Chapel Hill, Chapel Hill, NC, USA (1995).

26 Crocker, J. C. and Grier, D. G. Methods of Digital Video Microscopy for Colloidal Studies. J. Colloid Interface Sci. 179, 298–310 (1996).

27 Mazzaferri, J. Roy, J. Lefrancois, S. and Costantino, S. Adaptive settings for the nearest-neighbor particle tracking algorithm. Bioinformatics 31, 1279–1285 (2014).
